# Analysis of paralogs in target enrichment data pinpoints multiple ancient polyploidy events in *Alchemilla* s.l. (Rosaceae)

**DOI:** 10.1101/2020.08.21.261925

**Authors:** Diego F. Morales-Briones, Berit Gehrke, Chien-Hsun Huang, Aaron Liston, Hong Ma, Hannah E. Marx, David C. Tank, Ya Yang

## Abstract

Target enrichment is becoming increasingly popular for phylogenomic studies. Although baits for enrichment are typically designed to target single-copy genes, paralogs are often recovered with increased sequencing depth, sometimes from a significant proportion of loci, especially in groups experiencing whole-genome duplication (WGD) events. Common approaches for processing paralogs in target enrichment datasets include random selection, manual pruning, and mainly, the removal of entire genes that show any evidence of paralogy. These approaches are prone to errors in orthology inference or removing large numbers of genes. By removing entire genes, valuable information that could be used to detect and place WGD events is discarded. Here we use an automated approach for orthology inference in a target enrichment dataset of 68 species of *Alchemilla* s.l. (Rosaceae), a widely distributed clade of plants primarily from temperate climate regions. Previous molecular phylogenetic studies and chromosome numbers both suggested ancient WGDs in the group. However, both the phylogenetic location and putative parental lineages of these WGD events remain unknown. By taking paralogs into consideration, we identified four nodes in the backbone of *Alchemilla* s.l. with an elevated proportion of gene duplication. Furthermore, using a gene-tree reconciliation approach we established the autopolyploid origin of the entire *Alchemilla* s.l. and the nested allopolyploid origin of four major clades within the group. Here we showed the utility of automated tree-based orthology inference methods, previously designed for genomic or transcriptomic datasets, to study complex scenarios of polyploidy and reticulate evolution from target enrichment datasets.

---

Polyploidy, or whole genome duplication (WGD), is prevalent throughout the evolutionary history of plants (Cui et al. 2006; Jiao et al. 2011; Jiao et al. 2012; Leebens-Mack et al. 2019). As a result, plant genomes often contain large numbers of paralogous genes from recurrent gene and genome duplication events (Lynch and Conery 2000; Panchy et al. 2016). Paralogs are defined as homologous genes that share a common ancestor as the product of gene duplication (Fitch 1970), either from small scale duplications or WGD. One special case of WGD is allopolyploidy, where genome doubling is accompanied by hybridization between two different species. The duplicated genes in allopolyploids are not paralogs in the traditional sense and are referred to as homoeologs, which are expected to be sister to the orthologous in the parental taxa, rather than to each other (Smedmark et al., 2003). For practical purposes, however, we refer to the product of any kind of duplications found in gene trees hereafter as paralog, as homoeologs are indistinguishable from paralogs until diagnosed as resulting from allopolyploidy. With very few nuclear genes being truly single- or low-copy, careful evaluation of orthology is critical for phylogenetic analyses (Fitch 1970). Orthology inference has received much attention in the phylogenomic era with multiple pipelines available for this task (e.g., Li et al. 2003; Dunn et al. 2013; Kocot et al. 2013; Yang and Smith 2014; Emms and Kelly 2019, also see Glover et. al 2019 and Fernández et al. 2020 for recent reviews). But these approaches have been mainly applied to genomic or transcriptomic data sets. So far, few studies have employed automated, phylogeny-aware orthology inference in target enrichment datasets. The most common approach for dealing with paralogy in target enrichment datasets is removing entire genes that show any evidence of potential paralogy (e.g., Nicholls et al. 2015; Jones et al. 2019; Andermann et al. 2020; but see Moore et al. 2018). Removal of entire genes might seem appropriate in target enrichment datasets in which only a small number of genes show evidence of paralogy (e.g., Larridon et al. 2020), but in some datasets this could result in a significant reduction of the number of loci (e.g., Montes et al. 2019). More importantly, dealing with paralogy only by removal of entire genes assumes that target enrichment assembly pipelines (e.g., Faircloth 2016; Johnson et al. 2016; Andermann et al. 2018), have flagged all genes with paralogs. It also assumes that if no sequence in a gene is flagged, all sequences are all single-copy and orthologous. On the other hand, this approach also removes genes that show allelic variation instead of paralogs. Given the prevalence of WGD and reticulations these assumptions can lead to errors in orthology inference. As paralogous genes are prevalent in plants, more appropriate orthology inference methods need to be applied in target enrichment data. The same automated approaches used for genome and transcriptome datasets can be applied for target enrichment, as these are tree-based and agnostic to the data source for tree inference.

The ability to explicitly process paralogs opens the door for using target enrichment data for inferring gene duplication events and pinpointing the phylogenetic locations of putative WGDs. In the past, the phylogenetic placement of WGD events have most often been carried out using genome and transcriptome sequencing data (e.g., Li et al. 2015; Huang et al. 2016; McKain et al. 2016; Yang et al. 2018) using either the synonymous distance between paralog gene pairs (Ks; Lynch and Conery 2000) or tree-based reconciliation methods (e.g., Jiao et al. 2011; Li et al. 2015; Yang et al. 2015; Huang et al. 2016; Xiang et al. 2017; Leebens-Mack et al. 2019). Similar to orthology inference, tree-based methods used to investigate WGDs in genome and transcriptome datasets should be useful in target enrichment data. Target enrichment methods (e.g., Mandel et al. 2014; Weitemier et al. 2014; Buddenhagen et al. 2016) have been widely adopted to collect hundreds to over a thousand nuclear loci for plant systematics, allowing studies at different evolutionary scales (e.g., Villaverde et al. 2018), and the use of museum-preserved collections (e.g., Forrest et al. 2019). This creates new opportunities to adopt tree-based reconciliation methods to explore WGD patterns in groups for which genomic and transcriptomic resources are not available or feasible.

With at least 350 (–1,100) species worldwide, *Alchemilla* in the broad sense has been a challenging group to study due to the presence of reticulate evolution, polyploidy, and apomixis. Based on previous phylogenetic analyses, *Alchemilla* s.l. contains four clades: Afromilla, *Aphanes,* Eualchemilla, and *Lachemilla* (Table S1). Together they form a well-supported clade nested in the subtribe Fragariinae, which also includes the cultivated strawberries (Gehrke et al. 2008). Unlike the more commonly recognized members of the rose family (Rosaceae), *Alchemilla* s.l. is characterized by small flowers with no petals, and a reduced number (1–4[–5]) of stamens that have anthers with one elliptic theca on the ventral side of the connective that opens by one transverse split (Perry 1929; Soják 2008). Gehrke et al. (2008) presented the first phylogeny of *Alchemilla* s.l. and established the paraphyly of traditional *Alchemilla* s.s. as consisted of a primarily African clade, Afromilla, and a Eurasian clade, Eualchemilla. Gehrke et al. (2008) also suggested treating Afromilla and Eualchemilla, along with *Aphanes* and *Lachemilla* as a single genus based on nomenclatural stability and the lack of morphological characters to distinguish between Afromilla and Eualchemilla. The four clades within *Alchemilla* s.l. are mainly defined by geographic distribution, as well as the number and insertion of the stamens on the disk lining the hypanthium (Table S1). Phylogenetic analyses using at least one nuclear and one chloroplast marker (Gehrke et al. 2008; 2016) found significant cytonuclear discordance regarding the relationships among the four major clades. Similar patterns, often attributed to hybridization and allopolyploidy, have been detected in other genera of Fragariinae (Lundberg 2009; Eriksson et al. 2015; Gehrke et al. 2016, Kamneva et al. 2017; Morales-Briones et al. 2018a), leaving the phylogenetic relationships of *Alchemilla* s.l. to the rest of Fragariinae unresolved. Unlike most members of Fragariinae that have predominantly diploid species, *Alchemilla* s.l. is known for high rates of polyploidy. The base chromosome number of *Alchemilla* s.l. is eight (*x* = 8), which differs from all other members in Fragariinae that have a base of number of seven (*x* = 7; Dickinson et al. 2007; Lundberg et al., 2009). Ploidy levels have been well documented in Eualchemilla that shows only polyploid species (2*n* = 64 to 220–224; octoploid to 28-ploid; e.g., Turesson 1943; Izmailow 1981; Walters and Bozman, 1967; Hayirhoğlu-Ayaz et al. 2006). *Aphanes* has mainly diploid species (2*n =* 16), with the exception of *Aphanes arvensis* that is an hexaploid (2*n =* 48; Montgomery et al. 1997). *Lachemilla* has mostly polyploid members (2*n* = 24 to 96; triploid to 12-ploid) with a single species reported to have diploid (2*n =* 16) and triploid (2*n =* 24) populations (Morales-Briones et al. 2018a). Lastly, little is known about ploidy levels in Afromilla, but so far, the two species reported were both polyploids (2*n* = 64 to 80; octoploid and decaploid; Hjelmqvist 1956; Morton 1993). A recent phylogenomic analysis focused on *Lachemilla* using target enrichment and 32 species of the group detected a high frequency of paralogs shared with Eualchemilla and Afromilla (Morales-Briones et al. 2018b). This paralog frequency suggested a possible ancient WGD event; however, the sampling was limited to one species each of Eualchemilla and Afromilla, and the location and mode of this putative WGD remained uncertain.

In this study we sampled 68 species across the major clades of *Alchemilla* s.l., and included 11 additional closely related species in Fragariinae, which allowed us to 1) test for polyploid events in the origin of *Alchemilla* s.l., and 2) explore the reticulate evolution among major clades of *Alchemilla* s.l. using a target enrichment dataset. Given the prevalence of polyploidy and reticulate history within *Alchemilla* s.l., this is an excellent group to explore the utility of tree-based methods for (1) processing paralogs, and (2) detecting and placing WGDs using target enrichment datasets.

## Materials and methods

### Taxon sampling and data collection

We sampled 68 species representing the four major clades of *Alchemilla* s.l. (sensu Gehrke et al. 2008), and 11 species to represent all other genera in Fragariinae (except *Chamaecallis*; sensu Dobeš et al. 2015; Morales-Briones and Tank 2019). Additionally, we sampled one species each of *Potentilla*, *Sanguisorba*, and *Rosa* as outgroups. Voucher information is provided in Table S2. We used a Hyb-Seq approach (Weitemier et al. 2014), that combines target enrichment and genome skimming, to capture nuclear exon sequences and off-target cpDNA. We used baits designed for *Fragaria vesca* (strawberry, also a member of Fragariinae) to target 1,419 exons in 257 genes (Kamneva et al. 2017). These genes were identified as single-copy orthologs among the apple (*Malus domestica*), peach (*Prunus persica*), and strawberry genomes based on reciprocal nucleotide similarity comparisons. The 257 genes resulted from first retaining only genes >960 bp long and with >85% similarity in pairwise comparisons among the three genomes. The remaining genes were further filtered by removing exons <80 bp long, with GC content <30% or >70%, and with >90% sequence similarity to annotated repetitive DNA in the genome, followed by removing exons with any paralogs with >90% sequence similarity in the same genome (Kamneva et al. 2017).

Of the 82 total species, only sequences for *Fragaria vesca*, were from a reference genome (Shulaev et. al 2010). Twenty-two were from a previously published Hyb-Seq dataset using the same bait set as this study (Morales-Briones et al. 2018b; Table S2), including 19 species of *Lachemilla* that did not show evidence of hybridization within *Lachemilla,* and one species each of Eualchemilla, Afromilla, and *Aphanes*. Newly generated sequence data for 55 species (Table S2) were collected as follows. Total genomic DNA was isolated from silica-dried or herbarium material with a modified CTAB method (Doyle and Doyle 1987). Probe synthesis, library preparation, capture enrichment, and high-throughput sequencing (HiSeq2000 instrument, 2 × 101 bp) were carried out at Rapid Genomics LLC (Gainesville, FL, USA). Data for the remaining four species, *Drymocallis glandulosa*, *Potentilla indica*, *Rosa woodsii*, and *Sanguisorba menziesii* were collected as described in Weitemier et al. (2014).

### Read processing and assembly

We removed sequencing adaptors and trimmed low-quality bases (Phred scores < 20) from raw reads with SeqyClean v.1.10.07 (Zhbannikov et al. 2017) using default settings. Plastomes were assembled using Alignreads v.2.5.2 (Straub et al. 2011) and 12 closely related plastome references (with one Inverted Repeat removed; Table S3). Plastome assemblies were annotated using *Fragaria vesca* as a reference in Geneious v.11.1.5 (Kearse et al. 2012). Assembly of nuclear loci was carried out with HybPiper v.1.3.1 (Johnson et al. 2016) using exons of *F. vesca* as references. Given the large number of paralogs detected in *Lachemilla,* Eualchemilla, and Afromilla, multi-exon gene assemblies resulted in chimeric sequences of exons from distinct paralogs (Morales-Briones et al. 2018b). To avoid chimeric sequences that can affect orthology inference and phylogenetic analyses, assemblies were performed on each exon separately. Only exons with a reference length of ≥ 150 bp were assembled (939 exons from 257 genes). Paralog detection was carried out for all exons with the ‘paralog_investigator’ option in HybPiper. This option flags loci with potential paralogs when multiple contigs cover at least 85% of the reference sequence length. Exon assemblies that included flagged paralogs were extracted using the ‘paralog_retriever’ command of HybPiper and used for orthology inference.

### Orthology inference for nuclear exons

To infer orthologs for phylogenetic analyses, all exons were processed as follows (Fig. 1a). Individual exons were aligned using MACSE v.2.03 (Ranwez et al. 2018) with default parameters. Codons with frameshifts (labeled with ‘!’ by MACSE) were replaced with gaps and aligned columns with more than 90% missing data were removed using Phyx (Brown et al. 2017). Initial homolog trees were built using RAxML v.8.2.11 (Stamatakis 2014) with a GTRCAT model and clade support assessed with 100 rapid bootstrap (BS) replicates. Clades and paraphyletic grades that belonged to the same taxon were pruned by keeping only the tip with the highest number of characters in the trimmed alignment following Yang and Smith (2014). To obtain the final homolog trees, outlier tips with unusually long branches were detected and removed by maximally reducing the tree diameter with TreeShrink v.1.3.2 (Mai and Mirarab 2018). Orthology inference was carried out using two outgroup-aware strategies from Yang and Smith (2014). We set *Potentilla*, *Sanguisorba*, and *Rosa* as outgroups and all members of Fragariinae as ingroups. First, we used the ‘monophyletic outgroup’ (MO) approach keeping only ortholog groups with at least 25 ingroup taxa. The MO approach filters for homolog trees with outgroup taxa being monophyletic and single-copy, and therefore filters for single- and low-copy genes. The second approach used was the ‘rooted ingroup’ (RT), with at least 25 ingroup taxa. The RT approach iteratively searches subtrees of ingroup taxa and cuts them out as rooted trees. Both approaches root the gene tree by the outgroups, traverse the rooted tree from root to tip, and remove the side with fewer taxa (MO) or keep both sides (RT) when gene duplication is detected at any given node. In the case of MO, homolog trees with non-monophyletic outgroups or duplicated taxa in the outgroups are discarded. If no taxon duplication is detected in a homolog tree, the MO approach outputs a one-to-one ortholog. The RT approach maximizes the number of orthologs compared to MO while not requiring monophyletic outgroups and allowing for duplicated taxa in the outgroups but removes outgroups from all orthologs. To add outgroups back to the RT orthologs for downstream analyses, we kept only RT orthologs from homologs that had a MO ortholog (i.e., using only homolog trees with monophyletic and non-duplicated outgroups for both MO and RT). Then we used the outgroups of the MO ortholog for all the RT orthologs of the same homolog (Fig. 1b). Scripts for orthology inference can be found at https://bitbucket.org/dfmoralesb/target_enrichment_orthology.

**Figure 1.**
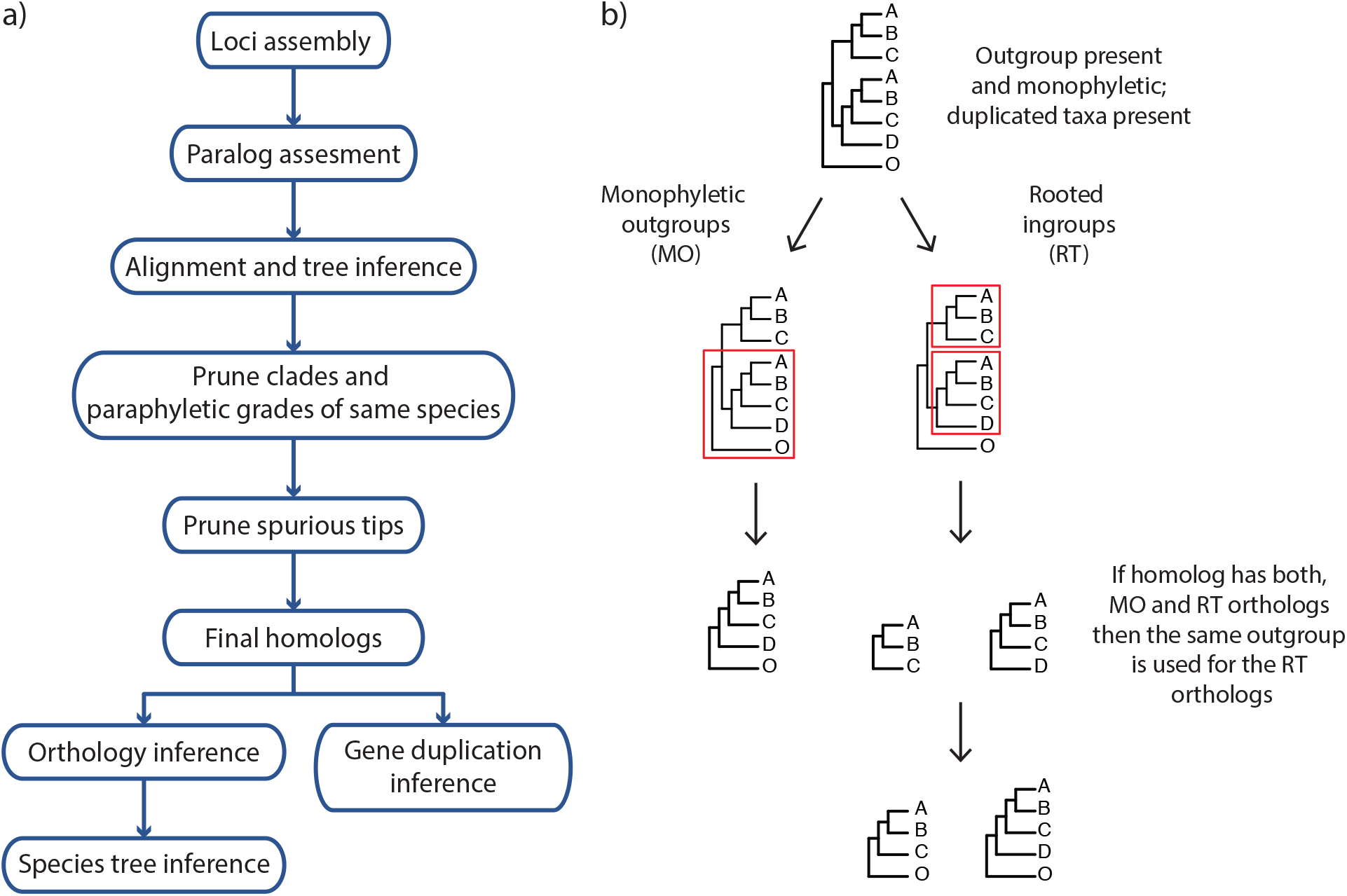
Paralog processing workflow and orthology inference methods used in *Alchemilla* s.l. homolog trees. a) Flow chart of paralog processing and homolog tree inference. b) Only homologs with outgroup present and monophyletic were used for orthology inference. Monophyletic outgroups (MO) will prune single-copy genes keeping clades with at least a user-defined minimum number of ingroup taxa. Rooted ingroups (RT) will keep all subtrees with at least a user-defined minimum number of ingroups taxa. If the homolog trees can be pruned using both MO and RT, then RT orthologs are added to the same root. Homologs that lack monophyletic outgroups were excluded from further consideration.

### Phylogenetic analyses

We used concatenation and coalescent-based methods to reconstruct the phylogeny of *Alchemilla* s.l. Analyses were carried out in the two sets of final orthologs, MO and RT, separately. Each ortholog was aligned using MACSE v.2.03 with default parameters. Codons with frameshifts were replaced with gaps, aligned columns with more than 90% missing data were removed using Phyx, and alignments with at least 150 characters and 25 taxa were retained. We first estimated a maximum likelihood (ML) tree from the concatenated matrices with RAxML using a partition by gene scheme with a GTRGAMMA model for each partition. Clade support was assessed with 100 rapid bootstrap (BS) replicates. To estimate a species tree that is statistically consistent with the multi-species coalescent (MSC), we first inferred individual ML gene trees using RAxML with a GTRGAMMA model, and 100 BS replicates to assess clade support. Individual gene trees were then used to estimate a species tree using ASTRAL-III v.5.6.3 (Zhang et al. 2018) using local posterior probabilities (LPP; Sayyari and Mirarab 2016) to assess clade support.

To evaluate nuclear gene tree discordance, we calculated the internode certainty all (ICA) value to quantify the degree of conflict on each node of the map tree (e.g., species tree) given individual gene trees (Salichos et al. 2014). Also, we calculated the number of conflicting and concordant bipartitions on each node of the map tree. We calculated both the ICA scores and the number of conflicting/concordant bipartitions with Phyparts (Smith et al. 2015) using individual ortholog trees with BS support of at least 50% for the corresponding node. Additionally, to distinguish strong conflict from weakly supported branches, we evaluated tree conflict and branch support with Quartet Sampling (QS; Pease et al. 2018) using 1,000 replicates. Quartet Sampling subsamples quartets from the input map tree (e.g., species tree) and concatenated alignment to assess the confidence, consistency, and informativeness of each internal branch by the relative frequency of the three possible quartet topologies at each node (Pease et al. 2018).

In addition to species tree construction using inferred orthologs, we used a recently developed quartet-based species tree method (ASTRAL-Pro; Zhang et al. 2020a) to estimate the phylogeny of *Alchemilla* s.l. ASTRAL-Pro directly uses multi-labeled gene trees while accounting for gene duplications and losses to estimate a species tree that is statistically consistent with the MSC and birth-death gene duplication and loss model. We used all 923 final homolog trees as input for ASTRAL-Pro, ignoring trees with less than 20 taxa, and estimated LPP to assess clade support. Additionally, we calculated ICA scores and the number of conflicting/concordant bipartitions with Phyparts using homolog trees with BS support of at least 50% for the corresponding nodes.

For the plastome phylogenetic analyses, 74 partial plastome assemblies and eight reference plastome sequences were included (Table S3). Contiguous plastome sequences were aligned using the default settings in MAFFT v.7.307 (Katoh and Standley 2013) and aligned columns with more than 70% missing data were removed with Phyx. We estimated an ML tree of the plastome alignment with RAxML using a partition by coding (CDS) and noncoding regions (introns and intergenic spacers) scheme, with a GTRGAMMA model for each partition and clade support assessed with 100 rapid BS replicates and QS using 1,000 replicates, to detect potential within-plastome conflict in the backbone of *Alchemilla* s.l. as recently reported in other groups (e.g., Gonçalves et al. 2019; Walker et al. 2019; Zhang et al. 2020b; Morales-Briones et al. 2021).

### Mapping whole genome duplications

We took two alternative approaches for detecting WGD events by mapping gene duplication events from gene trees onto a map tree (e.g., species tree). The first approach begins by extracting orthogroups from the final homolog trees. Orthogroups are rooted ingroup lineages separated by outgroups that include the complete set of genes in a lineage from a single copy in their common ancestor. We extracted orthogroups requiring at least 50 out of 79 species in Fragariinae. Gene duplication events were then recorded on the most recent common ancestor (MRCA) on the map tree when two or more species overlapped between the two daughter clades Each node on a map tree can be counted only once from each gene tree to avoid nested gene duplications inflating the number of recorded duplications (Yang et al. 2018; https://bitbucket.org/blackrim/clustering, ‘extract_clades.py’ and ‘map_dups_mrca.py’). We mapped duplication events onto both the MO and RT trees using orthogroups from all 923 final homologs, filtering orthogroups using an average BS of at least 50%. We carried out the mapping using two sets of orthogroups, one from all homologs, and the from the longest homologs (the single longest aligned exon per gene) to avoid inflating the counts in multi-exon genes.

For the second strategy of WGD mapping, we explicitly tested for polyploidy mode using GRAMPA (Thomas et al. 2017). GRAMPA uses MRCA reconciliation with multi-labeled gene trees to compare allo-or autopolyploid scenarios in singly- or multi-labeled map trees (e.g., species tree). To reduce the computational burden of searching all possible reconciliations, we constrained searches to only among crown nodes of major clades of *Alchemilla* s.l., which all are well supported (including the ‘dissected’ and ‘lobed’ clades of Eualchemilla; see results) and genera within Fragariinae. We ran reconciliation searches using all 923 final homologs, as well as using only the longest homologs (the single longest aligned exon per gene; 256), against either the MO or RT tree. We expected multiple WGD events within *Alchemilla* s.l. (see results), but GRAMPA can only infer one WGD at a time. To disentangle nested duplication events, we also carried out similar GRAMPA reconciliations using the MO tree and sequentially excluding major groups of *Alchemilla* s.l. that were identified as a polyploid clade. We only used the MO tree as it differs from the RT tree only by the location of the ‘lobed’ clade, which was the first clade identified as allopolyploid (see results) and was removed for subsequent GRAMPA analyses. Finally, to test for a polyploid origin of *Alchemilla* s.l., we carried out searches among the constrained crown node of *Alchemilla* s.l. and the rest of the genera within Fragariinae using the MO and cpDNA trees. The backbone of Fragariinae differed between the MO (same as RT) tree and the cpDNA tree. Thus, we tested how this affected the inference of the polyploid origin of *Alchemilla* s.l. We also carried out similar searches but using each of the five major clades individually.

Both approaches used here to detect WGD events use final homolog trees and as any other tree-based method they may be sensitive to tree informativeness. To explore node support across homologs, we run a conflict analysis with Phyparts using individual final homologs trees with a BS support filter of at least 50% for each node. We used both the MO and RT trees as map trees and ran the analysis using all homolog exons as well as only the longest homolog exon per gene.

### Distribution of synonymous distance among gene pairs (Ks plots)

To obtain further evidence for WGD events and compare them to those inferred from gene duplication events from target enrichment, we analyzed the distribution of synonymous distances (Ks) from RNA-seq data of four species of *Alchemilla* s.l. and nine species of Fragariinae (Table S4). Read processing and transcriptome assembly followed Morales-Briones et al. (2020). For each of the four species, a Ks plot of within-species paralog pairs based on BLASTP hits was done following Yang et al. (2018; https://bitbucket.org/blackrim/clustering; ‘ks_plots.py’). Ks peaks were identified using a mixture model as implemented in mixtools v.1.2.0 (Benaglia et al. 2009). The optimal number of mixing components was estimated using parametric bootstrap replicates of the likelihood ratio test statistic (McLachlan and Peel 2000). We tested up to five components using 500 bootstrap replicates in mixtools. Additionally, we used between-species Ks plots to determine the relative timing of the split between two species and compare it to that of WGD events inferred with within-species Ks plots. Ks plots of between-species also followed Yang et al. (2018; ‘ks_between_taxa_cds.py’). Lastly, we also attempted to build Ks plots using raw homologs from target enrichment, but the relatively low number of genes (256) failed to produce a meaningful distribution (not shown).

## Results

### Assembly and orthology inference

The number of assembled exons per species (with > 75% of the target length) ranged from 632 (*Alchemilla fissa*) to 934 (*Dasiphora fruticosa*) out of 939 single-copy exon references from *F. vesca*, with an average of 873 exons (Table S5). The number of exons with paralog warnings ranged from 10 in *Drymocallis glandulosa* to 746 in *Alchemilla mollis* (Table S5). The number of exon alignments with ≥ 25 species was 923 from 256 genes. The orthology inference resulted in 914 MO orthologs (Table S6), and 1,906 RT orthologs (Table S6). The trimmed alignments of the MO orthologs ranged from 136 to 5,740 characters with a mean of 425 characters (median = 268). The concatenated alignment of the MO orthologs, with at least 150 aligned characters and 25 species for each exon, included 910 exons and 387,042 characters with a matrix occupancy of 66%. The trimmed alignments of the RT orthologs ranged from 136 to 5,740 characters with a mean of 394 characters (median = 259). The concatenated alignment of the RT orthologs, with at least 150 aligned characters and 25 species, included 1,894 exons and 746,562 characters with a matrix occupancy of 54%. The chloroplast alignment included 124,079 characters with a matrix occupancy of 77%.

### Nuclear phylogenetic analyses

All nuclear analyses recovered the monophyly of *Alchemilla* s.l. with maximum support (i.e., bootstrap percentage [BS] = 100, local posterior probabilities [LPP] = 1.0; Fig. 2; Fig. S1), most informative gene trees being concordant with this node (858 out of 863 for MO; 977/984 for RT; 912/932 for ASTRAL-Pro; ICA = 0.95), and full QS support (1.0/–/1.0; i.e., all sampled quartets supported that node). Five major clades were identified within *Alchemilla* s.l.: Afromilla, *Aphanes*, Eualchemilla-‘dissected’, Eualchemilla-‘lobed,’ and *Lachemilla*. Moreover, the relationships among these clades showed high levels of discordance and varied among the MO and RT trees.

**Figure 2.**
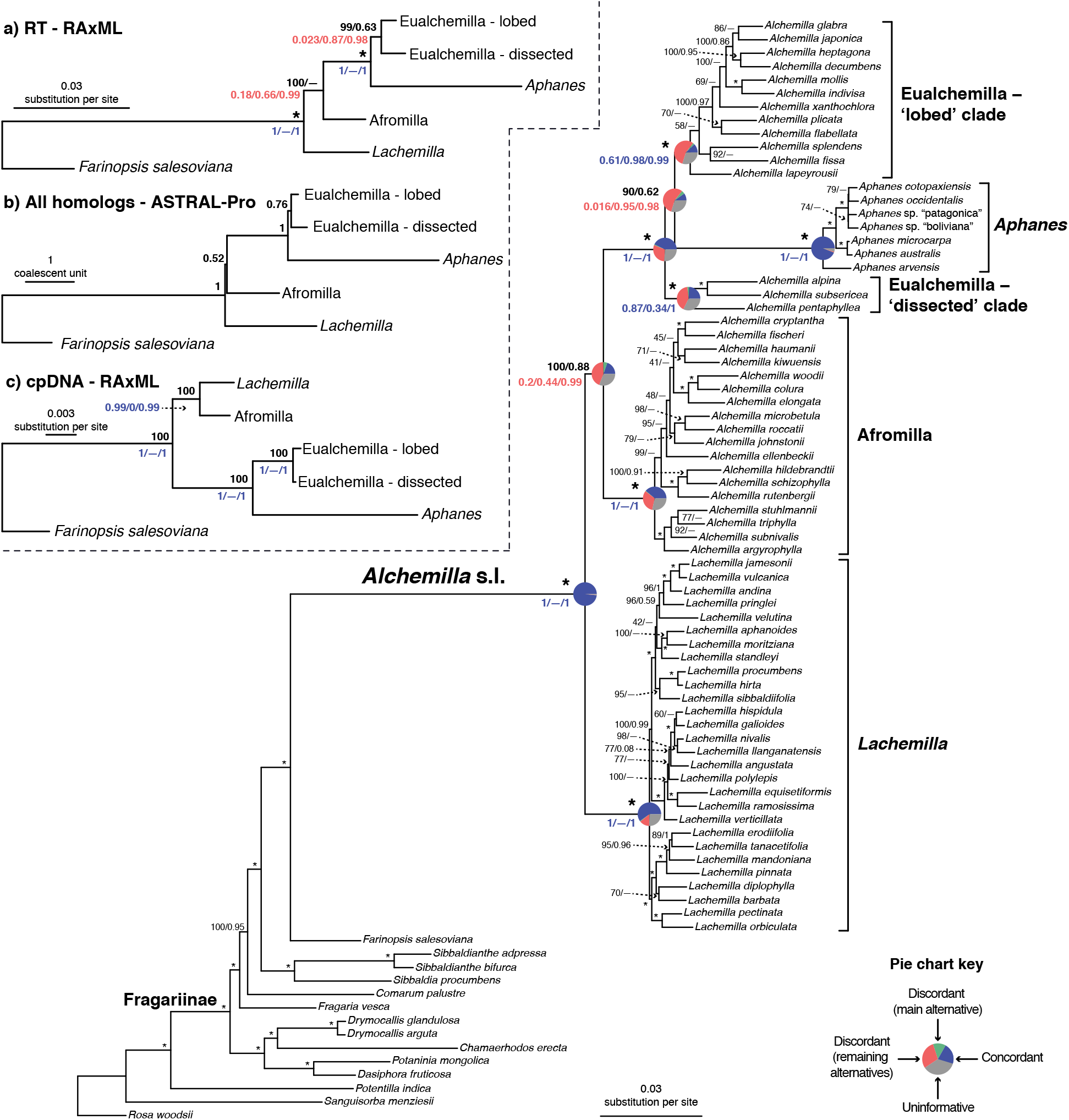
Maximum likelihood phylogeny of *Alchemilla* s.l. inferred from RAxML analysis of the concatenated 910-nuclear exon supermatrix from the ‘monophyletic outgroup’ (MO) orthologs. Bootstrap support (BS) and Local posterior probability (LLP) are shown above branches. Nodes with full support (BS= 100/LLP= 1) are noted with an asterisk (*). Em dashes (—) denoted alternative topology compared to the ASTRAL tree. Quartet Sampling (QS) scores for major clades are shown below branches. QS scores in blue indicate strong support and red scores indicate weak support. QS scores: Quartet concordance/Quartet differential/Quartet informativeness. Pie charts for major clades represent the proportion of exon ortholog trees that support that clade (blue), the proportion that support the main alternative bifurcation (green), the proportion that support the remaining alternatives (red), and the proportion (conflict or support) that have < 50% bootstrap support (gray). Gene trees with missing data that were uninformative for the node were ignored. Branch lengths are in number of substitutions per site (scale bar on the bottom). Inset: a) Summary Maximum likelihood phylogeny inferred from RAxML analysis of the concatenated 1,894-nuclear exon supermatrix from the ‘rooted ingroup’ orthologs (RT). BS and LLP are shown above branches and QS scores below the branches. Branch lengths are in number of substitutions per site; b) Summary ASTRAL-Pro tree inferred from 923 multi-labeled exon homolog trees. LLP are shown next to nodes. Branch lengths are in coalescent units. c) Summary Maximum likelihood phylogeny inferred from RAxML analysis of concatenated partial plastomes. BS and LLP are shown above branches and QS scores below the branches. Branch lengths are in number of substitutions per site.

Analyses of the MO orthologs using ASTRAL and concatenated ML approaches resulted in similar topologies for the backbone of *Alchemilla* s.l. (Fig. 2). The monophyly of the five major clades each received maximum support (BS = 100; LPP = 1.0) and had most trees being concordant (except for the two clades of Eualchemilla). Eualchemilla was paraphyletic and split into the ‘dissected’ and ‘lobed’ clades. Monophyly of the ‘dissected’ clade was supported by 118 out of 429 informative trees (ICA = 0.08) and strong QS score (0.87/0.34/1), while the ‘lobed’ clade was supported by 73 out of 420 informative trees (ICA = 0.06) and strong QS score (0.61/0.98/0.99). In both cases, the ‘dissected’ and ‘lobed’ clades had a relatively small percentage of supporting trees, but the conflict analysis and QS score did not reveal any well-supported alternative topology. *Aphanes* was recovered as sister to the Eualchemilla-‘lobed’ clade with relatively low support (BS = 90, LPP = 0.62), 60 concordant trees (out of 430 informative gene trees; ICA = 0.08), and weak QS score (0.016/0.95/0.98) with similar frequencies for the two discordant alternative topologies. The Eualchemilla-‘dissected’ clade was recovered as sister to Eualchemilla-‘lobed’ + *Aphanes* with maximum support, 279 concordant trees (out of 482 informative gene trees; ICA = 0.29), and full QS score. Afromilla was recovered as sister to the clade consisted of Eualchemilla (‘dissected and ‘lobed’) and *Aphanes* with high to low support (BS = 100, LPP = 0.88), only 146 concordant trees (out of 413 informative gene trees; ICA = 0.22), and weak QS support (0.2/0.44/0.99) with a skew in discordance suggesting a possible alternative topology (*Lachemilla* sister to Eualchemilla + *Aphanes*). Lastly, *Lachemilla* was recovered as the sister to the rest of *Alchemilla* s.l.

Analysis of the RT orthologs using ASTRAL and concatenated ML approaches both recovered the same major clades, but they differed in the relationship among these five clades (Fig. 2a; Fig. S1). In both analyses, *Lachemilla*, Afromilla, and *Aphanes* had maximum support (BS = 100; LPP = 1.0) and had most trees being concordant. Eualchemilla was recovered as monophyletic and composed of the ‘dissected’ and ‘lobed’ clades. The monophyly of Eualchemilla had high to low support (BS = 99, LPP = 0.63), only 231 concordant trees (out of 819 informative gene trees; ICA = 0.12), and weak QS support (0.023/0.87/0.98) with similar frequencies for the two discordant alternative topologies. Similar to the MO analyses, the ‘dissected’ and ‘lobed’ clades each had low number of concordant trees (218 out of 557 [ICA = 0.19] and 136 out of 707 [ICA = 0.08], respectively), and strong QS support (0.98/0/1 and 0.62/0.17/0.99, respectively). Eualchemilla was recovered as sister of *Aphanes* with maximum support (BS = 100), 348 concordant trees (out of 728 informative trees; ICA = 0.29) and full QS support. The ML concatenated tree (Fig. 2a; Fig. S1) placed Afromilla as sister to the clade formed of Eualchemilla and *Aphanes* with maximum support (BS = 100), 212 concordant gene trees (out of 771 informative trees; ICA = 0.27), and weak QS support (0.18/0.66/0.99) with no significant skew between the two discordant alternatives. *Lachemilla* was placed as sister to the rest of *Alchemilla* s.l. The ASTRAL tree in turn (Fig. S1a), retrieved *Lachemilla* as sister to the clade formed of Eualchemilla and *Aphanes* with no support (LPP = 0.01), 247 concordant trees (out of 953 informative trees; ICA = 0.19), and QS counter-support (−0.21/0.29/0.99), showing that the majority of the quartets supported one alternative topology (Afromilla sister to Eualchemilla + *Aphanes*). In this case, Afromilla was placed as sister to the rest of *Alchemilla* s.l.

The ASTRAL-Pro analysis using multi-labeled homolog trees recovered the same backbone topology of *Alchemilla* s.l. as the concatenated ML analysis from the RT orthologs (Fig. 2b; Figs S2–S3). All five major clades had the maximum support (LPP = 1.0). Eualchemilla, composed of the ‘dissected’ and ‘lobed’ clades, had moderate support (LPP = 0.76) and only 415 concordant trees (out of 1106 informative trees; ICA = 0.17). The ‘dissected’ and ‘lobed’ clades had low numbers of concordant trees (379 out of 941 [ICA = 0.23] and 65 out of 824 [ICA = 0.09], respectively), but did not show signal of any alternative topology. *Aphanes* was placed as the sister of Eualchemilla with maximum support (LPP = 1.0), 426 concordant trees (out of 952 trees; ICA = 0.34), and no support for any major alternative topology. Afromilla was recovered as sister to the clade formed of Eualchemilla and *Aphanes* with low support (LPP = 0.52), 492 concordant trees (out of 953 trees; ICA = 0.42), and no support for any alternative topology.

### Chloroplast phylogenetic analyses

The chloroplast ML tree (Fig. 2c; Fig. S4) recovered a well-supported backbone *Alchemilla* s.l. where the monophyly of *Aphanes,* Afromilla, and *Lachemilla,* had maximum or near maximum support (i.e., bootstrap percentage [BS] = 100, QS support = [1.0/–/1.0]). Eualchemilla, composed of the ‘dissected’ and ‘lobed’ clades, also had the maximum support. The ‘dissected’ and ‘lobed’ clades had strong support (BS = 75, QS = 0.8/0.43/0.88 and BS = 100, QS = 0.95/0.25/0.92, respectively). *Aphanes* and Eualchemilla formed, with maximum support, a clade as in the nuclear analyses. In turn, Afromilla and *Lachemilla* were recovered as sister clades with maximum support, which differed from the nuclear analyses.

### Mapping whole genome duplications

By mapping the most recent common ancestor (MRCA) of gene duplication events from orthogroup trees onto the MO and RT trees, we found four nodes in *Alchemilla* s.l. that each had an elevated proportion of gene duplications (Fig. 3a–b). This trend was consistent regardless of using all 923 homolog exons (868 after orthogroup inference and BS filtering) or using only the 256 longest homolog exons per gene (250 after orthogroup inference and BS filtering; Fig. S5). Therefore, here we describe the results only for the latter. These four clades include (Fig. 3a; Fig. S5): 1) the MRCA of *Alchemilla* s.l. (86.0% of the 250 genes show evidence of duplication), 2) the MRCA of Eualchemilla, *Aphanes*, and Afromilla (34.4%), 3) the MRCA of Eualchemilla + *Aphanes* (MO: 18.4%; RT:15.6%), and 4) the MRCA of *Lachemilla* (18.4%). These four nodes have an elevated proportion of gene duplications compared to all other nodes in Fragariinae (Fig. 3b) and it is consistent with the number of paralogs counted from the final homolog trees (after pruning of clades or paraphyletic grades of same species; Fig. 3c). Interestingly, although deeply nested in *Alchemilla* s.l., *Aphanes* had a lower number of paralogs than the rest of *Alchemilla* s.l., resembling the other members of Fragariinae (Fig. 3c).

**Figure 3.**
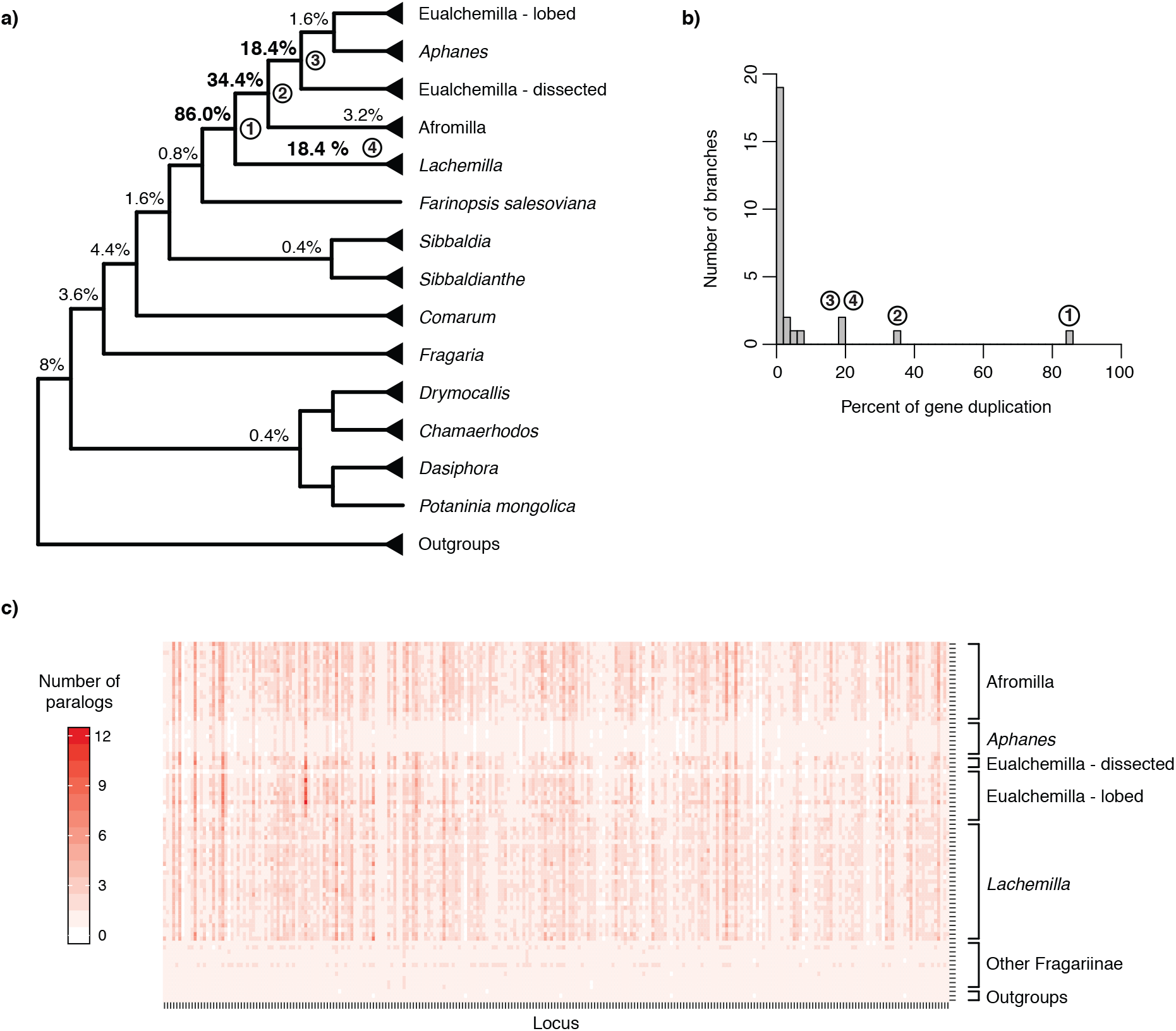
Orthogroup gene duplication mapping results. a) Summarized cladogram of *Alchemilla* s.l. from the ‘monophyletic outgroup’ (MO) ortholog tree. Percentages next to nodes denote the proportion of duplicated genes when using orthogroups from the longest homologs (250 after orthogroup inference and filtering). Nodes with elevated proportions of gene duplications are numbered 1–4 as referenced in the main text. See Fig. S5 for the full tree. b) Histogram of percentages of gene duplication per branch. c) Number of paralogs per taxa in the final homolog trees. In final homologs clades and paraphyletic grades of the same species were pruned leaving only one tip per species. Each locus is represented by the longest homolog (the single longest aligned exon per gene).

Bootstrap support for exon homologs were informative (BS ≥ 50%) at most nodes, especially regarding the relationship among the major clades of *Alchemilla* s.l. (Fig. S6). Therefore, uninformative homolog trees were unlikely to affect the results from WGD detection analysis overall. The proportion of uninformative nodes (BS < 50%) were at most 30% in the worst case (Eualchemilla + *Aphanes* + Afromilla) when using all homolog exons. This proportion reduces significantly when using only the longest homolog exons (Fig. S6).

Similar to the results of MRCA mapping, the GRAMPA analyses recovered the same results when using all 923 homologs or only the longest homologs (256). GRAMPA reconciliations using all major clades of *Alchemilla* s.l. recovered optimal multi-labeled trees with the best score (i.e., lowest reconciliation score; Fig. S7) where the ‘lobed’ clade of Eualchemilla was of an allopolyploid origin, but the putative parental lineages varied between the MO and RT trees. The reconciliations using the MO tree (reconciliation score [RS] = 70,250; Fig. 4a; Fig. S8) showed that the ‘lobed’ clade was of allopolyploid origin between an unsampled or extinct lineage sister to *Aphanes* and an unsampled or extinct lineage (‘lineage’ for short hereafter) sister to ‘dissected’ + *Aphanes*. In turn, the reconciliations using the RT tree (RS = 70,721; Fig. S8) showed that the ‘lobed’ clade was of allopolyploid origin between a ‘lineage’ sister to the ‘dissected’ clade, and also a ‘lineage’ sister to ‘dissected’ + *Aphanes*. Alternative multi-labeled trees had higher (worse) RSs (70,482 for MO and 70,739 for RT; Fig. S7). The GRAMPA reconciliations performed on the MO tree with removal of major clades of *Alchemilla* s.l. inferred as allopolyploid resulted in the identification of additional polyploidy events (Fig. 4b–d). First, we removed the ‘lobed’ clade, and this resulted in the recovery of Afromilla as an allopolyploid clade (RS = 127,836). Afromilla parental lineages were a ‘lineage’ sister to *Aphanes* + the ‘dissected’ clade, and a ‘lineage’ sister to all remaining *Alchemilla* s.l. (Fig. 4b). Alternative multi-labeled trees reconciliations had scores starting at 127,869 (Fig. S7). The further removal of Afromilla resulted in recovery of the ‘dissected’ clade as allopolyploid (RS = 167,545). The ‘dissected’ clade had as parental lineages the ‘lineage’ sister to *Aphanes* and the ‘lineage’ sister to all remaining *Alchemilla* s.l. except for *Lachemilla* (Fig. 4c). Other reconciliation alternatives had scores starting at 167,612 (Fig. S7). Finally, the removal of the ‘dissected’ clade resulted in the *Lachemilla* being recovered also as an allopolyploid clade (RS = 181,302). The parental lineages of *Lachemilla* were a ‘lineage’ sister to *Aphanes* and a ‘lineage’ sister to all remaining *Alchemilla* s.l. (Fig. 4d). Alternative multi-labeled trees reconciliations had scores starting at 181,564 (Fig. S7).

**Figure 4.**
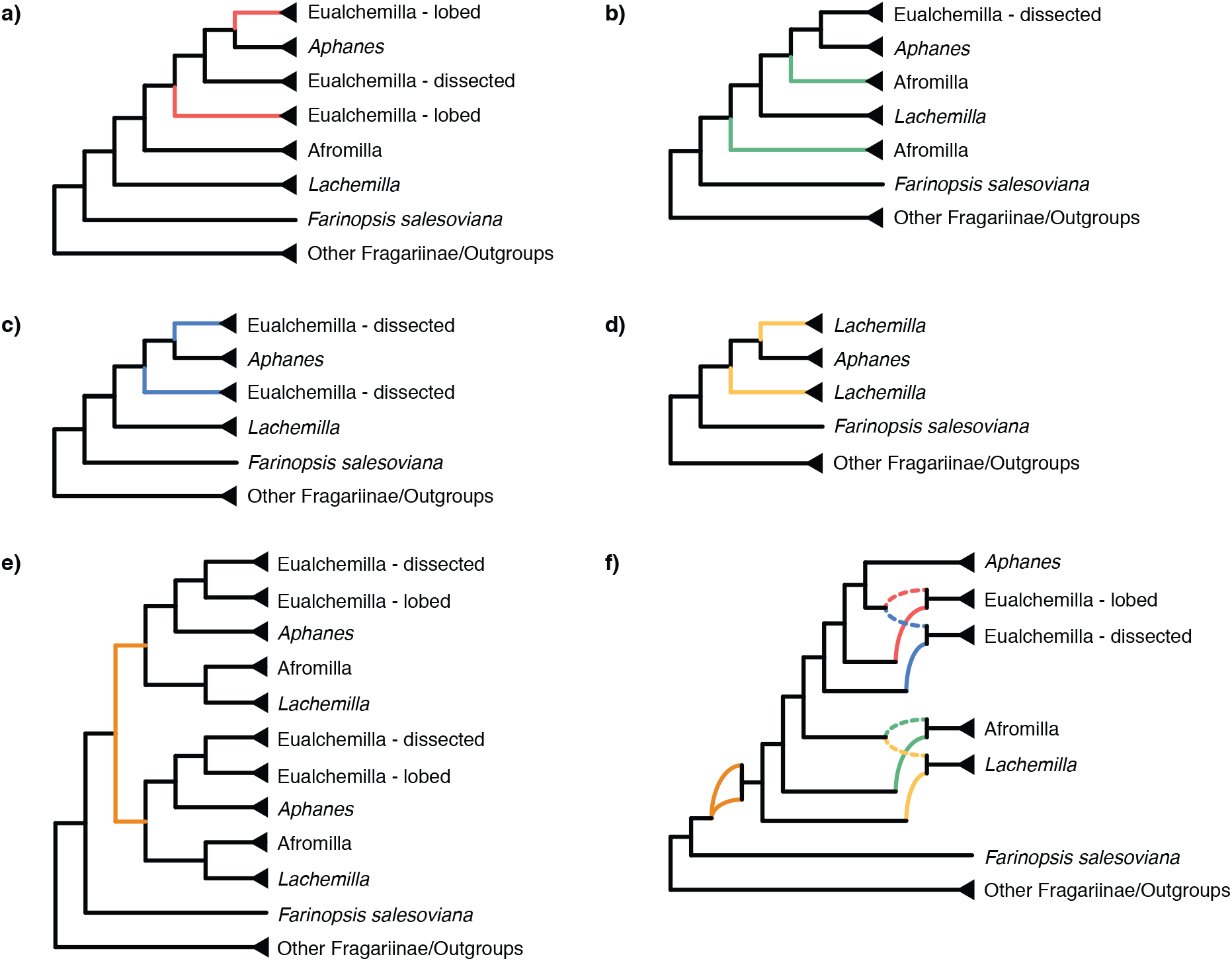
Summary of optimal multi-labeled tree (MUL-tree) inferred from GRAMPA analyses. a) MUL-tree based on reconciling homologs against the species tree inferred from ‘monophyletic outgroup’ (MO) orthologs including all taxa. Red branches denote the allopolyploid origin of the ‘lobed’ clade of Eualchemilla. b) MUL-tree after removing the ‘lobed’ clade of Eualchemilla as in a). Green branches denote the allopolyploid origin of Afromilla. c) MUL-tree after removing Afromilla as in b). Blue branches denote the allopolyploid origin of the ‘dissected’ clade of Eualchemilla. d). MUL-tree after further removing the ‘dissected’ clade as in c). Yellow lines denote the allopolyploid origin of *Lachemilla*. e) MUL-tree using constrained searches of the crown node of *Alchemilla* s.l. on the cpDNA tree. Orange branches denote the autopolyploid origin *Alchemilla* s.l. f) Putative summary network of all reticulation events in *Alchemilla* s.l. Colored curved branches denote different polyploid events as in a–e. Dashed curved lines represent the maternal lineage (cpDNA) in allopolyploid events.

The GRAMPA results from the analyses with constrained searches on the crown node of *Alchemilla* s.l. recovered different modes of polyploidy when using the MO tree or the cpDNA tree. The MO tree had *Farinopsis*, *Sibbaldianthe* + *Sibbaldia*, *Comarum*, and *Fragaria* forming a grade sister to *Alchemilla* s.l., while *Drymocallis*, *Chamaerhodos*, *Potaninia*, and *Dasiphora* form a clade that is sister to all other Fragariinae (Fig. 2). The reconciliations using the MO tree resulted in an allopolyploid event for the clade composed of *Alchemilla* s.l., *Farinopsis, Sibbaldianthe*, and *Sibbaldia* (RS = 339,755; Fig. S9). The parental lineages of this clade were a ‘lineage’ sister to *Comarum*, and a ‘lineage’ sister to the grade formed of *Comarum* and *Fragaria* (Fig. S9). Alternative multi-labeled trees had scores starting at 340,053 (Fig. S7). The reconciliations using individual major clades of *Alchemilla* s.l. resulted in identical patterns as in the full constrained analysis (Fig. S10). The cpDNA tree had *Alchemilla* s.l. as part of a grade formed along with *Farinopsis*, *Comarum*, and *Sibbaldianthe* + *Sibbaldia*, while *Fragaria* was recovered as sister to the clade composed of *Drymocallis*, *Chamaerhodos*, *Potaninia*, and *Dasiphora,* which is sister to all other Fragariinae (Fig. S4). The reconciliations on the cpDNA tree recovered *Alchemilla* s.l. as of autopolyploid origin (RS = 364,594; Fig. 4e; Fig. S9). Alternative multi-labeled trees had scores starting at 363,987 (Fig. S7). The analyses using individual major clades of *Alchemilla* s.l. recovered identical patterns as in the full constrained analysis, except for *Aphanes* that resulted in a singly-labeled tree (Fig. S10).

To further explore WGD events using alternative data sources, we analyzed Ks plots from genomes and transcriptomes across Fragariinae. The distribution of synonymous distances in the transcriptomes of four species of Eualchemilla (one ‘dissected’ and three ‘lobed’) shared three optimal mixing components with a Ks mean at approximately 0.1, 0.34, and 1.67, respectively (Fig. S11). The first two components partially overlapped and corresponded to at least two WGD events in all four sampled species of Eualchemilla, that happened before the splits between the lobed vs. the dissected clades of Eualchemilla (Ks ∼ 0.02; Fig. S12). The third shared component corresponds to a whole genome triplication event early in the core eudicots (Jiao et al. 2012; Fig. S11). All nine species from other genera in Fragariinae had two optimal mixing components. One component is a Ks peak at 1.61–1.78 corresponding to the whole genome triplication event early in eudicot (Fig. S11). In the case of the diploid species, the second component represents a small and very young (∼ 0.05) peak, most likely the product of small-scale recent gene duplications. The only two polyploid species from the other genera in Fragariinae, *Comarum palustre* (2n=28–64) and *Sibbaldianthe bifurca* (2n=28), had a single additional significant component at 0.11 and 0.08, respectively (Fig. S11). The Ks plots between species of Eualchemilla and Fragariinae species outside of *Alchemilla* s.l., and between species of Fragariinae showed that the WGD events detected in Eualchemilla were not shared with other genera outside of *Alchemilla* s.l. Likewise, the WGD events in *Comarum palustre* and *Sibbaldianthe bifurca* occurred after the split of the two species (Fig. S12).

## Discussion

### Processing paralogs in target enrichment datasets

The increased use of target enrichment methods in combination with reduced sequencing costs and higher read coverage have facilitated the recovery of paralogs in such datasets. Paralogy is sometimes viewed as a nuisance for phylogenetic reconstruction and is commonly aimed to be reduced in early stages of experimental design, by targeting only single- or low-copy genes during the selection of loci (e.g., Chamala et al. 2015; Nicholls et al. 2015; Gardner et al. 2016; Kamneva et al. 2017). Still, the recovery of paralogs is inevitable when working with groups where WGD is prevalent, especially in plants, leading to various strategies to remove them prior to phylogenetic analyses. Commonly used target enrichment assembly pipelines (e.g., Faircloth 2016; Johnson et al. 2016; Andermann et al. 2018) use different criteria to flag assembled loci with putative paralogs that are later filtered or processed prior to phylogenetic analysis. The most used common approach for dealing with paralogous loci in target enrichment datasets is removing the entire locus that show any signal of potential paralogy (e.g., Crowl et al. 2017; Montes et al. 2019; Bagley et al. 2020). The removal of paralogous loci can significantly reduce the size of target enrichment datasets and most often do not take in consideration the reason why a locus was flagged for putative paralogy (i.e., allelic variation or gene duplication). Orthology inference should be carried for all loci in target enrichment data, as relying on settings in assembly pipelines does not guarantee that non-removed or non-flagged loci are orthologous. Furthermore, removing paralogs before phylogenetic inference eliminates valuable information that could have been used to detect and place WGD events using target enrichment data. Other approaches either retain or remove contigs based on the distinction being putative allelic variation (flagged sequences from monophyletic conspecific groups) or putative paralogs (paralogs from the same species are non-monophyletic) in combination with study-specific threshold or random selection (e.g., Villaverde et al; 2018; Liu et al. 2019; Stubbs et al. 2019), or manual processing (e.g., Garcia et al. 2017; Karimi et al. 2019). As dataset size increases, manual processing becomes prohibitive.

The presence of WGDs also poses some challenges for locus assembly. Target enrichment design commonly includes multi-contig targets that assembly pipelines attempt to assemble into single contigs (e.g., Faircloth 2016) or ‘supercontigs’ composed of multiple exons and partially assembled introns (e.g., Johnson et al. 2016). In groups like *Alchemilla* s.l., where multiple, nested WGD events led to a prevalence of paralogs, ‘supercontigs’ can produce chimeric assemblies (Morales-Briones et al. 2018b). Instead, we assembled the exons individually to minimize chimeric loci, at the cost of working with some short exons that contribute little phylogenetic information, which can affect orthology inference and downstream analyses. Therefore, it is important to take this into consideration during target enrichment experimental design, and to preferentially target long exons when possible in groups where WGD is expected. An alternative strategy to avoid chimeric supercontigs when gene duplications are frequent is to perform a preliminary orthology inference in single exon-based trees and then use the inferred orthologs as a reference to reassemble the loci into ‘supercontigs’ (e.g., Gardner et al. 2020; Karimi et al. 2020). Another aspect to take in consideration during or right after assembly is allele phasing. While phasing heterozygous loci, from population or individual variation, have been shown to have minimal impact in phylogenetic reconstruction in target enrichment data (e.g., Kates et al. 2018), the effect on unphased or merged loci in cases of WGD can be larger and be a source of gene tree error. Here we were interested in ancient WGD in *Alchemilla* s.l. and relied on enough sequencing coverage and sequence dissimilarity to assemble separate paralogs (homoeologs in the case of allopolyploidy) that can be flagged as such by HybPiper. While we obtained a large number of deep paralogs across *Alchemilla* s.l. (Fig. 3c; Table S5), there is still the possibility of some locus included merged sequences from paralogs with high sequence similarity. Paralog merger should be more problematic in cases of recent allopolyploidy or neo-allopolyploidy taxa. To this end, recently developed tools have been designed to phase gene copies into polyploid subgenomes using phylogenetic and similarity approaches (e.g., Freyman et al. 2020, Nauheimer et al. 2020).

The utility of paralogs for phylogenetic reconstruction in target enrichment datasets is gaining more attention (e.g., Johnson et al. 2016; Gardner et al. 2020). A few studies have considered tree-based orthology inference to process affected loci (e.g., Garcia et al. 2017; Moore et al. 2018, Morales-Briones et al. 2018b), but in some cases the orthology approaches used cannot be applied to other groups. Here we demonstrated the utility of automated, tree-based orthology inference methods (Yang and Smith 2014), originally designed for genomic or transcriptomic datasets, to infer orthology from paralog-flagged loci in a target enrichment dataset. Our approach facilitates the automated inference of orthologs while maximizing the number of loci retained for downstream analyses. These methods are agnostic of the data source and should work for any type of target enrichment dataset (e.g., anchored phylogenomics, exon capture; Hyb-Seq, ultraconserved elements).

Orthology inference methods used here (Yang and Smith et al. 2014) are a powerful tool for target enrichment datasets. In the case of allopolyploidy, however, these methods can introduce bias in the distribution of ortholog trees inferred. In the case of MO, each time a gene duplication event is detected, the side with a smaller number of taxa is removed. When allopolyploidy occurs, MO may bias towards one subgenome due to 1) bias in gene loss between subgenomes. Even if the submissive subgenome is present in some parts of the genome, it is differentially lost in a higher number of loci, 2) bias in bait design. In the case of *Alchemilla* s.l., this is less likely as baits are designed in outgroups. If baits are designed according to ingroup taxa, depending on which taxa were used it can have higher affinity to one subgenome instead of another, and 3) unequal sampling of parental lineages. If one parental lineage is more densely sampled than the other or one parental lineage is unsampled, the two subgenomes will be in species-rich versus species-poor clades respectively in gene trees. One could alternatively preserve a random side each time a gene duplication event is identified. However, in practice, the side with a smaller number of taxa often contains misassembled or misplaced sequences. The RT method of separating duplicated gene copies, on the other hand, keeps any subtree with sufficient number of taxa, but removes outgroups, and worked best when hierarchical outgroups were included in the taxon sampling. Therefore, both MO and RT lose information, especially in cases with complex, nested polyploidy. Recently developed methods based on quartet similarity (Zhang et al. 2020a) or Robinson-Foulds distances (Molloy and Warnow 2020) can directly estimate species trees from multi-labeled trees that are consistent with the MSC and gene duplication and loss without inferring orthologs (for a recent review see Smith and Hahn 2020). However, their behavior on complex datasets using archival materials is yet to be explored. For example, both methods do not define ingroup-outgroup relationships a priori, and correctly inferring the root of homolog trees can be challenging with missing data, or when WGD occurs near the root. In addition, none of these above species tree reconstruction methods (Molloy and Warnow 2020; Zhang et al. 2020a) were designed to handle reticulate relationships. This can result in species tree topology that is an “average” between subgenomes. Depending on the topological distance of subgenomes, the resulting species tree may not represent any subgenome history. Finally, most current methods for evaluating node support still require orthologous gene trees as input. In such cases tools like Phyparts can still be used to visualize gene tree discordance and calculate ICA scores using multi-labeled trees.

### Phylogenetic implications in Alchemilla s.l

Previous phylogenetic studies established the monophyly *Alchemilla* s.l. and four major clades of the group (Gehrke et al. 2008; 2016), but the relationship among them and the placement of *Alchemilla* s.l. within Fragariinae remain unresolved. Our nuclear and plastid analyses both confirmed the monophyly of *Alchemilla* s.l. and its sister relationship to *Farinopsis*, as previously shown by Morales-Briones and Tank (2019) based on plastome sequences only. Gehrke et al. (2008) identified two well supported clades within Eualchemilla that were distinguished by leaf shape, namely the ‘dissected’ and ‘lobed’ clades. Most species of Eualchemilla have a leaf shape consistent with their clade placement, but some had different leaf shapes that were attributed to their hybrid origin between the two clades (Gehrke et al. 2008). More recently, Gehrke et al. (2016) and Morales-Briones and Tank (2019) found that Eualchemilla is not monophyletic in analyses that included the external transcribed spacer (ETS) of the nuclear ribosomal DNA (nrDNA) cistron. Both studies found *Aphanes* nested between the ‘dissected’ and ‘lobed’ clades of Eualchemilla. Our analyses of the nuclear loci supported the monophyly of ‘dissected’ and ‘lobed’ clades, but the monophyly of Eualchemilla had low support (Fig. 2; Fig. S1). The analysis using only the MO orthologs even weakly supported the ‘lobed’ clade as sister of *Aphanes* (Fig. 2). In contrast, our plastome analysis recovered a well-supported, monophyletic Eualchemilla, as well as well-supported ‘dissected’ and ‘lobed’ clades. Both nuclear and plastid analyses strongly supported the clade composed of *Aphanes* and both clades of Eualchemilla (Fig. 2; Fig. S4), a relationship that is consistent with previous nuclear and plastid analyses (Gehrke et al. 2008; 2016; Morales-Briones and Tank 2019). Given the revealed hybridization in the evolution and early divergence within *Alchemilla* s.l., the non-monophyly of Eualchemilla could be explained by ancient gene flow or an allopolyploid origin of the ‘dissected’ and ‘lobed’ clades (Fig. 4a,c; see below). Besides the well supported relationship of Eualchemilla + *Aphanes*, our nuclear analysis showed high levels of conflict among other major clades in *Alchemilla* s.l. (Fig. 2; Fig. S1) which could also be explained by additional ancient allopolyploid events (Fig. 4; see below).

### Ancient polyploidy in Alchemilla s.l

Whole-genome duplications are frequent across Rosaceae (Dickinson et al. 2007; Xiang et al. 2017), and allopolyploidy has been suggested as the primary source for the cytonuclear discordance in Fragariinae (Lundberg et al. 2009; Gehrke et al. 2016; Morales-Briones and Tank 2019). We recovered four nodes in *Alchemilla* s.l. with a high percentage of gene duplications (Fig. 3a; Fig. S5). One of the nodes showing a high percentage of gene duplication (18.4%) was the MRCA of *Aphanes* and both clades of Eualchemilla (node 3 in Fig. 3a; Fig. S5). This duplication event agreed with the MRCA of the ancestral lineages inferred with GRAMPA for the allopolyploid origin of the ‘lobed’ clade of Eualchemilla (Fig. 4a). Moreover, the GRAMPA reconciliations after the removal of the ‘lobed’ clade and Afromilla inferred a scenario where the ‘dissected’ clade is of allopolyploid origin with one of the parental lineages as sister to *Aphanes* (Fig. 4c). Although there is some uncertainty about the placement of the parental lineage of ‘dissected’ clade, due to the removal of major clades for the GRAMPA analyses, the cpDNA tree suggest that it is likely sister to the parental lineage of ‘lobed’ clade that is also sister to *Aphanes*. Ks plots of all species of the ‘dissected’ and ‘lobed’ clades had two peaks that are not shared with members of Fragariinae (Figs S11–S12), suggesting that at least two WGD events have happened between the stem lineage of *Alchemilla* s.l. to the crown node of the ‘dissected’ and the ‘lobed’ clades. The between-species Ks plots between ‘dissected’ and ‘lobed’ (Fig. S12), showed that the split between these two groups is more recent than the WGD events, suggesting a single origin (or very close in time) of both clades. Still, the sister relationship of the ‘dissected’ and ‘lobed’ clades is not supported by nuclear genes, suggesting that the two clades of Eualchemilla might had independent allopolyploid origins, while sharing the same or a closely related maternal lineage (cpDNA; Fig. 4f).

The GRAMPA reconciliation, after the removal of the ‘lobed’ clade, recovered an allopolyploid origin of Afromilla (Fig. 4b) with a MRCA of the ancestral lineages at the crown of the remaining *Alchemilla* s.l. Similarly, the further removal of both Afromilla and the ‘dissected’ clade recovered *Lachemilla* as allopolyploid, with the MRCA of parental lineages mapped to the crown of the remaining *Alchemilla* s.l. (Fig. 4d). In the case of *Lachemilla,* because of the removal of major clades for the GRAMPA analyses, there is also no certainty in the placement of its parental lineages. Still, Afromilla and *Lachemilla* are sisters in the cpDNA tree (Fig 1C.), suggesting these two share the same or a closely related maternal lineage (Fig. 4f). The high percentage of gene duplication (34.4%) placed at the MRCA of the clade composed of Afromilla, Eualchemilla, and *Aphanes* (node 2 in Fig. 3a), could be explained in part by the allopolyploid origin of Afromilla.

Finally, the node with the highest percentage of duplicated genes (86%) was placed at the MRCA of *Alchemilla* s.l. (node 1 in Fig. 3a). The GRAMPA analysis using the MO tree showed an allopolyploid event for the clade that included *Alchemilla* s.l., *Farinopsis salesoviana*, *Sibbaldia*, and *Sibbaldianthe* (Fig. S9). However, an allopolyploid origin of *Farinopsis salesoviana*, *Sibbaldia*, and *Sibbaldianthe* is not supported by chromosome numbers, orthogroup gene duplication counts, or Ks plots. All members in Fragariinae, with the exception of *Alchemilla* s.l. mainly consists of diploid species and base chromosome number of seven (*x* = 7), including *Sibbaldia* and *Sibbaldianthe.* On the other hand, *Alchemilla* s.l. has a base number of eight (*x* = 8) and contains mostly species with high ploidy levels (octoploid to 24-ploid), with the exception of most species of *Aphanes* (2*n*=16) and one species of *Lachemilla* (*L. mandoniana,* 2*n*=16). Also, our gene duplication counts show low percentages (1.6%) of gene duplication for the MRCA of the GRAMPA-inferred allopolyploid clade or the MRCA (3.6%) of the inferred parental lineages (Fig. 3). Previous phylotranscriptomic analyses of Rosaceae (Xiang et al. 2017) that included one species each of the ‘dissected’ and ‘lobed’ clades of Eualchemilla, found 33.21% of duplicated genes for the MRCA of these two clades, but did not recover any other node with elevated gene duplications within Fragariinae. The Ks plots of the four species of *Alchemilla* s.l. all showed peaks with similar Ks means, but these peaks were not shared with species of *Sibbaldia* and *Sibbadianthe* (Fig. S11). Furthermore, the between-species Ks plots showed that the WGD events detected in *Alchemilla* were more recent than the split with members of Fragariinae (Fig. S12). Although the chromosome number and Ks data for *Farinopsis salesoviana* are not available, all the above evidence suggest an unlikely allopolyploid origin of the clade consisting of *Farinopsis, Sibbaldia*, *Sibadianthe*, and *Alchemilla* sl. On the other hand, the GRAMPA reconciliations using the cpDNA tree resulted in an optimal multi-labeled tree where *Alchemilla* s.l. had an autopolyploid origin (Fig. 4e). This scenario is compatible with the high percentage of gene duplication at the MRCA of *Alchemilla* s.l. and the low percentage of gene duplication in the backbone of the rest of Fragariinae. Another compatible scenario is an allopolyploid origin of *Alchemilla* s.l. where both parental lineages are missing or extinct, but this scenario is indistinguishable from autopolyploidy. The atypical high proportion of gene duplication at the base of *Alchemilla* s.l. can be explained by the autopolyploid event at this branch. In addition, given the short branch lengths among major clades within *Alchemilla* s.l., gene tree estimation error (e.g., uninformative genes), incomplete lineage sorting (ILS), allopolyploid events among major clades of *Alchemilla* s.l., and/or homoeologous exchanges among subgenomes (Edger et al. 2018; McKain et al. 2018) can all contribute to additional gene duplication events mapped to the MRCA of *Alchemilla* s.l.

Although deeply nested in *Alchemilla* s.l., remarkably, *Aphanes* showed a significantly lower number of paralogs than the rest of *Alchemilla* s.l. (Fig. 3). The relatively low number of paralogs, its diploid species being mainly diploid, and the best GRAMPA reconciliation resulting in a singly-labeled tree (Fig. S10), suggesting that *Aphanes* is a functional diploid clade. One plausible scenario is that post-polyploid diploidization (reviewed in Mandáková and Lysak 2018) occurred after the autopolyploidy event at the base of *Alchemilla* s.l. After diploidization, Afromilla, Eualchemilla (‘lobed’ and ‘dissected’ clades), and *Lachemilla* originated from allopolyploid events (Fig. 4f). On the other hand, *Aphanes* seems to descend from a diploidized ancestor that did not duplicate further. The orthogroup gene duplication mapping showed *Aphanes* as part of a clade that had nested high proportions of gene duplication in the orthogroup mapping (Fig. 3a-b, nodes 2–3). But this does not necessarily mean that *Aphanes* should show the same duplication pattern, or neither does it contradict its diploid condition, as a duplication event does not affect or include all descendants of the mapped MRCA in the map tree (e.g., species trees).

GRAMPA has been shown to be useful to identify multiple polyploidy events in the same tree (e.g., Thomas et al. 2017; Guo et al. 2020; Koenen et al. 2020), but a tree-based approach can also be sensitive to gene tree estimation error or ILS (Thomas et al. 2017). Methods to infer species networks in the presence of ILS (e.g., Solís-Lemus and Ané 2016; Wen et al. 2018) could also be used to explore the prevalence of ancient hybridization in *Alchemilla* s.l. Although these methods are under constant development and improvement, they are still only tractable in simple scenarios with few reticulation events (Hejase and Liu 2016; Kamneva and Rosenberg 2017). Similarly, the signal of the *D*-Statistic (Green et al. 2010; Durand et al. 2011), commonly used to detect introgression, can be lost or distorted in presence of multiple reticulations (Elworth et al. 2018). Complex reticulate scenarios like *Alchemilla* s.l. are likely to face these problems and have phylogenetic network and *D*-statistic identifiability issues as seen in other groups (e.g., Morales-Briones et al. 2021).

### Conclusions

In this study, we have shown the utility of target enrichment datasets in combination with tree-based methods for orthology inference and WGD investigation. Here, we used *Alchemilla* s.l. to highlight the importance of processing paralogs, rather than discarding them before phylogenetic analysis, to shed light on the complex polyploidy histories. We showed evidence that the entire *Alchemilla* s.l. is the product of an ancient autopolyploidy event, and that Afromilla, Eualchemilla (‘lobed’ and ‘dissected’ clades), and *Lachemilla* originated from subsequent and nested ancient allopolyploid events. Our results from analyzing target enrichment data corroborated with previously published chromosome numbers and distribution of Ks values from transcriptomes. Our analyses has several important implications for future target enrichment projects, including 1) design baits to obtain a relatively large number of loci as this is required for accurate species tree and networks estimation in complex scenarios (e.g., higher levels of ILS; Solís-Lemus and Ané 2016; Nute et al. 2018), 2) increase the length of individual loci to improve the information content of individual gene trees for proper tree-based orthology inference and identifying gene duplication events, and 3) design baits to minimize lineage-specific and paralog-specific capture efficiency and missing data. Furthermore, in target enrichment, unlike genome or transcriptome data, only a few hundreds of genes are typically recovered with levels of missing data that varies by lineage and are non-random. This limits the utility of target enrichment for generating Ks plots, and creates the need to carefully scrutinize the variation in percentage of gene duplications among nodes. In the end, even with these limitations, target enrichment is an overall valuable and cost-effective approach of genomic subsampling to explore patterns of reticulation and WGD, especially in groups for which whole genome or transcriptome data are not possible to generate, including from museum/herbarium specimens. As research continues to deepen in other clades across the Tree of Life using similar target enrichment methods, we expect that other complex patterns of duplication and reticulation, as those shown here in *Alchemilla* s.l. will continue to emerge.

## Supplementary material

Data available from the Dryad Digital Repository: http://dx.doi.org/10.5061/.[NNNN]

## Acknowledgments

This work was supported by the University of Minnesota, Graduate Student Research Grants from the Botanical Society of America, American Society of Plant Taxonomists, International Association of Plant Taxonomists, the University of Idaho Stillinger Herbarium Expedition Funds to D.F.M.-B., and a National Science Foundation Doctoral Dissertation Improvement Grant to D.C.T. for D.F.M.-B. (DEB-1502049).

## References

Andermann T., Cano Á., Zizka A., Bacon C., Antonelli A. 2018. SECAPR—a bioinformatics pipeline for the rapid and user-friendly processing of targeted enriched Illumina sequences, from raw reads to alignments. PeerJ. 6:e5175.

Andermann T., Torres Jiménez M.F., Matos-Maraví P., Batista R., Blanco-Pastor J.L., Gustafsson A.L.S., Kistler L., Liberal I.M., Oxelman B., Bacon C.D., Antonelli A. 2020. A Guide to Carrying Out a Phylogenomic Target Sequence Capture Project. Front. Genet. 10:1407.

Bagley J.C., Uribe-Convers S., Carlsen M.M., Muchhala N. 2020. Utility of targeted sequence capture for phylogenomics in rapid, recent angiosperm radiations: Neotropical Burmeistera bellflowers as a case study. Mol. Phylogenet. Evol.:106769.

Benaglia T., Chauveau D., Hunter D.R., Young D. 2009. mixtools : An R Package for Analyzing Finite Mixture Models. J. Stat. Softw. 32:1–29.

Brown J.W., Walker J.F., Smith S.A. 2017. Phyx - phylogenetic tools for unix. Bioinformatics. 33:1886–1888.

Buddenhagen C., Lemmon A.R., Lemmon E.M., Bruhl J., Cappa J., Clement W.L., Donoghue M.J., Edwards E.J., Hipp A.L., Kortyna M., Mitchell N., Moore A., Prychid C.J., Segovia-Salcedo M.C., Simmons M.P., Soltis P.S., Wanke S., Mast A. 2016. Anchored Phylogenomics of Angiosperms I: Assessing the Robustness of Phylogenetic Estimates. bioRxiv.:086298.

Chamala S., García N., Godden G.T., Krishnakumar V., Jordon-Thaden I.E., De Smet R., Barbazuk W.B., Soltis D.E., Soltis P.S. 2015. MarkerMiner 1.0: A new application for phylogenetic marker development using angiosperm transcriptomes. Appl. Plant Sci. 3:1400115.

Crowl A.A., Manos P.S., McVay J.D., Lemmon A.R., Lemmon E.M., Hipp A.L. 2019. Uncovering the genomic signature of ancient introgression between white oak lineages (Quercus). New Phytol.:nph.15842.

Dickinson T.A., Lo E., Talent N. 2007. Polyploidy, reproductive biology, and Rosaceae: understanding evolution and making classifications. Plant Syst. Evol. 266:59–78.

Dobeš C., Lückl A., Kausche L., Scheffknecht S., Prohaska D., Sykora C., Paule J. 2015. Parallel origins of apomixis in two diverged evolutionary lineages in tribe Potentilleae (Rosaceae): Origin of Apomixis in Potentilleae. Bot. J. Linn. Soc. 177:214–229.

Doyle J.J., Doyle J.L. 1987. A rapid DNA isolation procedure for small quantities of fresh leaf tissue. Phytochem. Bull. 19:11–15.

Dunn C.W., Howison M., Zapata F. 2013. Agalma: an automated phylogenomics workflow. BMC Bioinformatics. 14:330.

Durand E.Y., Patterson N., Reich D., Slatkin M. 2011. Testing for Ancient Admixture between Closely Related Populations. Mol. Biol. Evol. 28:2239–2252.

Edger P.P., McKain M.R., Bird K.A., VanBuren R. 2018. Subgenome assignment in allopolyploids: challenges and future directions. Curr. Opin. Plant Biol. 42:76–80.

Elworth R.A.L., Allen C., Benedict T., Dulworth P., Nakhleh L.K. 2018. DGEN: A Test Statistic for Detection of General Introgression Scenarios. WABI.

Emms D.M., Kelly S. 2019. OrthoFinder: phylogenetic orthology inference for comparative genomics. Genome Biol. 20:238.

Eriksson T., Lundberg M., Töpel M., Östensson P., Smedmark J.E.E. 2015. Sibbaldia: a molecular phylogenetic study of a remarkably polyphyletic genus in Rosaceae. Plant Syst. Evol. 301:171–184.

Faircloth B.C. 2016. PHYLUCE is a software package for the analysis of conserved genomic loci. Bioinforma. Oxf. Engl. 32:786–788.

Fernández R., Gabaldon T., Dessimoz C. 2020. Orthology: Definitions, Prediction, and Impact on Species Phylogeny Inference. In: Scornavacca C., Delsuc F., Galtier N., editors. Phylogenetics in the Genomic Era. No commercial publisher | Authors open access book. p. 2.4:1--2.4:14.

Fitch W.M. 1970. Distinguishing Homologous from Analogous Proteins. Syst. Biol. 19:99–113.

Forrest L.L., Hart M.L., Hughes M., Wilson H.P., Chung K.-F., Tseng Y.-H., Kidner C.A. 2019. The Limits of Hyb-Seq for Herbarium Specimens: Impact of Preservation Techniques. Front. Ecol. Evol. 7:439.

Freyman W.A., Johnson M.G., Rothfels C.J. 2020. homologizer: Phylogenetic phasing of gene copies into polyploid subgenomes. bioRxiv 2020.10.22.351486

García N., Folk R.A., Meerow A.W., Chamala S., Gitzendanner M.A., Oliveira R.S. de, Soltis D.E., Soltis P.S. 2017. Deep reticulation and incomplete lineage sorting obscure the diploid phylogeny of rain-lilies and allies (Amaryllidaceae tribe Hippeastreae). Mol. Phylogenet. Evol. 111:231–247.

Gardner E.M., Johnson M.G., Pereira J.T., Ahmad Puad A.S., Arifiani D., Sahromi, Wickett N.J., Zerega N.J.C. 2020. Paralogs and off-target sequences improve phylogenetic resolution in a densely-sampled study of the breadfruit genus (Artocarpus, Moraceae). Syst. Bio. syaa073.

Gardner E.M., Johnson M.G., Ragone D., Wickett N.J., Zerega N.J.C. 2016. Low-coverage, whole-genome sequencing of Artocarpus camansi (Moraceae) for phylogenetic marker development and gene discovery. Appl. Plant Sci. 4:1600017.

Gehrke B., Bräuchler C., Romoleroux K., Lundberg M., Heubl G., Eriksson T. 2008. Molecular phylogenetics of Alchemilla, Aphanes and Lachemilla (Rosaceae) inferred from plastid and nuclear intron and spacer DNA sequences, with comments on generic classification. Mol. Phylogenet. Evol. 47:1030–1044.

Gehrke B., Kandziora M., Pirie M.D. 2016. The evolution of dwarf shrubs in alpine environments: a case study of Alchemilla in Africa. Ann. Bot. 117:121–131.

Glover N., Dessimoz C., Ebersberger I., Forslund S.K., Gabaldón T., Huerta-Cepas J., Martin M.-J., Muffato M., Patricio M., Pereira C., da Silva A.S., Wang Y., Sonnhammer E., Thomas P.D. 2019. Advances and Applications in the Quest for Orthologs. Mol. Biol. Evol. 36:2157–2164.

Gonçalves D.J.P., Simpson B.B., Ortiz E.M., Shimizu G.H., Jansen R.K. 2019. Incongruence between gene trees and species trees and phylogenetic signal variation in plastid genes. Mol. Phylogenet. Evol. 138:219–232.

Green R.E., Krause J., Briggs A.W., Maricic T., Stenzel U., Kircher M., Patterson N., Li H., Zhai W., Fritz M.H.Y., Hansen N.F., Durand E.Y., Malaspinas A.S., Jensen J.D., Marques-Bonet T., Alkan C., Prufer K., Meyer M., Burbano H.A., Good J.M., Schultz R., Aximu-Petri A., Butthof A., Hober B., Hoffner B., Siegemund M., Weihmann A., Nusbaum C., Lander E.S., Russ C., Novod N., Affourtit J., Egholm M., Verna C., Rudan P., Brajkovic D., Kucan Z., Gusic I., Doronichev V.B., Golovanova L.V., Lalueza-Fox C., de la Rasilla M., Fortea J., Rosas A., Schmitz R.W., Johnson P.L.F., Eichler E.E., Falush D., Birney E., Mullikin J.C., Slatkin M., Nielsen R., Kelso J., Lachmann M., Reich D., Paabo S. 2010. A Draft Sequence of the Neandertal Genome. Science. 328:710–722.

Guo X., Mandáková T., Trachtová K., Özüdoğru B., Liu J., Lysak M.A. 2020. Linked by ancestral bonds: multiple whole-genome duplications and reticulate evolution in a Brassicaceae tribe. Mol. Biol. Evol. msaa327.

Hayirhoğlu-Ayaz S., İnceer H., Frost-Olsen P. 2006. Chromosome counts in the genus *Alchemilla* (Rosaceae) from SW Europe. Folia Geobot. 41:335–344.

Hejase H.A., Liu K.J. 2016. A scalability study of phylogenetic network inference methods using empirical datasets and simulations involving a single reticulation. BMC Bioinformatics. 17:422.

Hjelmqvist H. 1956. The embryology of some African *Alchemilla* species. Bot. Not. 109:21–32.

Huang C.-H., Zhang C., Liu M., Hu Y., Gao T., Qi J., Ma H. 2016. Multiple Polyploidization Events across Asteraceae with Two Nested Events in the Early History Revealed by Nuclear Phylogenomics. Mol. Biol. Evol. 33:2820–2835.

Izmailow R. Karyological studies in species of *Alchemilla* L. from the series *Calycinae* Bus. (section *Brevicaulon* Rothm.). Acta Biol. Cracoviensia Ser. Bot. 23:117–130.

Jiao Y., Wickett N.J., Ayyampalayam S., Chanderbali A.S., Landherr L., Ralph P.E., Tomsho L.P., Hu Y., Liang H., Soltis P.S., Soltis D.E., Clifton S.W., Schlarbaum S.E., Schuster S.C., Ma H., Leebens-Mack J., dePamphilis C.W. 2011. Ancestral polyploidy in seed plants and angiosperms. Nature. 473:97–100.

Jiao Y., Leebens-Mack J., Ayyampalayam S., Bowers J.E., McKain M.R., McNeal J., Rolf M., Ruzicka D.R., Wafula E., Wickett N.J., Wu X., Zhang Y., Wang J., Zhang Y., Carpenter E.J., Deyholos M.K., Kutchan T.M., Chanderbali A.S., Soltis P.S., Stevenson D.W., McCombie R., Pires J., Wong G., Soltis D.E., dePamphilis C.W. 2012. A genome triplication associated with early diversification of the core eudicots. Genome Biol. 13:R3.

Johnson M.G., Gardner E.M., Liu Y., Medina R., Goffinet B., Shaw A.J., Zerega N.J.C., Wickett N.J. 2016. HybPiper: Extracting Coding Sequence and Introns for Phylogenetics from High-Throughput Sequencing Reads Using Target Enrichment. Appl. Plant Sci. 4:1600016.

Jones K.E., Fér T., Schmickl R.E., Dikow R.B., Funk V.A., Herrando-Moraira S., Johnston P.R., Kilian N., Siniscalchi C.M., Susanna A., Slovák M., Thapa R., Watson L.E., Mandel J.R. 2019. An empirical assessment of a single family-wide hybrid capture locus set at multiple evolutionary timescales in Asteraceae. Appl. Plant Sci. 7:e11295.

Kamneva O.K., Rosenberg N.A. 2017. Simulation-Based Evaluation of Hybridization Network Reconstruction Methods in the Presence of Incomplete Lineage Sorting. Evol. Bioinforma. 13:117693431769193.

Kamneva O.K., Syring J., Liston A., Rosenberg N.A. 2017. Evaluating allopolyploid origins in strawberries (Fragaria) using haplotypes generated from target capture sequencing. BMC Evol. Biol. 17:401.

Karimi N., Grover C.E., Gallagher J.P., Wendel J.F., Ané C., Baum D.A. 2020. Reticulate Evolution Helps Explain Apparent Homoplasy in Floral Biology and Pollination in Baobabs (Adansonia; Bombacoideae; Malvaceae). Syst. Biol. 69:462–478.

Katoh K., Standley D.M. 2013. MAFFT Multiple Sequence Alignment Software Version 7: Improvements in Performance and Usability. Mol. Biol. Evol. 30:772–780.

Kates H.R., Johnson M.G., Gardner E.M., Zerega N.J.C., Wickett N.J. 2018. Allele phasing has minimal impact on phylogenetic reconstruction from targeted nuclear gene sequences in a case study of Artocarpus. Am. J. Bot. 105:404–416.

Kearse M., Moir R., Wilson A., Stones-Havas S., Cheung M., Sturrock S., Buxton S., Cooper A., Markowitz S., Duran C., Thierer T., Ashton B., Meintjes P., Drummond A. 2012. Geneious Basic: An integrated and extendable desktop software platform for the organization and analysis of sequence data. Bioinformatics. 28:1647–1649.

Kocot K.M., Citarella M.R., Moroz L.L., Halanych K.M. 2013. PhyloTreePruner: A Phylogenetic Tree-Based Approach for Selection of Orthologous Sequences for Phylogenomics. Evol. Bioinforma. Online. 9:429–435.

Koenen E.J.M., Ojeda D.I., Bakker F.T., Wieringa J.J., Kidner C., Hardy O.J., Pennington R.T., Herendeen P.S., Bruneau A., Hughes C.E. 2020. The Origin of the Legumes is a Complex Paleopolyploid Phylogenomic Tangle closely associated with the Cretaceous-Paleogene (K-Pg) Mass Extinction Event. Syst. Biol. syaa041.

Larridon I., Villaverde T., Zuntini A.R., Pokorny L., Brewer G.E., Epitawalage N., Fairlie I., Hahn M., Kim J., Maguilla E., Maurin O., Xanthos M., Hipp A.L., Forest F., Baker W.J. 2020. Tackling Rapid Radiations With Targeted Sequencing. Front. Plant Sci. 10:1655.

Leebens-Mack J.H., Barker M.S., Carpenter E.J., Deyholos M.K., Gitzendanner M.A., Graham S.W., Grosse I., Li Z., Melkonian M., Mirarab S., Porsch M., Quint M., Rensing S.A., Soltis D.E., Soltis P.S., Stevenson D.W., Ullrich K.K., Wickett N.J., DeGironimo L., Edger P.P., Jordon-Thaden I.E., Joya S., Liu T., Melkonian B., Miles N.W., Pokorny L., Quigley C., Thomas P., Villarreal J.C., Augustin M.M., Barrett M.D., Baucom R.S., Beerling D.J., Benstein R.M., Biffin E., Brockington S.F., Burge D.O., Burris J.N., Burris K.P., Burtet-Sarramegna V., Caicedo A.L., Cannon S.B., Çebi Z., Chang Y., Chater C., Cheeseman J.M., Chen T., Clarke N.D., Clayton H., Covshoff S., Crandall-Stotler B.J., Cross H., dePamphilis C.W., Der J.P., Determann R., Dickson R.C., Di Stilio V.S., Ellis S., Fast E., Feja N., Field K.J., Filatov D.A., Finnegan P.M., Floyd S.K., Fogliani B., García N., Gâteblé G., Godden G.T., Goh F. (Qi Y., Greiner S., Harkess A., Heaney J.M., Helliwell K.E., Heyduk K., Hibberd J.M., Hodel R.G.J., Hollingsworth P.M., Johnson M.T.J., Jost R., Joyce B., Kapralov M.V., Kazamia E., Kellogg E.A., Koch M.A., Von Konrat M., Könyves K., Kutchan T.M., Lam V., Larsson A., Leitch A.R., Lentz R., Li F.-W., Lowe A.J., Ludwig M., Manos P.S., Mavrodiev E., McCormick M.K., McKain M., McLellan T., McNeal J.R., Miller R.E., Nelson M.N., Peng Y., Ralph P., Real D., Riggins C.W., Ruhsam M., Sage R.F., Sakai A.K., Scascitella M., Schilling E.E., Schlösser E.-M., Sederoff H., Servick S., Sessa E.B., Shaw A.J., Shaw S.W., Sigel E.M., Skema C., Smith A.G., Smithson A., Stewart C.N., Stinchcombe J.R., Szövényi P., Tate J.A., Tiebel H., Trapnell D., Villegente M., Wang C.-N., Weller S.G., Wenzel M., Weststrand S., Westwood J.H., Whigham D.F., Wu S., Wulff A.S., Yang Y., Zhu D., Zhuang C., Zuidof J., Chase M.W., Pires J.C., Rothfels C.J., Yu J., Chen C., Chen L., Cheng S., Li J., Li R., Li X., Lu H., Ou Y., Sun X., Tan X., Tang J., Tian Z., Wang F., Wang J., Wei X., Xu X., Yan Z., Yang F., Zhong X., Zhou F., Zhu Y., Zhang Y., Ayyampalayam S., Barkman T.J., Nguyen N., Matasci N., Nelson D.R., Sayyari E., Wafula E.K., Walls R.L., Warnow T., An H., Arrigo N., Baniaga A.E., Galuska S., Jorgensen S.A., Kidder T.I., Kong H., Lu-Irving P., Marx H.E., Qi X., Reardon C.R., Sutherland B.L., Tiley G.P., Welles S.R., Yu R., Zhan S., Gramzow L., Theißen G., Wong G.K.-S., One Thousand Plant Transcriptomes Initiative. 2019. One thousand plant transcriptomes and the phylogenomics of green plants. Nature. 574:679–685.

Li L. 2003. OrthoMCL: Identification of Ortholog Groups for Eukaryotic Genomes. Genome Res. 13:2178–2189.

Li Z., Baniaga A.E., Sessa E.B., Scascitelli M., Graham S.W., Rieseberg L.H., Barker M.S. 2015. Early genome duplications in conifers and other seed plants. Sci. Adv. 1:e1501084.

Liu Y., Johnson M.G., Cox C.J., Medina R., Devos N., Vanderpoorten A., Hedenäs L., Bell N.E., Shevock J.R., Aguero B., Quandt D., Wickett N.J., Shaw A.J., Goffinet B. 2019. Resolution of the ordinal phylogeny of mosses using targeted exons from organellar and nuclear genomes. Nat. Commun. 10:1485.

Lundberg M., Töpel M., Eriksen B., Nylander J.A.A., Eriksson T. 2009. Allopolyploidy in Fragariinae (Rosaceae): Comparing four DNA sequence regions, with comments on classification. Mol. Phylogenet. Evol. 51:269–280.

Lynch M., Conery J.S. 2000. The Evolutionary Fate and Consequences of Duplicate Genes. Science. 290:1151–1155.

Mai U., Mirarab S. 2018. TreeShrink: fast and accurate detection of outlier long branches in collections of phylogenetic trees. BMC Genomics. 19:272.

Mandáková T., Lysak M.A. 2018. Post-polyploid diploidization and diversification through dysploid changes. Curr. Opin. Plant Biol. 42:55–65.

Mandel J.R., Dikow R.B., Funk V.A., Masalia R.R., Staton S.E., Kozik A., Michelmore R.W., Rieseberg L.H., Burke J.M. 2014. A Target Enrichment Method for Gathering Phylogenetic Information from Hundreds of Loci: An Example from the Compositae. Appl. Plant Sci. 2:1300085.

Nauheimer L., Weigner N., Joyce E., Crayn D., Clarke C., Nargar K. 2020. HybPhaser: a workflow for the detection and phasing of hybrids in target capture datasets. bioRxiv 2020.10.27.354589

McKain M.R., Estep M.C., Pasquet R., Layton D.J., Vela Díaz D.M., Zhong J., Hodge J.G., Malcomber S.T., Chipabika G., Pallangyo B., Kellogg E.A. 2018. Ancestry of the two subgenomes of maize. bioRxiv.:352351.

McKain M.R., Tang H., McNeal J.R., Ayyampalayam S., Davis J.I., dePamphilis C.W., Givnish T.J., Pires J.C., Stevenson D.W., Leebens-Mack J.H. 2016. A Phylogenomic Assessment of Ancient Polyploidy and Genome Evolution across the Poales. Genome Biol. Evol. 8:1150–1164.

McLachlan G., Peel D. 2000. Finite Mixture Models. New York: Wiley.

Molloy E.K., Warnow T. 2020. FastMulRFS: fast and accurate species tree estimation under generic gene duplication and loss models. Bioinformatics. 36:i57–i65.

Montes J.R., Peláez P., Willyard A., Moreno-Letelier A., Piñero D., Gernandt D.S. 2019. Phylogenetics of Pinus Subsection Cembroides Engelm. (Pinaceae) Inferred from Low-Copy Nuclear Gene Sequences. Syst. Bot. 44:501–518.

Montgomery L. 1997. Contributions to a cytological catalogue of the British and Irish flora, 5. Watsonia. 21:365–368.

Moore A.J., Vos J.M.D., Hancock L.P., Goolsby E., Edwards E.J. 2018. Targeted Enrichment of Large Gene Families for Phylogenetic Inference: Phylogeny and Molecular Evolution of Photosynthesis Genes in the Portullugo Clade (Caryophyllales). Syst. Biol. 67:367–383.

Morales-Briones D.F., Romoleroux K., Kolář F., Tank D.C. 2018a. Phylogeny and Evolution of the Neotropical Radiation of *Lachemilla* (Rosaceae): Uncovering a History of Reticulate Evolution and Implications for Infrageneric Classification. Syst. Bot. 43:17–34.

Morales-Briones D.F., Liston A., Tank D.C. 2018b. Phylogenomic analyses reveal a deep history of hybridization and polyploidy in the Neotropical genus *Lachemilla* (Rosaceae). New Phytol. 218:1668–1684.

Morales-Briones D.F., Tank D.C. 2019. Extensive allopolyploidy in the neotropical genus *Lachemilla* (Rosaceae) revealed by. Am. J. Bot. 106:415–437.

Morales-Briones D.F., Kadereit G., Tefarikis D.T., Moore M.J., Smith S.A., Brockington S.F., Timoneda A., Yim W.C., Cushman J.C., Yang Y. 2021. Disentangling Sources of Gene Tree Discordance in Phylogenomic Datasets: Testing Ancient Hybridizations in Amaranthaceae s.l. Syst. Biol. 70:219–235

Morton J. 1993. Chromosome numbers and polyploidy in the flora of Cameroons Mountain. Opera Bot. 121:159–172.

Nicholls J.A., Pennington R.T., Koenen E.J.M., Hughes C.E., Hearn J., Bunnefeld L., Dexter K.G., Stone G.N., Kidner C.A. 2015. Using targeted enrichment of nuclear genes to increase phylogenetic resolution in the neotropical rain forest genus Inga (Leguminosae: Mimosoideae). Front. Plant Sci. 6:710.

Nute M., Chou J., Molloy E.K., Warnow T. 2018. The performance of coalescent-based species tree estimation methods under models of missing data. BMC Genomics. 19:286.

Panchy N., Lehti-Shiu M., Shiu S.-H. 2016. Evolution of Gene Duplication in Plants. Plant Physiol. 171:2294–2316.

Pease J.B., Brown J.W., Walker J.F., Hinchliff C.E., Smith S.A. 2018. Quartet Sampling distinguishes lack of support from conflicting support in the green plant tree of life. Am. J. Bot. 105:385–403.

Perry L.M. 1929. A Tentative Revision of Alchemilla § Lachemilla. Contrib. Gray Herb. Harv. Univ.:1–57.

Ranwez V., Douzery E.J.P., Cambon C., Chantret N., Delsuc F. 2018. MACSE v2: Toolkit for the Alignment of Coding Sequences Accounting for Frameshifts and Stop Codons. Mol. Biol. Evol. 35:2582–2584.

Salichos L., Stamatakis A., Rokas A. 2014. Novel Information Theory-Based Measures for Quantifying Incongruence among Phylogenetic Trees. Mol. Biol. Evol. 31:1261–1271.

Sayyari E., Mirarab S. 2016. Fast Coalescent-Based Computation of Local Branch Support from Quartet Frequencies. Mol. Biol. Evol. 33:1654–1668.

Shulaev V., Sargent D.J., Crowhurst R.N., Mockler T.C., Folkerts O., Delcher A.L., Jaiswal P., Mockaitis K., Liston A., Mane S.P., Burns P., Davis T.M., Slovin J.P., Bassil N., Hellens R.P., Evans C., Harkins T., Kodira C., Desany B., Crasta O.R., Jensen R.V., Allan A.C., Michael T.P., Setubal J.C., Celton J.-M., Rees D.J.G., Williams K.P., Holt S.H., Rojas J.J.R., Chatterjee M., Liu B., Silva H., Meisel L., Adato A., Filichkin S.A., Troggio M., Viola R., Ashman T.-L., Wang H., Dharmawardhana P., Elser J., Raja R., Priest H.D., Bryant D.W., Fox S.E., Givan S.A., Wilhelm L.J., Naithani S., Christoffels A., Salama D.Y., Carter J., Girona E.L., Zdepski A., Wang W., Kerstetter R.A., Schwab W., Korban S.S., Davik J., Monfort A., Denoyes-Rothan B., Arus P., Mittler R., Flinn B., Aharoni A., Bennetzen J.L., Salzberg S.L., Dickerman A.W., Velasco R., Borodovsky M., Veilleux R.E., Folta K.M. 2011. The genome of woodland strawberry (Fragaria vesca). Nat. Genet. 43:109–116.

Smedmark J.E.E., Eriksson T., Evans R.C., Campbell C.S. 2003. Ancient Allopolyploid Speciation in Geinae (Rosaceae): Evidence from Nuclear Granule-Bound Starch Synthase (GBSSI) Gene Sequences. Syst. Biol. 52:374–385.

Smith M.L., Hahn M.W. 2020. New Approaches for Inferring Phylogenies in the Presence of Paralogs. Trends Genet.

Smith S.A., Moore M.J., Brown J.W., Yang Y. 2015. Analysis of phylogenomic datasets reveals conflict, concordance, and gene duplications with examples from animals and plants. BMC Evol. Biol. 15:150.

Soják J. 2008. Notes on Potentilla XXI. A new division of the tribe Potentilleae (Rosaceae) and notes on generic delimitations. Bot. Jahrb. Für Syst. Pflanzengesch. Pflanzengeogr. 127:349–358.

Solís-Lemus C., Ané C. 2016. Inferring Phylogenetic Networks with Maximum Pseudolikelihood under Incomplete Lineage Sorting. PLoS Genet. 12:e1005896–21.

Stamatakis A. 2014. RAxML version 8 - a tool for phylogenetic analysis and post-analysis of large phylogenies. Bioinformatics. 30:1312–1313.

Straub S.C., Fishbein M., Livshultz T., Foster Z., Parks M., Weitemier K., Cronn R.C., Liston A. 2011. Building a model: developing genomic resources for common milkweed (Asclepias syriaca) with low coverage genome sequencing. BMC Genomics. 12:211.

Stubbs R.L., Folk R.A., Xiang C.-L., Chen S., Soltis D.E., Cellinese N. 2020. A Phylogenomic Perspective on Evolution and Discordance in the Alpine-Arctic Plant Clade Micranthes (Saxifragaceae). Front. Plant Sci. 10:1773.

Thomas G.W.C., Ather S.H., Hahn M.W. 2017. Gene-Tree Reconciliation with MUL-Trees to Resolve Polyploidy Events. Syst. Biol. 66:1007–1018.

Villaverde T., Pokorny L., Olsson S., Rincón-Barrado M., Johnson M.G., Gardner E.M., Wickett N.J., Molero J., Riina R., Sanmartín I. 2018. Bridging the micro- and macroevolutionary levels in phylogenomics: Hyb-Seq solves relationships from populations to species and above. New Phytol. 220:636–650.

Walker J.F., Walker-Hale N., Vargas O.M., Larson D.A., Stull G.W. 2019. Characterizing gene tree conflict in plastome-inferred phylogenies. PeerJ. 7:e7747.

Walters S., Boznan V. *Alchemilla faeroensis* (Lange) Buser and *A. alpina* L. Proc. Bot. Soc. Br. Isles. 7:83.

Weitemier K., Straub S.C.K., Cronn R.C., Fishbein M., Schmickl R., McDonnell A., Liston A. 2014. Hyb-Seq: Combining Target Enrichment and Genome Skimming for Plant Phylogenomics. Appl. Plant Sci. 2:1400042.

Wen D., Yu Y., Zhu J., Nakhleh L. 2018. Inferring Phylogenetic Networks Using PhyloNet. Syst. Biol. 67:735–740.

Xiang Y., Huang C.-H., Hu Y., Wen J., Li S., Yi T., Chen H., Xiang J., Ma H. 2017. Evolution of Rosaceae Fruit Types Based on Nuclear Phylogeny in the Context of Geological Times and Genome Duplication. Mol. Biol. Evol. 34:262–281.

Yang Y., Moore M.J., Brockington S.F., Mikenas J., Olivieri J., Walker J.F., Smith S.A. 2018. Improved transcriptome sampling pinpoints 26 ancient and more recent polyploidy events in Caryophyllales, including two allopolyploidy events. New Phytol. 217:855– 870.

Yang Y., Moore M.J., Brockington S.F., Soltis D.E., Wong G.K.-S., Carpenter E.J., Zhang Y., Chen L., Yan Z., Xie Y., Sage R.F., Covshoff S., Hibberd J.M., Nelson M.N., Smith S.A. 2015. Dissecting Molecular Evolution in the Highly Diverse Plant Clade Caryophyllales Using Transcriptome Sequencing. Mol. Biol. Evol. 32:2001–2014.

Yang Y., Smith S.A. 2014. Orthology Inference in Nonmodel Organisms Using Transcriptomes and Low-Coverage Genomes: Improving Accuracy and Matrix Occupancy for Phylogenomics. Mol. Biol. Evol. 31:3081–3092.

Zhang C., Rabiee M., Sayyari E., Mirarab S. 2018. ASTRAL-III: polynomial time species tree reconstruction from partially resolved gene trees. BMC Bioinformatics. 19:153.

Zhang C., Scornavacca C., Molloy E.K., Mirarab S. 2020a. ASTRAL-Pro: Quartet-Based Species-Tree Inference despite Paralogy. Mol. Biol. Evol. 37:3292–3307.

Zhang R., Wang Y.-H., Jin J.-J., Stull G.W., Bruneau A., Cardoso D., De Queiroz L.P., Moore M.J., Zhang S.-D., Chen S.-Y., Wang J., Li D.-Z., Yi T.-S. 2020b. Exploration of Plastid Phylogenomic Conflict Yields New Insights into the Deep Relationships of Leguminosae. Syst. Biol. 69:613–622.

Zhbannikov I.Y., Hunter S.S., Foster J.A., Settles M.L. 2017. SeqyClean: A Pipeline for High-throughput Sequence Data Preprocessing. Proc. 8th ACM Int. Conf. Bioinforma. Comput. Biol. Health Inform.:407–416.

