## Supplemental Material for "Analysis of paralogs in target enrichment data pinpoints multiple ancient polyploidy events in *Alchemilla* s.l. (Rosaceae)"

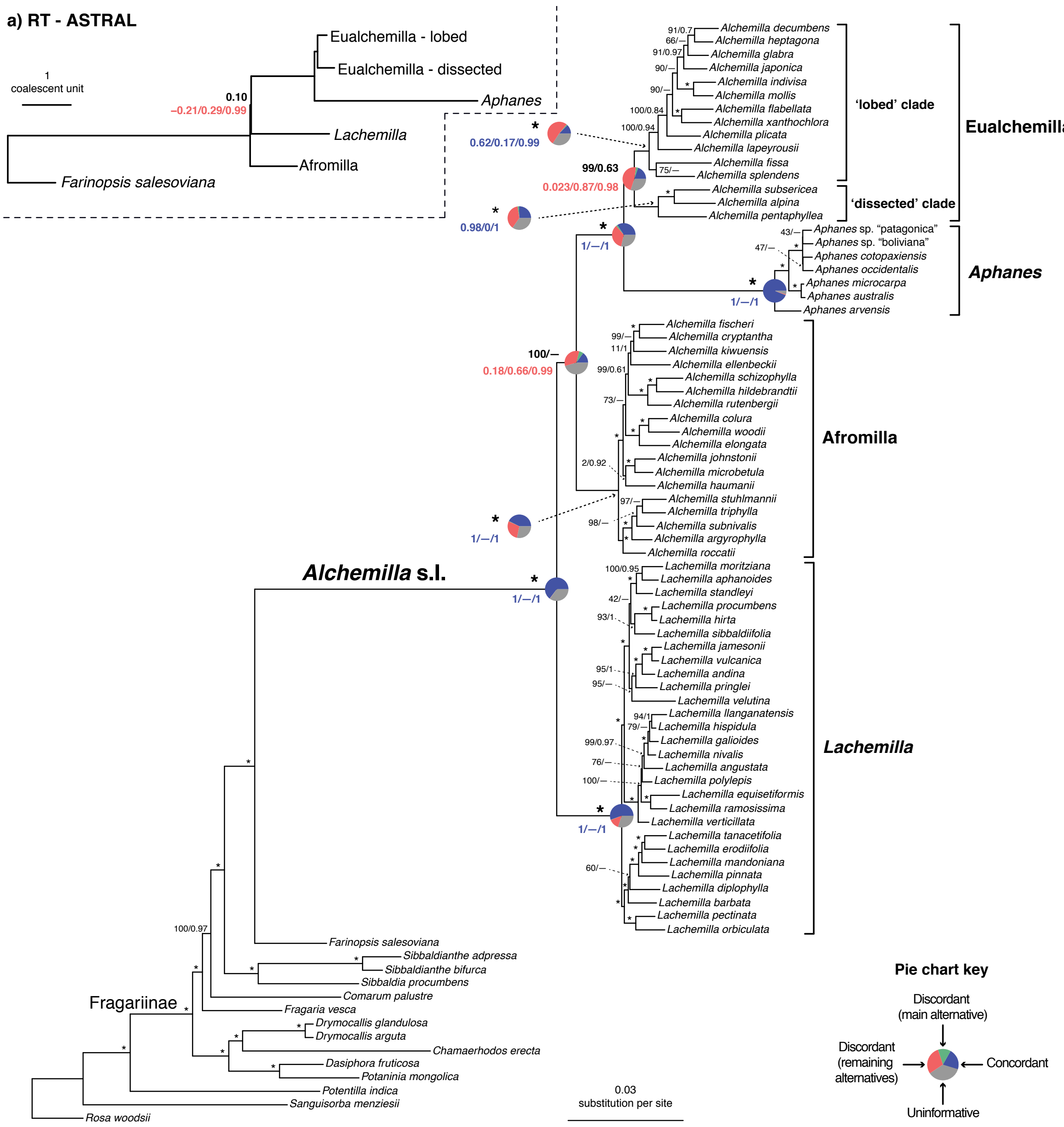


**Figure S1.** Maximum likelihood phylogeny of *Alchemilla* s.l. inferred from RAxML analysis of the concatenated 1,894-exon nuclear supermatrix from the 'rooted ingroup' (RT) orthologs. Bootstrap support (BS) and Local posterior probability (LLP) are shown above branches. Nodes with full support (BS= 100/LLP= 100) are noted with an asterisk (*). Em dashes (—) denoted alternative topology compared to the ASTRAL tree. Quartet Sampling (QS) scores for major clades are shown below branches. QS scores in blue indicate strong support and red scores indicate weak support. QS scores: Quartet concordance/Quartet differential/Quartet informativeness. Pie charts for major clades represent the proportion of ortholog trees that support that clade (blue), the proportion that support the main alternative bifurcation (green), the proportion that support the remaining alternatives (red), and the proportion (conflict or support) that have < 50% bootstrap support (gray). Gene trees with missing data that were uninformative for the node were ignored.  Branch lengths are in number of substitutions per site (scale bar on the bottom). Inset: a) Summary ASTRAL inferred from 1,894 nuclear trees from the RT orthologs. LLP and QS are shown above and below the branch that differ from the RAxML tree. Branch lengths are in coalescent units.

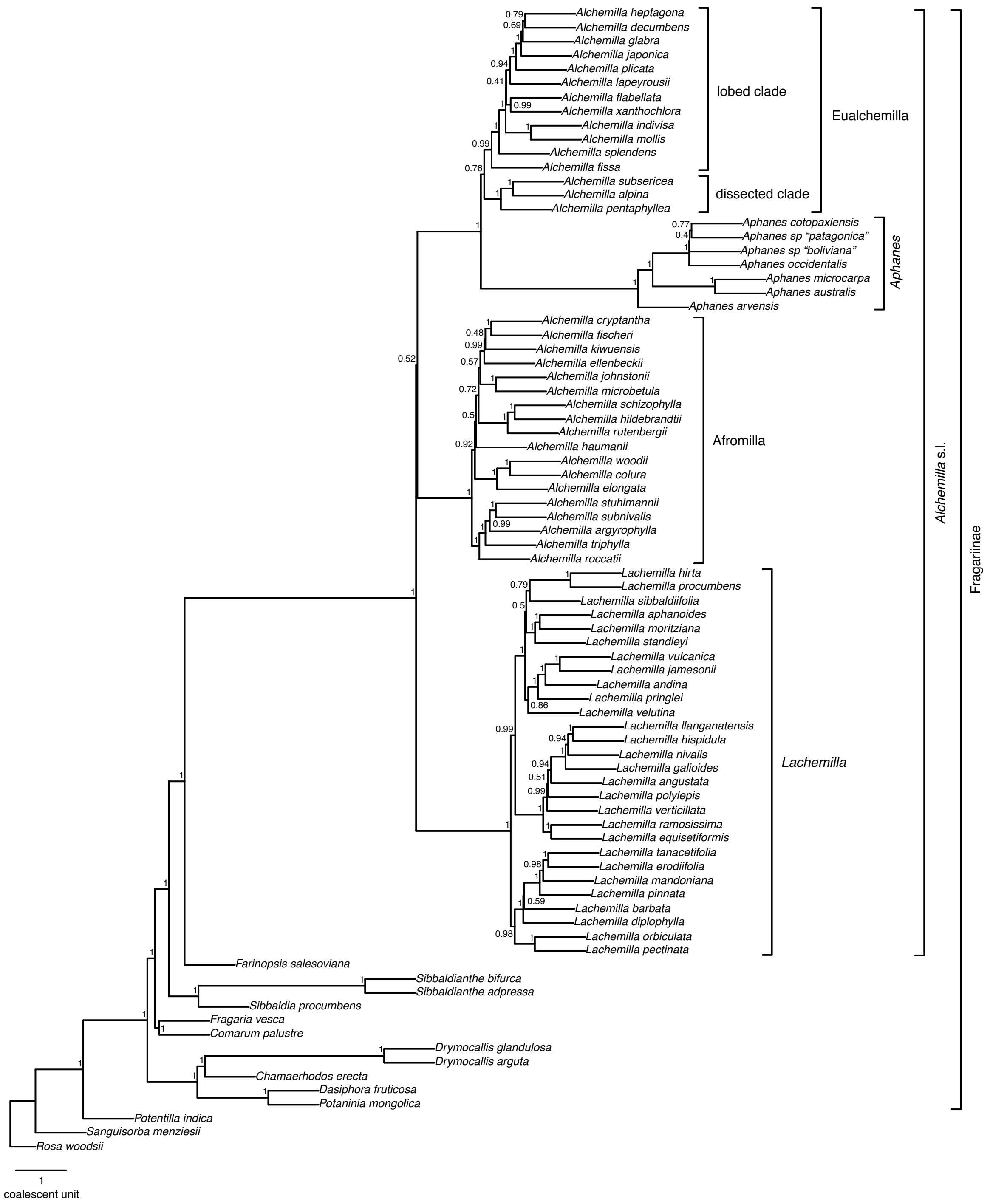


**Figure S2.** ASTRAL-Pro tree of *Alchemilla* s.l. inferred from 923 multi-labeled homolog exon trees. Local posterior probabilities (LLP) are shown next to nodes. Branch lengths are in coalescent units (scale bar on the bottom).

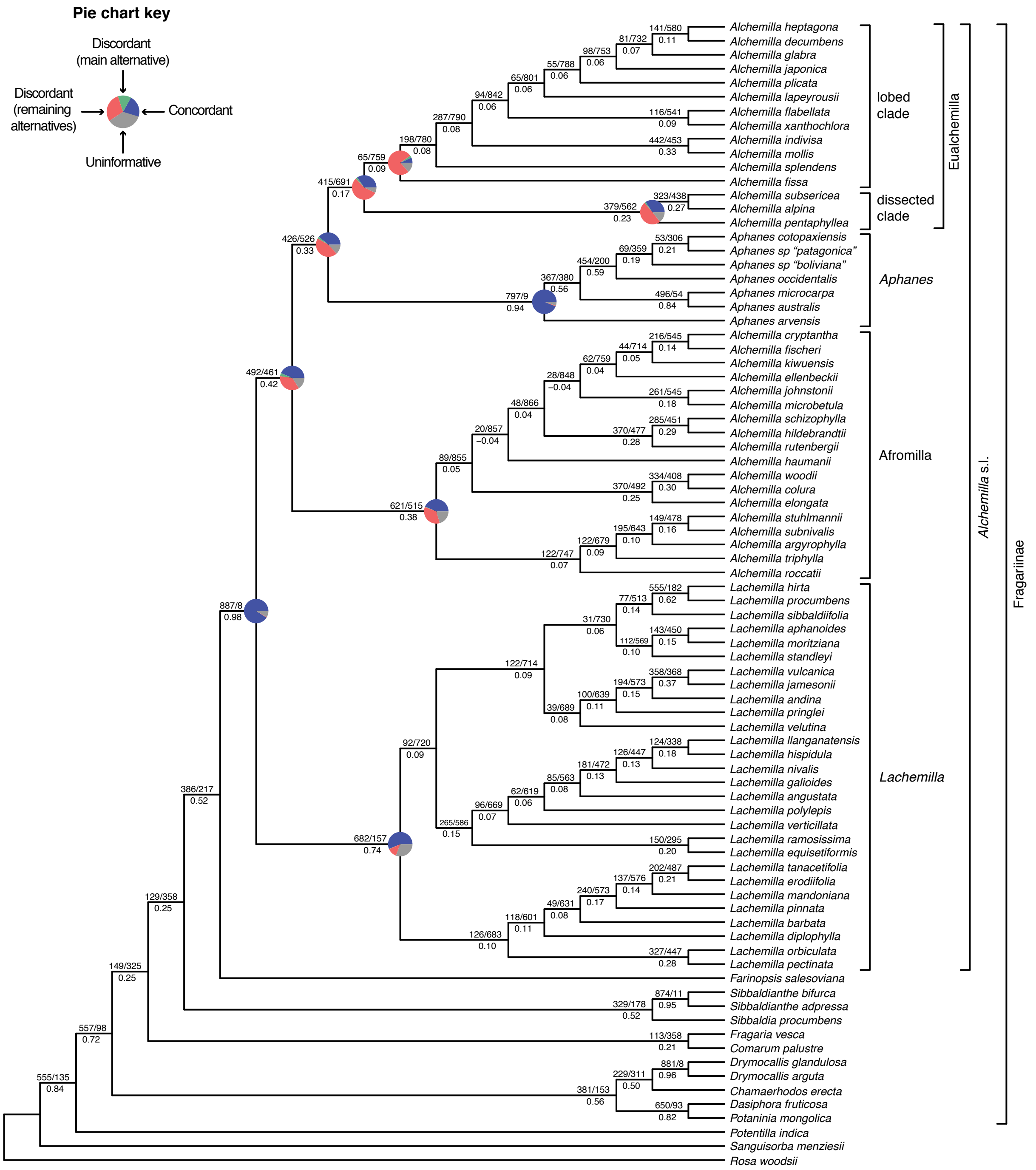


**Figure S3.** ASTRAL-Pro cladogram of *Alchemilla* s.l. inferred from 923 multi-labeled homolog exon trees. Local posterior probabilities (LLP) are shown next to nodes. Numbers above branches indicate the number of trees concordant/conflicting with that node in the ASTRAL-Pro tree. Numbers below the branches are the Internode Certainty All (ICA) score. Pie charts for major clades represent the proportion of ortholog trees that support that clade (blue), the proportion that support the main alternative bifurcation (green), the proportion that support the remaining alternatives (red), and the proportion (conflict or support) that have < 50% bootstrap support (gray).

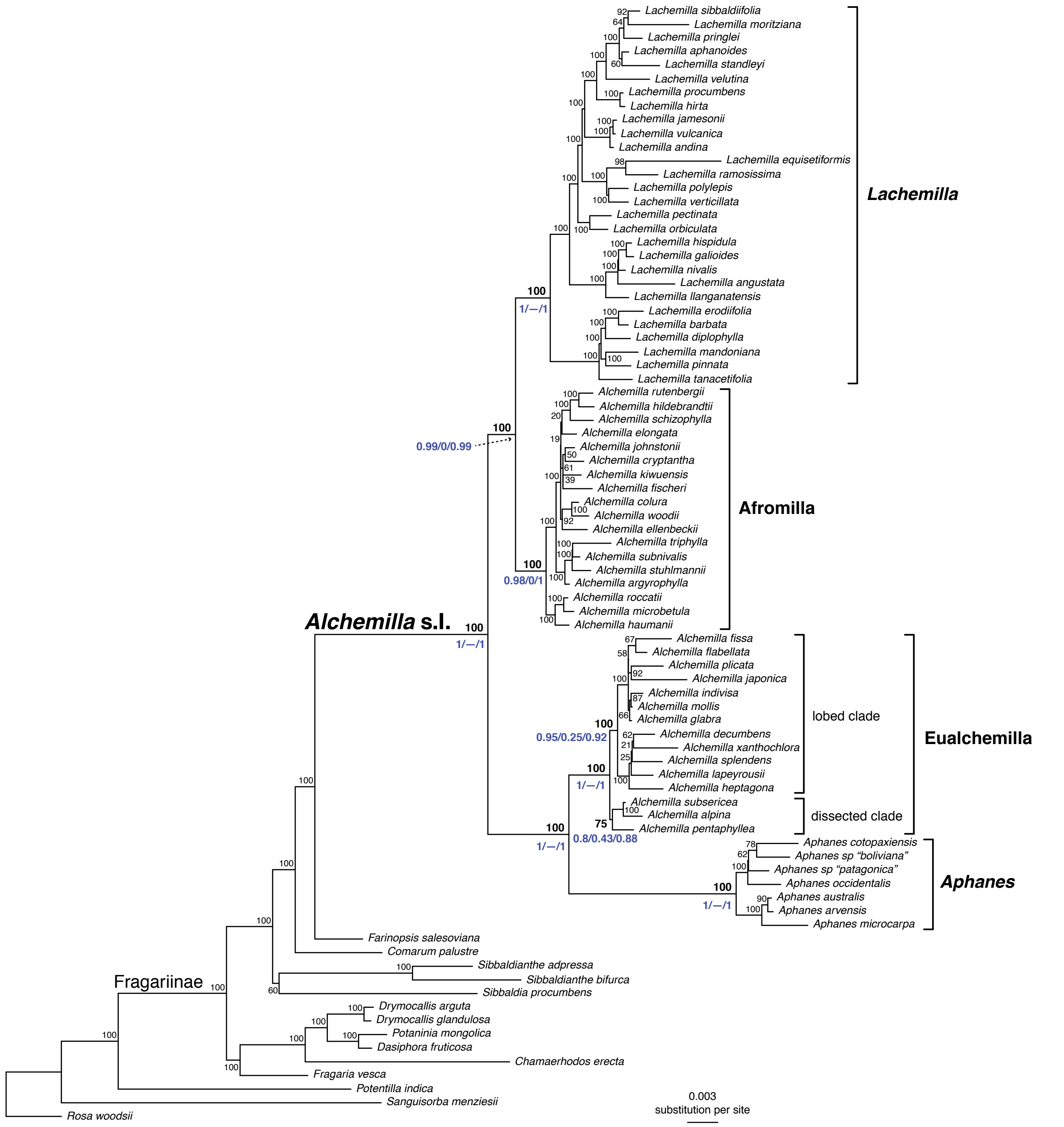


**Figure S4.** Maximum likelihood phylogeny of *Alchemilla* s.l. inferred from RAxML analysis of concatenated partial plastomes. Bootstrap support (BS) is shown above branches. Quartet Sampling (QS) scores for major clades are shown below branches. QS scores in blue indicate strong support and red scores indicate weak support. QS scores: Quartet concordance/Quartet differential/Quartet informativeness. Branch lengths are in number of substitutions per site (scale bar on the bottom).

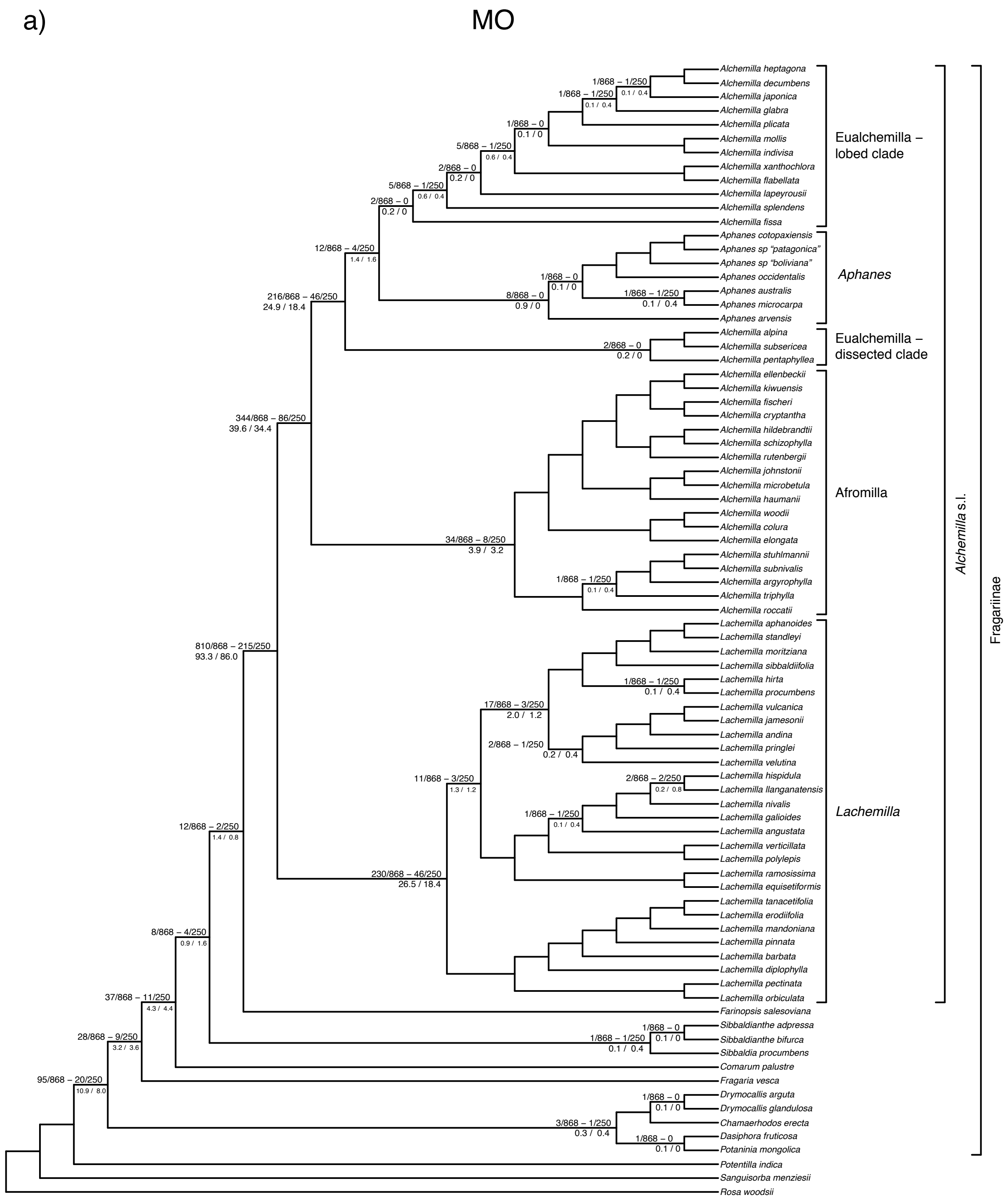
a)
MO
Alchemilla heptagona
Alchemilla decumbens
Alchemilla japonica
Alchemilla glabra
Alchemilla plicata
Alchemilla mollis
Alchemilla indivisa
Alchemilla xanthochlora
Alchemilla flabellata
Alchemilla lapeyrousii
Alchemilla splendens
Alchemilla fissa
Aphanes cotopaxiensis
Aphanes sp "patagonica"
Aphanes sp "boliviana"
Aphanes occidentalis
Aphanes australis
Aphanes microcarpa
Aphanes arvensis
Alchemilla alpina
Alchemilla subsericea
Alchemilla pentaphyllea
Alchemilla ellenbeckii
Alchemilla kiwuensis
Alchemilla fischeri
Alchemilla cryptantha
Alchemilla hildebrandtii
Alchemilla schizophylla
Alchemilla rutenbergii
Alchemilla johnstonii
Alchemilla microbetula
Alchemilla haumanii
Alchemilla woodii
Alchemilla colura
Alchemilla elongata
Alchemilla stuhlmannii
Alchemilla subnivalis
Alchemilla argyrophylla
Alchemilla triphylla
Alchemilla roccatii
Lachemilla aphanoides
Lachemilla standleyi
Lachemilla moritziana
Lachemilla sibbaldiifolia
Lachemilla hirta
Lachemilla procumbens
Lachemilla vulcanica
Lachemilla jamesonii
Lachemilla andina
Lachemilla pringlei
Lachemilla velutina
Lachemilla hispidula
Lachemilla llanganatensis
Lachemilla nivalis
Lachemilla galioides
Lachemilla angustata
Lachemilla verticillata
Lachemilla polylepis
Lachemilla ramosissima
Lachemilla equisetiformis
Lachemilla tanacetifolia
Lachemilla erodiifolia
Lachemilla mandoniana
Lachemilla pinnata
Lachemilla barbata
Lachemilla diplophylla
Lachemilla pectinata
Lachemilla orbiculata
Farinopsis salesoviana
Sibbaldianthe adpressa
Sibbaldianthe bifurca
Sibbaldia procumbens
Comarum palustre
Fragaria vesca
Drymocallis arguta
Drymocallis glandulosa
Chamaerhodos erecta
Dasiphora fruticosa
Potaninia mongolica
Potentilla indica
Sanguisorba menziesii
Rosa woodsii
Eualchemilla – lobed clade
Aphanes
Eualchemilla – dissected clade
Afromilla
Lachemilla
Alchemilla s.l.
Fragariinae
1/868 – 1/250
0.1 / 0.4
1/868 – 1/250
0.1 / 0.4
1/868 – 0
0.1 / 0
5/868 – 1/250
0.6 / 0.4
2/868 – 0
0.2 / 0
5/868 – 1/250
0.6 / 0.4
2/868 – 0
0.2 / 0
12/868 – 4/250
1.4 / 1.6
216/868 – 46/250
24.9 / 18.4
1/868 – 0
0.1 / 0
8/868 – 0
0.9 / 0
1/868 – 1/250
0.1 / 0.4
2/868 – 0
0.2 / 0
344/868 – 86/250
39.6 / 34.4
34/868 – 8/250
3.9 / 3.2
1/868 – 1/250
0.1 / 0.4
810/868 – 215/250
93.3 / 86.0
1/868 – 1/250
0.1 / 0.4
17/868 – 3/250
2.0 / 1.2
2/868 – 1/250
0.2 / 0.4
11/868 – 3/250
1.3 / 1.2
2/868 – 2/250
0.2 / 0.8
1/868 – 1/250
0.1 / 0.4
12/868 – 2/250
1.4 / 0.8
230/868 – 46/250
26.5 / 18.4
8/868 – 4/250
0.9 / 1.6
37/868 – 11/250
4.3 / 4.4
28/868 – 9/250
3.2 / 3.6
1/868 – 0
0.1 / 0
1/868 – 1/250
0.1 / 0.4
95/868 – 20/250
10.9 / 8.0
1/868 – 0
0.1 / 0
3/868 – 1/250
0.3 / 0.4
1/868 – 0
0.1 / 0

b) RT

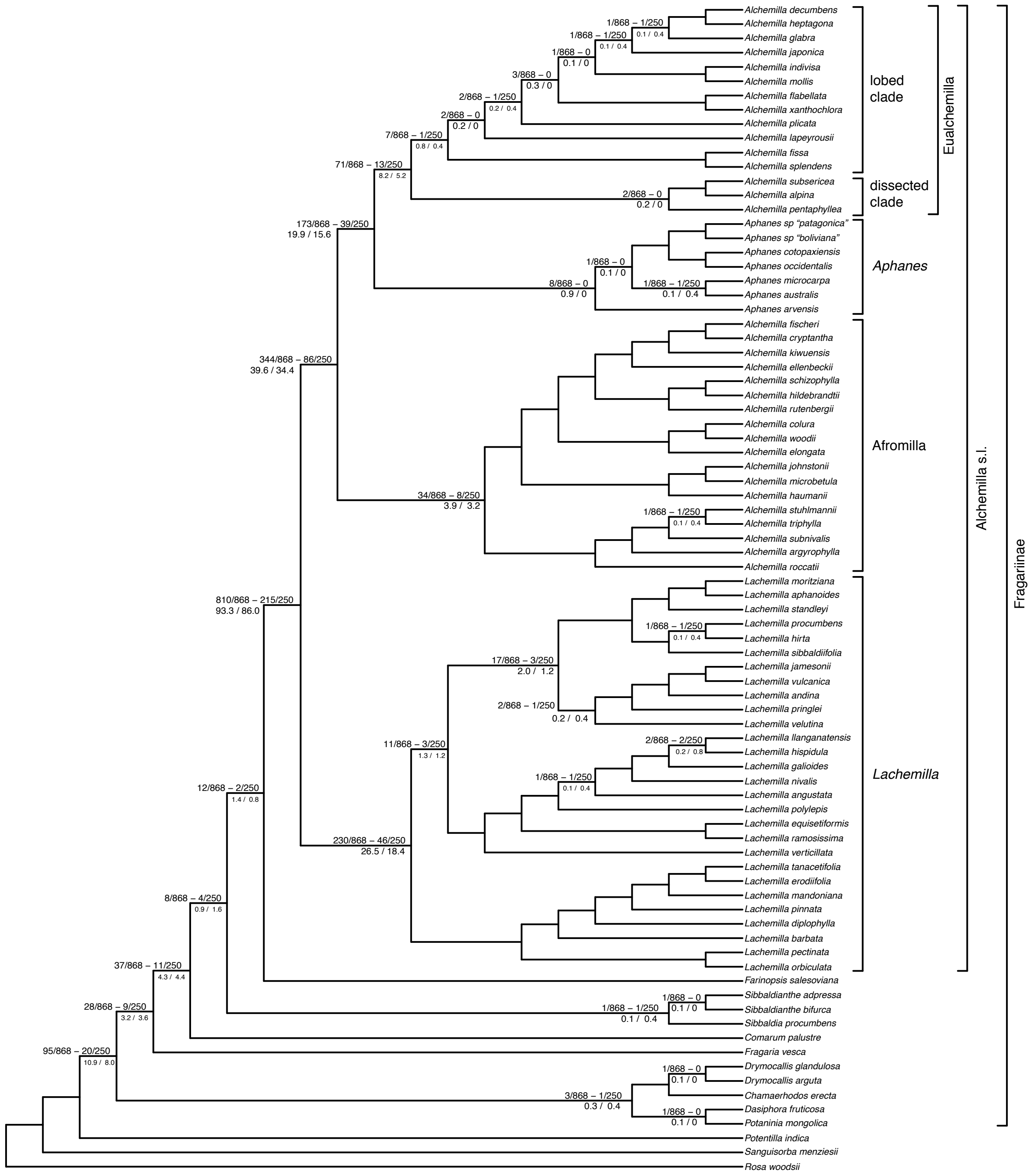


**Figure S5**. a) Maximum likelihood cladogram of *Alchemilla* s.l. inferred from RAxML analysis of the concatenated 910-nuclear exon supermatrix from the 'monophyletic outgroup' (MO) orthologs with gene duplication estimates from the subclade orthogroup tree topology approach. b) Maximum likelihood cladogram of *Alchemilla* s.l. inferred from RAxML analysis of the concatenated 1,894-exon nuclear supermatrix from the 'rooted ingroup' (RT) orthologs with gene duplication estimates from the subclade orthogroup tree topology approach. Numbers below branches denote gene duplication counts when using all homolog exons (868) and only the longest homolog exons per gene (250). Numbers above branches denote gene duplication percentages when using all homologs and only the longest homologs. Unlabeled nodes had no evidence of gene duplication.

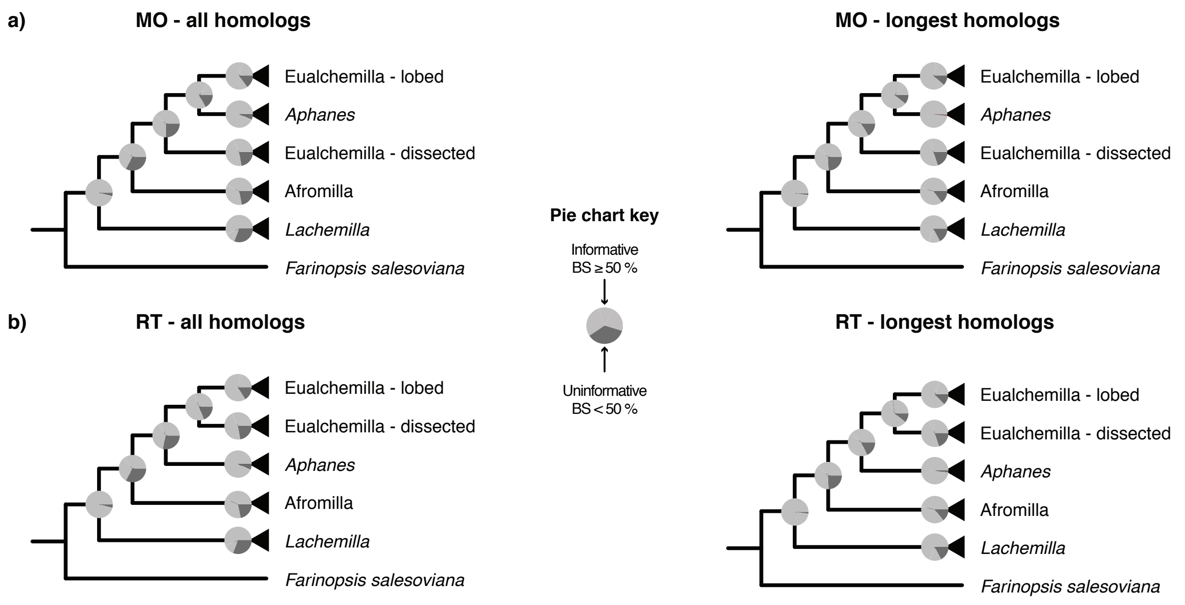


**Figure S6.** Proportion of nodes from homolog exon trees that were considered "informative" [bootstrap support (BS) ≥ 50%; light gray] versus uninformative (BS < 50; dark gray) summarized on a) the MO RAxML tree and b) the RT RAxML tree.

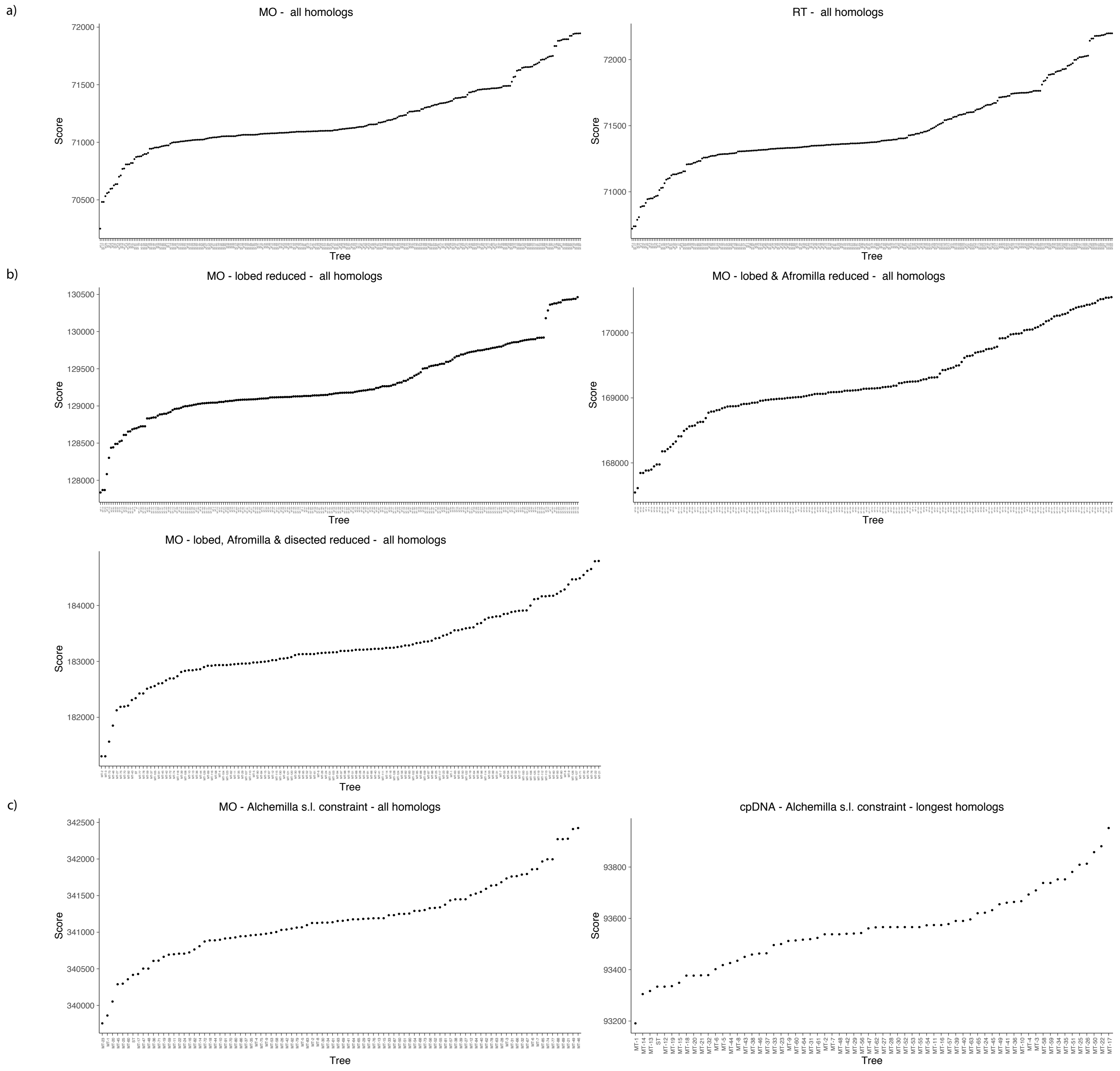


**Figure S7**. Distribution of GRAMPA reconciliations scores. The optimal topology is the one with the lowest reconciliation scores. a) GRAMPA analyses using species trees inferred from 'monophyletic outgroup' (MO) and 'rooted ingroup' (RT) orthologs (Fig. S6). b) GRAMPA analyses using species trees inferred from MO orthologs after sequential removal of major clades of *Alchemilla* s.l. in homolog exon trees (Fig. 4b–d). c) GRAMPA analyses using searches constrained to the base of *Alchemilla* s.l. on the 'monophyletic outgroup' (MO) and cpDNA trees (Fig. S9).

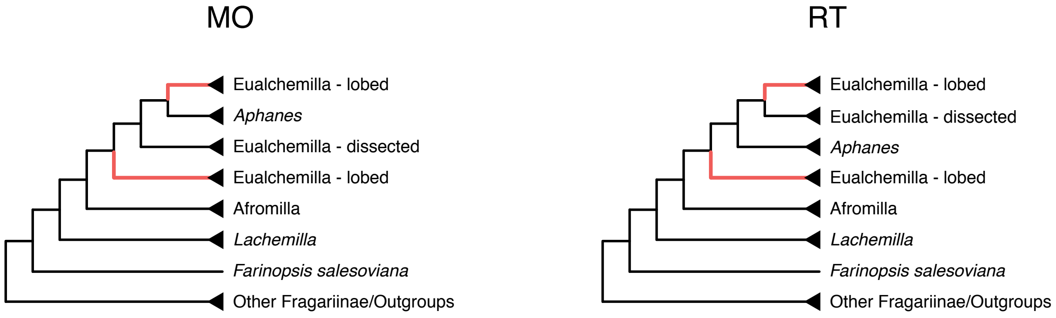


**Figure S8**. Optimal multi-labeled trees from GRAMPA analyses by mapping homolog trees onto species trees inferred using ‘monophyletic outgroup’ (MO) and ‘rooted ingroup’ (RT) ortholog trees. Red branches denote the allopolyploid origin of the 'lobed' clade of Eualchemilla.

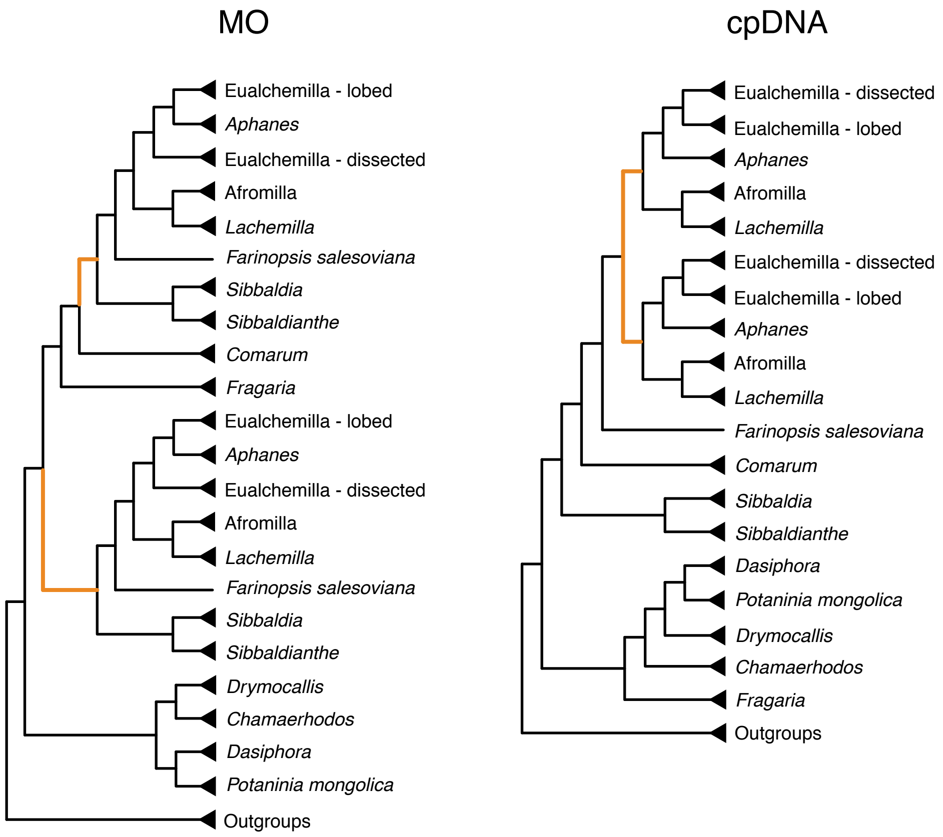


**Figure S9**. Optimal summarized multi-labeled trees from GRAMPA analyses using searches constrained to the crown node of *Alchemilla* s.l. on the species trees inferred from 'monophyletic outgroup' (MO) orthologs and cpDNA. Orange branches denote the allopolyploid origin of the clade composed of *Alchemilla* s.l., *Farinopsis, Sibbaldianthe*, and *Sibbaldia* (MO) or the autopolyploid origin *Alchemilla* s.l. (cpDNA).

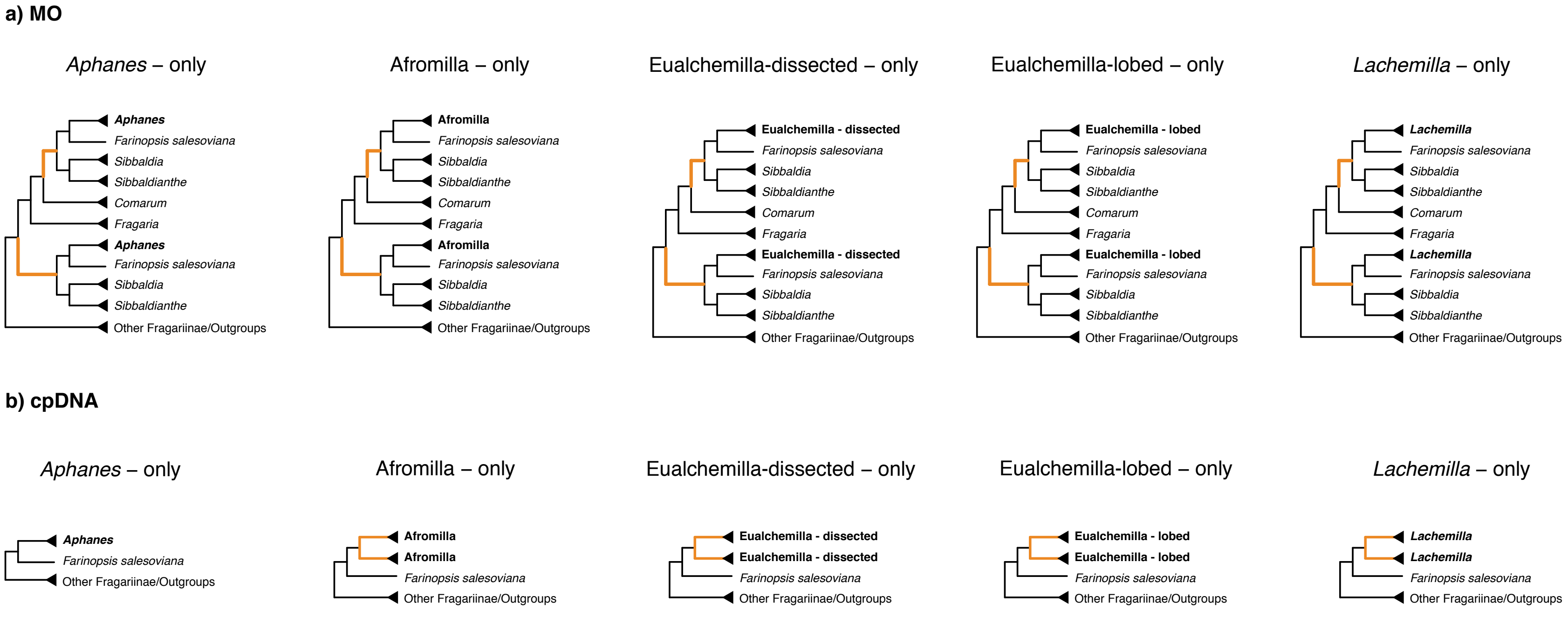


**Figure S10**. Optimal summarized multi-labeled trees from GRAMPA analyses using gene trees removing taxa in *Alchemilla* s.l. except one of the five major clades *Alchemilla* s.l. on to species trees inferred from the 'monophyletic outgroup' (MO) orthologs (a) and cpDNA (b). Orange branches denote the allopolyploid events (MO) or autopolyploid events (cpDNA).

a)

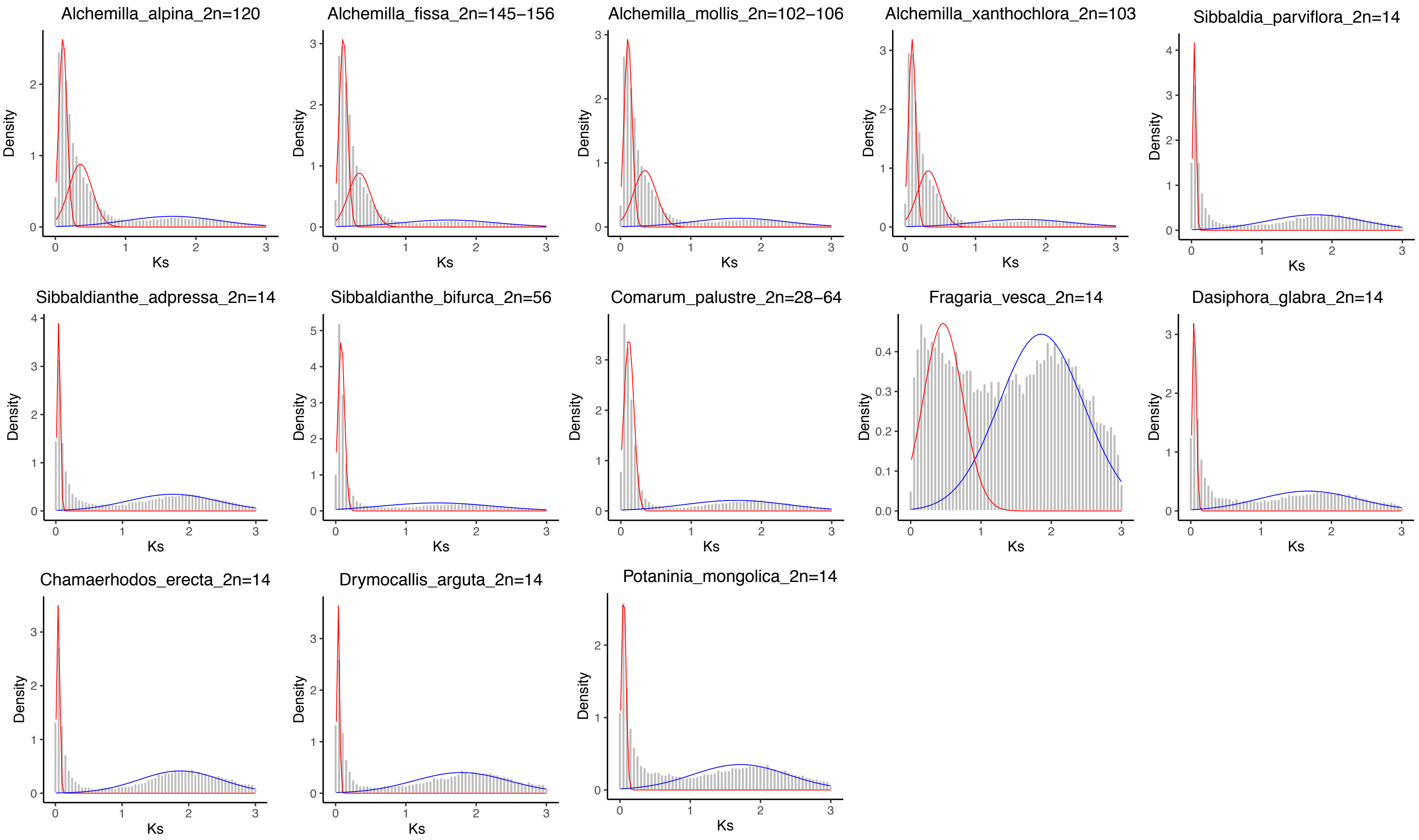

Alchemilla_alpina_2n=120
Alchemilla_fissa_2n=145−156
Alchemilla_mollis_2n=102−106
Alchemilla_xanthochlora_2n=103
Sibbaldia_parviflora_2n=14
Sibbaldianthe_adpressa_2n=14
Sibbaldianthe_bifurca_2n=56
Comarum_palustre_2n=28−64
Fragaria_vesca_2n=14
Dasiphora_glabra_2n=14
Chamaerhodos_erecta_2n=14
Drymocallis_arguta_2n=14
Potaninia_mongolica_2n=14
Density
Ks

b)

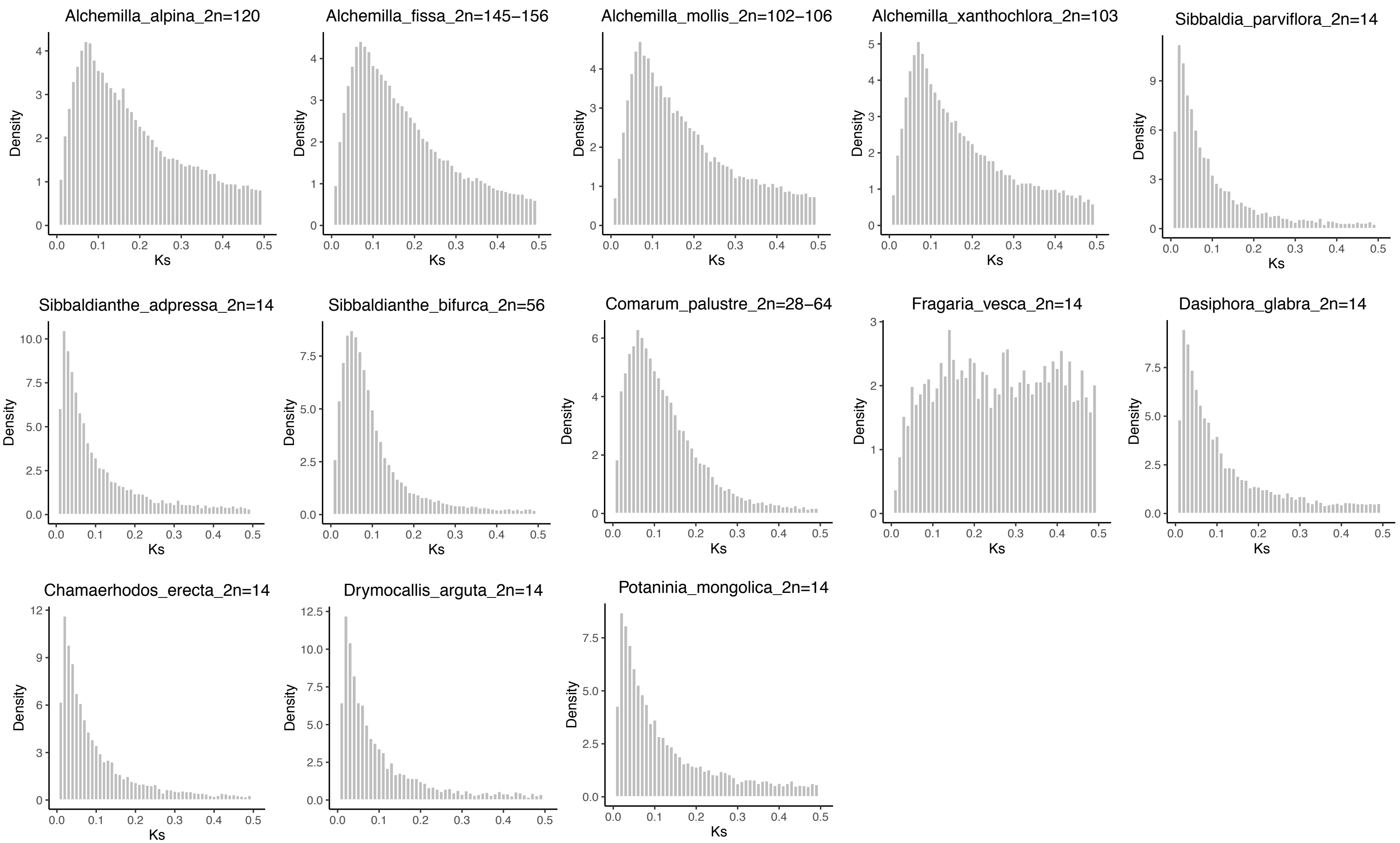
Alchemilla_alpina_2n=120
Alchemilla_fissa_2n=145−156
Alchemilla_mollis_2n=102−106
Alchemilla_xanthochlora_2n=103
Sibbaldia_parviflora_2n=14
Sibbaldianthe_adpressa_2n=14
Sibbaldianthe_bifurca_2n=56
Comarum_palustre_2n=28−64
Fragaria_vesca_2n=14
Dasiphora_glabra_2n=14
Chamaerhodos_erecta_2n=14
Drymocallis_arguta_2n=14
Potaninia_mongolica_2n=14
Density
Ks

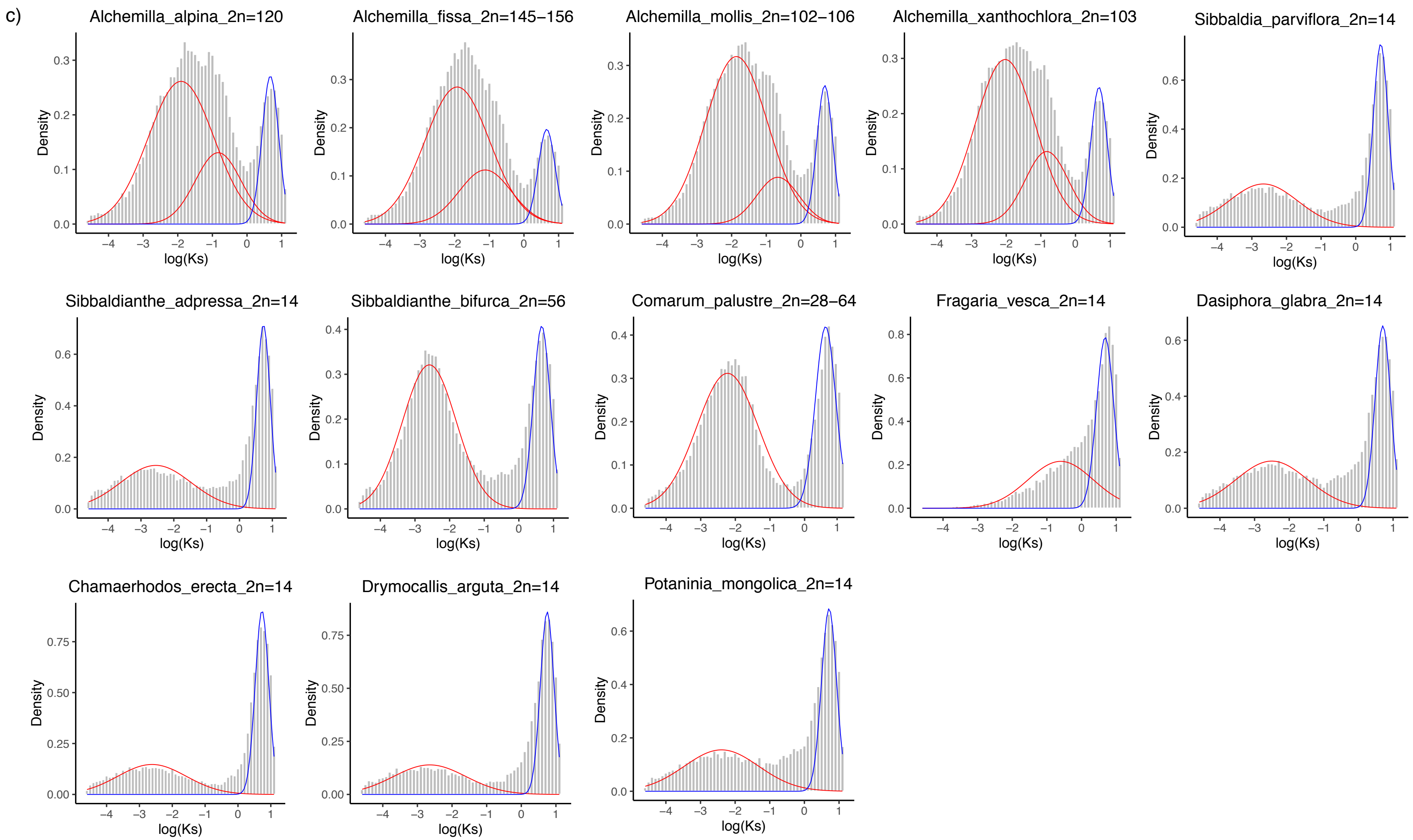


**Figure S11**. Distribution density of synonymous distance among gene pairs (Ks plots) for four species of Eualchemilla and nine species representing other genera in Fragariinae. *Alchemilla alpina* belongs to the 'dissected' clade, and *A. fissa*, *A. mollis*, and *A. xanthochlora* belong to the ‘lobed’ clade. Reported chromosome numbers for each species are noted next to the species name. a) Plots of raw Ks distances between 0 and 3. b) Plots of Ks distances between 0 and 0.5. c) Plots of log-transformed Ks distances. Colored lines denote components inferred using a mixture model. Red lines are unique components for each species. Blue lines denote a component from an ancestral whole genome triplication event in the core eudicots.

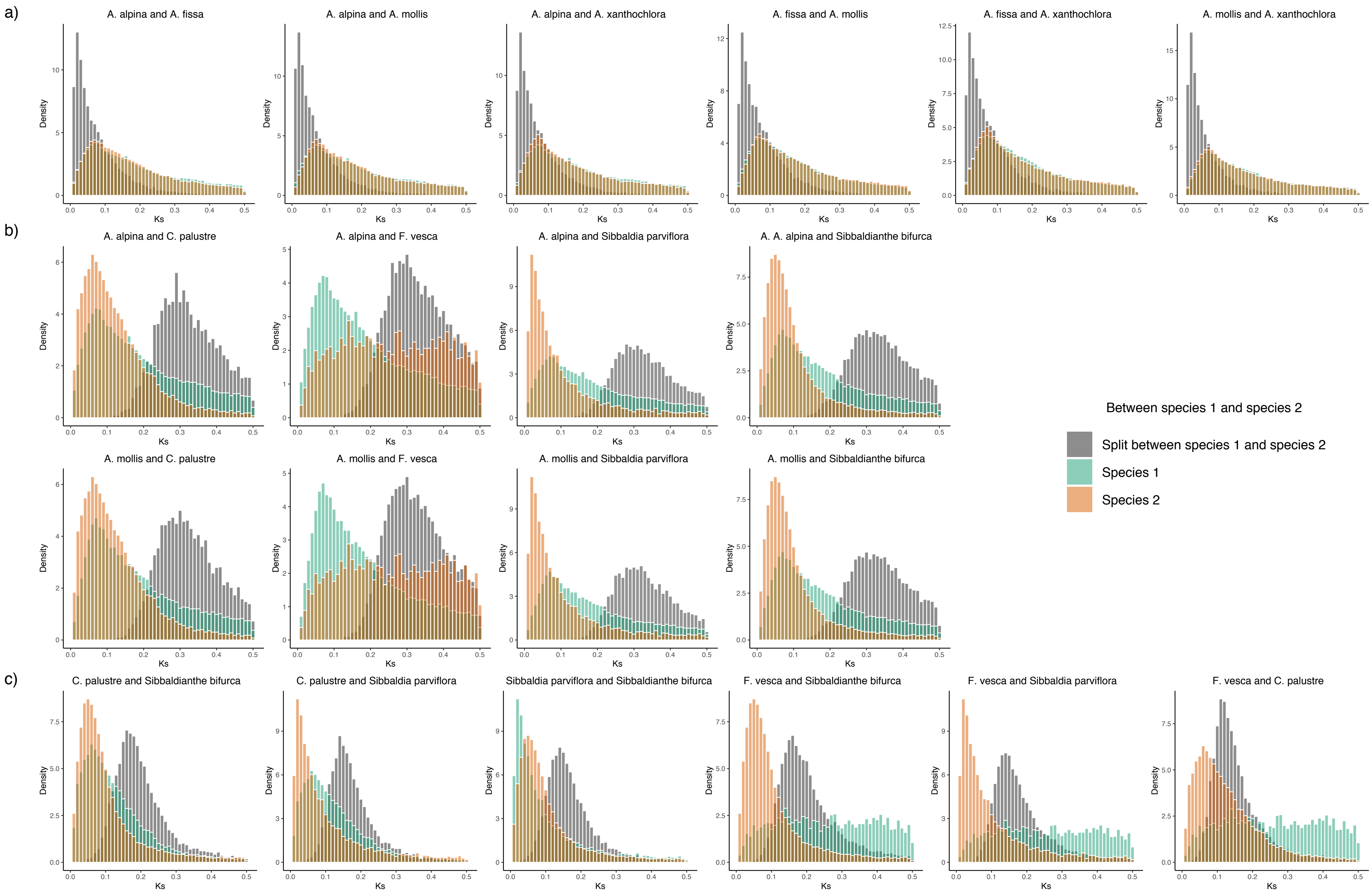


**Figure S12**. Distribution density of synonymous distance among gene pairs (Ks plots). a) Between species pairs in Eualchemilla. *Alchemilla alpina* belongs to the 'dissected' clade, and *A. fissa*, *A. mollis*, and *A. xanthochlora* belong to the 'lobed' clade. b) Top row: between *A. alpina* versus members of Fragariinae outside of *Alchemilla* s.l. Bottom row: *A. mollis* versus members of Fragariinae outside of *Alchemilla* s.l. c) Between members of Fragariinae outside of *Alchemilla* s.l.

**Table S1.** Main characteristics of majors clades of *Alchemilla* s.l.

| Characters | Eualchemilla | Afromilla | *Lachemilla* | *Aphanes* |
|---|---|---|---|---|
| Life cycle | Perennial | Perennial | Perennial | Annual or biennial |
| Habit | Rosette-forming herbs, prostrate | Rosette-forming herbs, trailing | Rosette-forming herbs, trailing | Slender ascending herbs |
| Leaf morphology | Lobed to dissected | Lobed to dissected | Lobed, dissected, pinnate, bi- | Dissected |
| Number of species | at least ca. 200 (–900) | ca. 70 | ca. 65 | ca. 20 |
| Distribution | Holarctic, with a center of | Sub-Saharan and Madagascar, | Mountains of the Neotropics, | Temperate regions of the world, |
| Stamens | Four stamens inserted on the | Four stamens inserted on the outer | Two stamens inserted on the | One stamens inserted on the inner |
| Ploidy ($x = 8$) | Octoploid to 28-ploid | Octoploid and decaploid (data | Diploid to dodecaploid (mostly | Diploid, tetraploid, and hexaploid |
| Apomixis | Autonomous | Present (mechanism unknown) | Autonomous | Autonomous |

**Table S2.** List of species and vouchers used in this study.

| Species | Collection number | Herbarium | Major group | Section/clade within | Source | SRA accesion number | Notes |
|---|---|---|---|---|---|---|---|
| *Alchemilla johnstonii* Oliv. | B. Gehrke 364 | ZT | Afromilla | Sect. Geraniifoliae | This study | SRR12516439 | |
| *Alchemilla roccatii* Cort. | B. Gehrke 365 | ZT | Afromilla | Sect. Geraniifoliae | This study | SRR12516432 | |
| *Alchemilla haumanii* Rothm. | B. Gehrke 204 | ZT | Afromilla | Sect. Grandifoliae | This study | SRR12516444 | |
| *Alchemilla colura* Hilliard | B. Gehrke 464 | ZT | Afromilla | Sect. Longicaules | This study | SRR12516445 | |
| *Alchemilla cryptantha* Steud. ex A.Rich. | B. Gehrke 306 | ZT | Afromilla | Sect. Longicaules | This study | SRR12516434 | |
| *Alchemilla ellenbeckii* Engl. | B. Gehrke 104 | ZT | Afromilla | Sect. Longicaules | This study | SRR12516401 | |
| *Alchemilla elongata* Eckl. & Zeyh. | B. Gehrke 446 | ZT | Afromilla | Sect. Longicaules | This study | SRR12516390 | |
| *Alchemilla fischeri* Engl. | B. Gehrke 018 | ZT | Afromilla | Sect. Longicaules | This study | SRR12516415 | |
| *Alchemilla kiwuensis* Engl. | B. Gehrke 057 | ZT | Afromilla | Sect. Longicuales | This study | SRR12516438 | |
| *Alchemilla woodii* Kuntze | B. Gehrke 453 | ZT | Afromilla | Sect. Longicuales | This study | SRR12516424 | |
| *Alchemilla argyrophylla* Oliv. | W. Rauh T139 | M | Afromilla | Sect. Parvifoliae | Morales-Briones et al. 2018b | SRR6667468 | |
| *Alchemilla microbetula* T.C.E.Fr. | B. Gehrke 360 | ZT | Afromilla | Sect. Parvifoliae | This study | SRR12516436 | |
| *Alchemilla hildebrandtii* Engl. | B. Gehrke 258 | ZT | Afromilla | Sect. Schizophyllae | This study | SRR12516442 | |
| *Alchemilla rutenbergii* O.Hoffm. | B. Gehrke 253 | ZT | Afromilla | Sect. Schizophyllae | This study | SRR12516431 | |
| *Alchemilla schizophylla* Bak. | B. Gehrke 282 | ZT | Afromilla | Sect. Schizophyllae | This study | SRR12516430 | |
| *Alchemilla stuhlmannii* Engl. | B. Gehrke 363 | ZT | Afromilla | Sect. Subcuneatifoliae | This study | SRR12516428 | |
| *Alchemilla subnivalis* Bak. | B. Gehrke 362 | ZT | Afromilla | Sect. Subcuneatifoliae | This study | SRR12516427 | |
| *Alchemilla triphylla* Rothm. | B. Gehrke 361 | ZT | Afromilla | Sect. Subcuneatifoliae | This study | SRR12516425 | |
| *Aphanes arvensis* L. | Bot. Garden Zürich s.n. | ZT | *Aphanes* | Sect. Inaequidentatae | This study | SRR12516410 | |
| *Aphanes australis* Rydb. | D.C. Tank 1107 | ID | *Aphanes* | Sect. Inaequidentatae | Morales-Briones et al. 2018b | SRR6667469 | |
| *Aphanes cotopaxiensis* Romol. & Frost-Olsen | D.F. Morales-Briones et al. 276 | ID | *Aphanes* | Sect. Inaequidentatae | This study | SRR12516409 | |
| *Aphanes microcarpa* (Boiss. & Reut.) Rothm. | L. Koepke 201000524-1 | OSU | *Aphanes* | Sect. Inaequidentatae | This study | SRR12516408 | |
| *Aphanes occidentalis* (Nutt.) Rydb. | D.F. Morales-Briones & D. Caetano 579 | ID | *Aphanes* | Sect. Inaequidentatae | This study | SRR12516407 | |
| *Aphanes* sp. "boliviana" | D.F. Morales-Briones et al. 291 | ID | *Aphanes* | Sect. Inaequidentatae | This study | SRR12516406 | Aphanes sp[a] |
| *Aphanes* sp. "patagonica" | Marticoerena et al. 941 | CONC | *Aphanes* | Sect. Inaequidentatae | This study | SRR12516405 | Aphanes neglecta[a] |
| *Alchemilla alpina* L. | SL 11498 | ZT | Eualchemilla - Dissected clade | Sect. Alpinae | This study | SRR12516446 | |
| *Alchemilla subsericea* Reut. | P. Frost-Olsen 11984 | AAU | Eualchemilla - Dissected clade | Sect. Glaciales | This study | SRR12516426 | |
| *Alchemilla pentaphyllea* L. | J. Kadereit & M. Lauterbach 201252 | JGU | Eualchemilla - Dissected clade | Sect. Pentaphylleae | This study | SRR12516435 | |
| *Alchemilla xanthochlora* Rothm. | Lippert s.n. | M | Eualchemilla - Lobed clade | Sect. Alchemilla | This study | SRR12516411 | |
| *Alchemilla fissa* Gunth. & Schumm. | P. Frost-Olsen 8240 | AAU | Eualchemilla - Lobed clade | Sect. Calycinae | This study | SRR12516414 | |
| *Alchemilla glabra* Neygenf. | P. Frost-Olsen 12658 | AAU | Eualchemilla - Lobed clade | Sect. Coriaceae | This study | SRR12516412 | |
| *Alchemilla decumbens* Buser | B. Gehrke 662 | ZT | Eualchemilla - Lobed clade | Sect. Decumbentes | This study | SRR12516423 | |
| *Alchemilla indivisa* Formánek ex Rothm. | P. Frost-Olsen 3383 | AAU | Eualchemilla - Lobed clade | Sect. Erectae | This study | SRR12516441 | |
| *Alchemilla mollis* (Buser) Rothm. | D.F. Morales-Briones 687 | ID | Eualchemilla - Lobed clade | Sect. Erectae | Morales-Briones et al. 2018b | SRR6667455 | |
| *Alchemilla flabellata* Buser | P. Frost-Olsen 11859 | AAU | Eualchemilla - Lobed clade | Sect. Flabellatae | This study | SRR12516413 | |
| *Alchemilla japonica* Nakai & H.Hara | B. Dichore 1986 | GOET | Eualchemilla - Lobed clade | Sect. Plicatae | This study | SRR12516440 | |
| *Alchemilla plicata* Buser | P. Frost-Olsen 11575 | AAU | Eualchemilla - Lobed clade | Sect. Plicatae | This study | SRR12516433 | |
| *Alchemilla lapeyrousii* Buser | P. Frost-Olsen 10029 | AAU | Eualchemilla - Lobed clade | Sect. Pubescentes | This study | SRR12516437 | |
| *Alchemilla splendens* Christ | P. Frost-Olsen 7587 | AAU | Eualchemilla - Lobed clade | Sect. Splendentes | This study | SRR12516429 | |
| *Alchemilla heptagona* Juz. | P. Frost-Olsen 6999 | AAU | Eualchemilla - Lobed clade | Sect. Ultravulgares | This study | SRR12516443 | |
| *Lachemilla orbiculata* (Ruiz & Pav.) Rydb. | D.F. Morales-Briones et al. 108 | QCA | *Lachemilla* | Orbiculate | Morales-Briones et al. 2018b | SRR6667480 | |
| *Lachemilla pectinata* (Kunth) Rothm. | D.F. Morales-Briones & K. Romoleroux 161 | QCA | *Lachemilla* | Orbiculate | Morales-Briones et al. 2018b | SRR6667481 | |
| *Lachemilla barbata* (C.Presl) Rothm. | D.F. Morales-Briones & S. Uribe-Convers 229 | ID | *Lachemilla* | Pinnate | Morales-Briones et al. 2018b | SRR6667472 | |
| *Lachemilla diplophylla* (Diels) Rothm. | D.F. Morales-Briones & E. Morales-Checa 30 | QCA | *Lachemilla* | Pinnate | Morales-Briones et al. 2018b | SRR6667473 | |
| *Lachemilla erodiifolia* (Wedd.) Rothm. | D.F. Morales-Briones & K. Romoleroux 119 | QCA | *Lachemilla* | Pinnate | Morales-Briones et al. 2018b | SRR6667470 | |
| *Lachemilla mandoniana* (Wedd.) Rothm. | D.F. Morales-Briones et al. 112 | QCA | *Lachemilla* | Pinnate | Morales-Briones et al. 2018b | SRR6667482 | |
| *Lachemilla pinnata* (Ruiz & Pav.) Rothm. | D.F. Morales-Briones et al. 113 | QCA | *Lachemilla* | Pinnate | Morales-Briones et al. 2018b | SRR6667484 | |
| *Lachemilla tanacetifolia* Rothm. | D.F. Morales-Briones et al. 122 | QCA | *Lachemilla* | Pinnate | Morales-Briones et al. 2018b | SRR6667464 | |
| *Lachemilla andina* (L.M.Perry) Rothm. | D.F. Morales-Briones & K. Romoleroux 162 | QCA | *Lachemilla* | Tripartite | Morales-Briones et al. 2018b | SRR6667466 | |
| *Lachemilla aphanoides* (Mutis ex L.f.) Rothm. | D.F. Morales-Briones et al. 115 | QCA | *Lachemilla* | Tripartite | Morales-Briones et al. 2018b | SRR6667467 | |
| *Lachemilla hirta* (L.M.Perry) Rothm. | D.F. Morales-Briones & K. Romoleroux 118 | QCA | *Lachemilla* | Tripartite | Morales-Briones et al. 2018b | SRR6667475 | |
| *Lachemilla jamesonii* (L.M.Perry) Rothm. | K. Romoleroux et al. 4684 | QCA | *Lachemilla* | Tripartite | Morales-Briones et al. 2018b | SRR6667476 | |
| *Lachemilla moritziana* Dammer | D.F. Morales-Briones & S. Uribe-Convers 427 | ID | *Lachemilla* | Tripartite | This study | SRR12516394 | |
| *Lachemilla pringlei* Rydb. | D.F. Morales-Briones & P. Tenorio-Lezama 595 | ID | *Lachemilla* | Tripartite | This study | SRR12516393 | |
| *Lachemilla procumbens* (Rose) Rydb. | K. Romoleroux et al. 5011 | QCA | *Lachemilla* | Tripartite | Morales-Briones et al. 2018b | SRR6667461 | |
| *Lachemilla sibbaldiifolia* (Kunth) Rydb. | D.F. Morales-Briones & P. Tenorio-Lezama 599 | ID | *Lachemilla* | Tripartite | This study | SRR12516391 | |
| *Lachemilla standleyi* (L.M.Perry) Rothm. | K. Romoleroux et al. 5005 | QCA | *Lachemilla* | Tripartite | This study | SRR12516389 | |
| *Lachemilla velutina* (S.Watson) Rydb. | D.F. Morales-Briones & P. Tenorio-Lezama 607 | ID | *Lachemilla* | Tripartite | This study | SRR12516388 | |
| *Lachemilla vulcanica* (Schltdl. & Cham.) Rydb. | D.F. Morales-Briones et al. 117 | QCA | *Lachemilla* | Tripartite | Morales-Briones et al. 2018b | SRR6667486 | |

| | | | | | | |
|---|---|---|---|---|---|---|
| *Lachemilla angustata* Romol. | K. Romoleroux et al.1562 | QCA | *Lachemilla* | Verticillate | This study | SRR12516397 |
| *Lachemilla equisetiformis* (Trevir.) Rothm. | Muller 91-28 | MSB | *Lachemilla* | Verticillate | This study | SRR12516396 |
| *Lachemilla galioides* (Benth.) Rothm. | K. Romoleroux et al. 4699 | QCA | *Lachemilla* | Verticillate | Morales-Briones et al. 2018b | SRR6667474 |
| *Lachemilla hispidula* (L.M.Perry) Rothm. | D.F. Morales-Briones & K. Romoleroux 110 | QCA | *Lachemilla* | Verticillate | Morales-Briones et al. 2018b | SRR6667478 |
| *Lachemilla llanganatensis* Romol. | D.F. Morales-Briones et al. 63 | QCA | *Lachemilla* | Verticillate | This study | SRR12516395 |
| *Lachemilla nivalis* (Kunth) Rothm. | K. Romoleroux et al. 4580 | QCA | *Lachemilla* | Verticillate | Morales-Briones et al. 2018b | SRR6667483 |
| *Lachemilla polylepis* (Wedd.) Rothm. | P. Sklenář 12207 | QCA | *Lachemilla* | Verticillate | Morales-Briones et al. 2018b | SRR6667485 |
| *Lachemilla ramosissima* (Rothm.) Rothm. | Aristeguieta 3882 | NY | *Lachemilla* | Verticillate | This study | SRR12516392 |
| *Lachemilla verticillata* (Fielding & Gardner) Rothm. | K. Romoleroux et al. 5007 | QCA | *Lachemilla* | Verticillate | Morales-Briones et al. 2018b | SRR6667457 |
| *Chamaerhodos erecta* (L.) Bunge | R.L. Hartman & B.E Nelson 54641 | ID | Fragariinae | N/A | This study | SRR12516404 |
| *Comarum palustre* L. | J. Duemmel et al. 29-08 | ID | Fragariinae | N/A | This study | SRR12516403 |
| *Dasiphora fruticosa* (L.) Rydb. | H.E. Marx & D. Daly 2013-209 | ID | Fragariinae | N/A | This study | SRR12516402 |
| *Drymocallis arguta* (Pursh) Rydb. | J.F. Smith 8736 | ID | Fragariinae | N/A | This study | SRR12516400 |
| *Drymocallis glandulosa* (Lindl.) Rydb. | A. Liston 1373 | OSC | Fragariinae | N/A | This study | SRR12516399 |
| *Farinopsis salesoviana* (Stephan) Chrtek & Soják | B. Bartholomew et al. 8364 | MO | Fragariinae | N/A | This study | SRR12516398 |
| *Fragaria vesca* L. | N/A | N/A | Fragariinae | N/A | Shulaev, et. al 2010 | Genome v1.1/Phytozome |
| *Potaninia mongolica* Maxim. | L. Yingxin & Z. Xiufu 93039 | MO | Fragariinae | N/A | This study | SRR12516422 |
| *Sibbaldia procumbens* L. | H.E. Marx & Z. Klos 2012-041 | ID | Fragariinae | N/A | This study | SRR12516418 |
| *Sibbaldianthe adpressa* (Bunge) Juz. | И.М. Краснобоpов 320 | MO | Fragariinae | N/A | This study | SRR12516417 |
| *Sibbaldianthe bifurca* (L.) Kurtto & T.Erikss. | D.E. Boufford et al. 29796 | MO | Fragariinae | N/A | This study | SRR12516416 |
| *Potentilla indica* (Andrews) Th.Wolf | A. Liston 1363 | OSC | Outgroup | N/A | This study | SRR12516421 |
| *Rosa woodsii* Lindl. | A. Liston 1360 | OSC | Outgroup | N/A | This study | SRR12516420 |
| *Sanguisorba menziesii* Rydb. | A. Liston 1342 | OSC | Outgroup | N/A | This study | SRR12516419 |

[a]Name in alignment and tree files

**Table S3.** Plastomes used for assembly and phylogenetic inference.

| Species | GenBank accession | Reference |
|---|---|---|
| *Chamaerhodos erecta*[a] | KY420001 | Zhang et al., 2017 |
| *Dasiphora fruticosa*[a] | KY420016 | Zhang et al., 2017 |
| *Drymocallis glandulosa*[a] | KY420015 | Zhang et al., 2017 |
| *Farinopsis salesoviana*[a] | KY420034 | Zhang et al., 2017 |
| *Fragaria vesca*[a] | JF345175 | Shulaev et al., 2011 |
| *Lachemilla pectinata*[b] | KY419937 | Zhang et al., 2017 |
| *Potaninia mongolica*[a] | KY419959 | Zhang et al., 2017 |
| *Potentilla indica*[a] | KY420014 | Zhang et al., 2017 |
| *Rosa lichiangensis*[b] | KY419934 | Zhang et al., 2017 |
| *Sanguisorba officinalis*[b] | KY419975 | Zhang et al., 2017 |
| *Sibbaldia procumbens*[a] | KY419935 | Zhang et al., 2017 |
| *Sibbaldianthe sericea*[b] | KY419993 | Zhang et al., 2017 |

[a]Used for plastome assembly and phylogenetic inference.

[b]Used only for plastome assembly.

**Tables S4.** List of species used for synonymous distance among gene pairs (Ks) plots.

| Species | Ploidy ($x$ = 8 in | Major group | Section/clade | Source | Accesion number |
|---|---|---|---|---|---|
| *Alchemilla alpina* | 2$n$ = 120 | Eualchemilla | Dissected clade | Xiang et al. 2017 | SRR13526600 |
| *Alchemilla fissa* | 2$n$ = 145–156 | Eualchemilla | Lobed clade | Xiang et al. 2017 | SRR13526599 |
| *Alchemilla mollis* | 2$n$ = 102–106 | Eualchemilla | Lobed clade | Unpublished | SRR11487710 |
| *Alchemilla xanthocholora* | 2$n$ = 103 | Eualchemilla | Lobed clade | Unpublished | SRR11487709 |
| *Chamaerhodos erecta* | 2$n$ = 14 | Fragariinae | N/A | Xiang et al. 2017 | SRR13526598 |
| *Comarum palustre* | 2$n$ = 28–64 | Fragariinae | N/A | Xiang et al. 2017 | SRR13526597 |
| *Dasiphora glabra* | 2$n$ = 14 | Fragariinae | N/A | Xiang et al. 2017 | SRR13526596 |
| *Drymocallis arguta* | 2$n$ = 14 | Fragariinae | N/A | Xiang et al. 2017 | SRR13526595 |
| *Fragaria vesca* | 2$n$ = 14 | Fragariinae | N/A | Shulaev et. al 2010 | Phytozome - Genome v1.1 |
| *Potaninia mongolica* | 2$n$ = 14 | Fragariinae | N/A | Xiang et al. 2017 | SRR13526594 |
| *Sibbaldia parviflora* | 2$n$ = 14 | Fragariinae | N/A | Xiang et al. 2017 | SRR13526593 |
| *Sibbaldianthe bifurca* | 2$n$ = 56 | Fragariinae | N/A | Xiang et al. 2017 | SRR13526592 |
| *Sibbaldianthe adpressa* | 2$n$ = 14 | Fragariinae | N/A | Xiang et al. 2017 | SRR13526591 |

**Table S5.** HybPiper assembly statistics

| Species | Number of reads | Number of | Percent reads | Number of exons | Number of exons | Number of exons | Number of exons | Number of exons | Number of exons | Number of exons | Number of exons |
|---|---|---|---|---|---|---|---|---|---|---|---|
| *Alchemilla_alpina* | 524,481 | 358,784 | 68.4% | 926 | 862 | 860 | 857 | 849 | 821 | 3 | 370 |
| *Alchemilla_argyrophylla* | 9,423,799 | 2,302,455 | 24.4% | 930 | 909 | 906 | 905 | 889 | 863 | 0 | 686 |
| *Alchemilla_colura* | 2,339,559 | 1,659,277 | 70.9% | 930 | 919 | 916 | 916 | 909 | 895 | 0 | 623 |
| *Alchemilla_cryptantha* | 4,234,959 | 3,104,915 | 73.3% | 930 | 927 | 924 | 924 | 915 | 891 | 2 | 671 |
| *Alchemilla_decumbens* | 1,718,446 | 1,080,216 | 62.9% | 932 | 914 | 909 | 907 | 895 | 864 | 2 | 572 |
| *Alchemilla_ellenbeckii* | 2,234,479 | 1,633,354 | 73.1% | 933 | 923 | 919 | 918 | 912 | 890 | 0 | 662 |
| *Alchemilla_elongata* | 3,384,889 | 2,452,689 | 72.5% | 931 | 924 | 921 | 919 | 911 | 888 | 2 | 655 |
| *Alchemilla_fischeri* | 2,371,539 | 1,681,160 | 70.9% | 932 | 925 | 921 | 919 | 911 | 891 | 1 | 654 |
| *Alchemilla_fissa* | 168,831 | 108,950 | 64.5% | 923 | 695 | 692 | 688 | 671 | 632 | 0 | 66 |
| *Alchemilla_flabellata* | 743,681 | 473,981 | 63.7% | 923 | 887 | 881 | 880 | 865 | 834 | 2 | 458 |
| *Alchemilla_glabra* | 2,295,784 | 1,556,467 | 67.8% | 929 | 921 | 915 | 914 | 904 | 880 | 1 | 579 |
| *Alchemilla_haumanii* | 2,234,226 | 1,608,164 | 72.0% | 927 | 920 | 915 | 914 | 908 | 888 | 1 | 617 |
| *Alchemilla_heptagona* | 1,856,644 | 1,210,799 | 65.2% | 925 | 918 | 912 | 910 | 905 | 881 | 1 | 585 |
| *Alchemilla_hildebrandtii* | 484,180 | 326,667 | 67.5% | 928 | 870 | 867 | 865 | 856 | 826 | 1 | 470 |
| *Alchemilla_indivisa* | 1,734,141 | 1,107,718 | 63.9% | 929 | 915 | 910 | 909 | 902 | 876 | 1 | 575 |
| *Alchemilla_japonica* | 2,217,591 | 1,514,768 | 68.3% | 930 | 921 | 916 | 915 | 909 | 882 | 0 | 583 |
| *Alchemilla_johnstonii* | 3,378,617 | 2,400,151 | 71.0% | 930 | 925 | 924 | 921 | 915 | 899 | 0 | 645 |
| *Alchemilla_kiwuensis* | 1,709,266 | 1,186,794 | 69.4% | 929 | 912 | 909 | 906 | 902 | 886 | 1 | 601 |
| *Alchemilla_lapeyrousii* | 1,096,905 | 691,343 | 63.0% | 933 | 895 | 891 | 887 | 874 | 848 | 2 | 477 |
| *Alchemilla_microbetula* | 4,323,618 | 3,122,993 | 72.2% | 933 | 927 | 923 | 923 | 918 | 897 | 0 | 642 |
| *Alchemilla_mollis* | 10,805,273 | 4,148,516 | 38.4% | 930 | 919 | 915 | 911 | 902 | 886 | 1 | 746 |
| *Alchemilla_pentaphyllea* | 1,931,279 | 1,257,408 | 65.1% | 929 | 918 | 916 | 916 | 908 | 876 | 4 | 619 |
| *Alchemilla_plicata* | 473,046 | 309,663 | 65.5% | 929 | 860 | 856 | 855 | 845 | 818 | 0 | 384 |
| *Alchemilla_roccatii* | 3,508,263 | 2,523,772 | 71.9% | 933 | 927 | 924 | 924 | 917 | 895 | 0 | 630 |
| *Alchemilla_rutenbergii* | 1,781,618 | 1,268,216 | 71.2% | 932 | 921 | 919 | 919 | 912 | 891 | 1 | 595 |
| *Alchemilla_schizophylla* | 770,984 | 530,610 | 68.8% | 930 | 899 | 898 | 896 | 885 | 855 | 1 | 541 |
| *Alchemilla_splendens* | 881,924 | 569,639 | 64.6% | 928 | 891 | 890 | 884 | 869 | 835 | 4 | 437 |
| *Alchemilla_stuhlmannii* | 501,136 | 335,671 | 67.0% | 926 | 888 | 887 | 885 | 869 | 846 | 1 | 428 |
| *Alchemilla_subnivalis* | 1,852,807 | 1,225,895 | 66.2% | 929 | 923 | 920 | 920 | 910 | 879 | 0 | 581 |
| *Alchemilla_subsericea* | 2,400,286 | 1,636,676 | 68.2% | 929 | 921 | 918 | 918 | 909 | 882 | 2 | 579 |
| *Alchemilla_triphylla* | 573,770 | 374,779 | 65.3% | 928 | 883 | 882 | 880 | 871 | 846 | 1 | 428 |
| *Alchemilla_woodii* | 657,131 | 444,252 | 67.6% | 930 | 887 | 886 | 882 | 868 | 842 | 0 | 497 |
| *Alchemilla_xanthochlora* | 952,341 | 671,634 | 70.5% | 929 | 914 | 911 | 908 | 899 | 871 | 0 | 541 |
| *Aphanes_arvensis* | 2,637,012 | 1,690,388 | 64.1% | 900 | 875 | 871 | 864 | 853 | 821 | 0 | 256 |
| *Aphanes_australis* | 15,601,983 | 3,760,913 | 24.1% | 902 | 848 | 843 | 840 | 833 | 813 | 0 | 152 |
| *Aphanes_cotopaxiensis* | 402,741 | 232,904 | 57.8% | 892 | 825 | 821 | 818 | 804 | 780 | 0 | 124 |
| *Aphanes_microcarpa* | 467,018 | 277,304 | 59.4% | 905 | 830 | 825 | 823 | 810 | 786 | 0 | 120 |
| *Aphanes_occidentalis* | 1,643,973 | 1,081,203 | 65.8% | 884 | 859 | 851 | 847 | 838 | 814 | 0 | 133 |
| *Aphanes_sp "boliviana"* | 651,401 | 390,319 | 59.9% | 897 | 838 | 834 | 831 | 819 | 800 | 0 | 128 |
| *Aphanes_sp "patagonica"* | 3,576,077 | 2,484,557 | 69.5% | 902 | 867 | 860 | 854 | 846 | 823 | 0 | 143 |
| *Chamaerhodos_erecta* | 2,384,006 | 1,808,610 | 75.9% | 938 | 926 | 926 | 926 | 925 | 922 | 0 | 29 |
| *Comarum_paluste* | 1,436,708 | 1,173,401 | 81.7% | 938 | 937 | 937 | 936 | 936 | 932 | 0 | 388 |
| *Dasiphora_fruticosa* | 965,137 | 743,804 | 77.1% | 939 | 938 | 938 | 938 | 938 | 934 | 0 | 48 |
| *Drymocallis_arguta* | 3,183,973 | 2,585,201 | 81.2% | 939 | 939 | 939 | 939 | 939 | 934 | 0 | 32 |
| *Drymocallis_glandulosa* | 1,805,073 | 562,801 | 31.2% | 939 | 927 | 927 | 927 | 921 | 900 | 0 | 10 |
| *Farinopsis_salesoviana* | 2,831,516 | 2,428,596 | 85.8% | 939 | 939 | 939 | 937 | 927 | 906 | 0 | 429 |
| *Lachemilla_andina* | 11,120,534 | 4,403,408 | 39.6% | 929 | 919 | 915 | 915 | 909 | 892 | 0 | 628 |
| *Lachemilla_angustata* | 2,107,992 | 1,682,792 | 79.8% | 931 | 919 | 915 | 911 | 899 | 873 | 1 | 475 |
| *Lachemilla_aphanoides* | 18,547,039 | 7,933,424 | 42.8% | 933 | 921 | 913 | 913 | 911 | 903 | 2 | 654 |
| *Lachemilla_barbata* | 23,347,654 | 9,207,955 | 39.4% | 932 | 923 | 917 | 917 | 910 | 904 | 1 | 653 |
| *Lachemilla_diplophylla* | 13,925,112 | 5,978,669 | 42.9% | 930 | 917 | 914 | 913 | 907 | 898 | 0 | 638 |
| *Lachemilla_equisetiformis* | 455,469 | 343,612 | 75.4% | 929 | 883 | 881 | 877 | 860 | 822 | 0 | 302 |

| | | | | | | | | | | | |
|---|---|---|---|---|---|---|---|---|---|---|---|
| *Lachemilla_erodiifolia* | 8,525,733 | 3,441,532 | 40.4% | 928 | 915 | 912 | 910 | 906 | 896 | 1 | 645 |
| *Lachemilla_galioides* | 8,234,295 | 3,333,021 | 40.5% | 930 | 908 | 904 | 904 | 899 | 888 | 1 | 651 |
| *Lachemilla_hirta* | 14,748,363 | 6,391,178 | 43.3% | 930 | 918 | 915 | 915 | 910 | 900 | 3 | 625 |
| *Lachemilla_hispidula* | 8,337,125 | 3,367,380 | 40.4% | 928 | 914 | 912 | 912 | 906 | 895 | 1 | 619 |
| *Lachemilla_jamesonii* | 10,935,704 | 4,421,523 | 40.4% | 931 | 917 | 914 | 914 | 909 | 892 | 2 | 644 |
| *Lachemilla_llanganatensis* | 529,599 | 344,731 | 65.1% | 928 | 887 | 883 | 882 | 878 | 858 | 1 | 409 |
| *Lachemilla_mandoniana* | 7,238,811 | 2,537,702 | 35.1% | 931 | 908 | 906 | 906 | 902 | 888 | 1 | 624 |
| *Lachemilla_moritziana* | 501,013 | 319,142 | 63.7% | 933 | 880 | 878 | 877 | 871 | 851 | 1 | 336 |
| *Lachemilla_nivalis* | 7,912,741 | 2,643,787 | 33.4% | 925 | 905 | 902 | 902 | 897 | 888 | 1 | 604 |
| *Lachemilla_orbiculata* | 9,159,390 | 3,519,420 | 38.4% | 927 | 913 | 910 | 909 | 905 | 893 | 0 | 625 |
| *Lachemilla_pectinata* | 13,747,010 | 5,885,754 | 42.8% | 930 | 918 | 917 | 917 | 914 | 900 | 1 | 692 |
| *Lachemilla_pinnata* | 21,402,835 | 7,412,832 | 34.6% | 930 | 921 | 916 | 916 | 912 | 901 | 0 | 641 |
| *Lachemilla_polylepis* | 14,256,485 | 4,708,023 | 33.0% | 929 | 917 | 915 | 915 | 909 | 901 | 1 | 637 |
| *Lachemilla_pringlei* | 613,473 | 418,391 | 68.2% | 928 | 889 | 887 | 886 | 881 | 865 | 0 | 410 |
| *Lachemilla_procumbens* | 8,827,483 | 3,277,780 | 37.1% | 928 | 910 | 906 | 906 | 902 | 889 | 0 | 618 |
| *Lachemilla_ramosissima* | 1,629,249 | 1,313,895 | 80.6% | 934 | 919 | 914 | 909 | 899 | 873 | 1 | 460 |
| *Lachemilla_sibbaldiifolia* | 645,613 | 445,494 | 69.0% | 929 | 902 | 898 | 898 | 889 | 874 | 1 | 406 |
| *Lachemilla_standleyi* | 3,039,388 | 2,329,689 | 76.6% | 929 | 926 | 921 | 919 | 908 | 890 | 1 | 501 |
| *Lachemilla_tanacetifolia* | 5,333,257 | 2,323,802 | 43.6% | 929 | 908 | 905 | 905 | 897 | 882 | 0 | 647 |
| *Lachemilla_velutina* | 655,979 | 453,092 | 69.1% | 925 | 901 | 896 | 896 | 891 | 873 | 0 | 445 |
| *Lachemilla_verticillata* | 9,252,744 | 4,020,526 | 43.5% | 932 | 916 | 913 | 912 | 906 | 897 | 1 | 652 |
| *Lachemilla_vulcanica* | 9,285,254 | 4,015,561 | 43.2% | 932 | 915 | 911 | 911 | 907 | 892 | 1 | 641 |
| *Potaninia_mongolica* | 2,828,344 | 2,419,418 | 85.5% | 938 | 933 | 932 | 932 | 930 | 923 | 0 | 63 |
| *Potentilla_indica* | 4,917,501 | 1,534,558 | 31.2% | 939 | 925 | 925 | 925 | 921 | 890 | 1 | 96 |
| *Rosa_woodsii* | 4,510,487 | 1,057,848 | 23.5% | 939 | 933 | 933 | 933 | 928 | 902 | 0 | 28 |
| *Sanguisorba_menziesii* | 3,033,042 | 648,547 | 21.4% | 939 | 901 | 900 | 899 | 882 | 844 | 1 | 122 |
| *Sibbaldia_procumbens* | 538,539 | 426,339 | 79.2% | 936 | 933 | 933 | 933 | 933 | 931 | 0 | 32 |
| *Sibbaldianthe_adpressa* | 1,455,531 | 1,269,404 | 87.2% | 936 | 935 | 934 | 934 | 931 | 921 | 1 | 30 |
| *Sibbaldianthe_bifurca* | 2,801,305 | 2,416,512 | 86.3% | 936 | 936 | 936 | 936 | 935 | 923 | 0 | 117 |

**Table S6.** MO and RT orthologs statistics.

| Species | Number of MO orthologs | Percentage of MO orthologs (total 914) | Number of RT orthologs | Percentage of RT orthologs (total 1906) |
|---|---|---|---|---|
| *Alchemilla_alpina* | 396 | 43.3% | 712 | 37.4% |
| *Alchemilla_argyrophylla* | 614 | 67.2% | 1250 | 65.6% |
| *Alchemilla_colura* | 582 | 63.7% | 1170 | 61.4% |
| *Alchemilla_cryptantha* | 604 | 66.1% | 1234 | 64.7% |
| *Alchemilla_decumbens* | 485 | 53.1% | 902 | 47.3% |
| *Alchemilla_ellenbeckii* | 615 | 67.3% | 1244 | 65.3% |
| *Alchemilla_elongata* | 605 | 66.2% | 1234 | 64.7% |
| *Alchemilla_fischeri* | 590 | 64.6% | 1224 | 64.2% |
| *Alchemilla_fissa* | 252 | 27.6% | 408 | 21.4% |
| *Alchemilla_flabellata* | 431 | 47.2% | 812 | 42.6% |
| *Alchemilla_glabra* | 510 | 55.8% | 927 | 48.6% |
| *Alchemilla_haumanii* | 570 | 62.4% | 1182 | 62.0% |
| *Alchemilla_heptagona* | 503 | 55.0% | 930 | 48.8% |
| *Alchemilla_hildebrandtii* | 481 | 52.6% | 965 | 50.6% |
| *Alchemilla_indivisa* | 487 | 53.3% | 896 | 47.0% |
| *Alchemilla_japonica* | 518 | 56.7% | 949 | 49.8% |
| *Alchemilla_johnstonii* | 583 | 63.8% | 1207 | 63.3% |
| *Alchemilla_kiwuensis* | 554 | 60.6% | 1159 | 60.8% |
| *Alchemilla_lapeyrousii* | 441 | 48.2% | 850 | 44.6% |
| *Alchemilla_microbetula* | 607 | 66.4% | 1239 | 65.0% |
| *Alchemilla_mollis* | 589 | 64.4% | 1060 | 55.6% |
| *Alchemilla_pentaphyllea* | 516 | 56.5% | 933 | 49.0% |
| *Alchemilla_plicata* | 398 | 43.5% | 745 | 39.1% |
| *Alchemilla_roccatii* | 580 | 63.5% | 1206 | 63.3% |
| *Alchemilla_rutenbergii* | 573 | 62.7% | 1163 | 61.0% |
| *Alchemilla_schizophylla* | 536 | 58.6% | 1069 | 56.1% |
| *Alchemilla_splendens* | 431 | 47.2% | 802 | 42.1% |
| *Alchemilla_stuhlmannii* | 467 | 51.1% | 964 | 50.6% |
| *Alchemilla_subnivalis* | 563 | 61.6% | 1154 | 60.5% |
| *Alchemilla_subsericea* | 496 | 54.3% | 932 | 48.9% |
| *Alchemilla_triphylla* | 477 | 52.2% | 967 | 50.7% |
| *Alchemilla_woodii* | 497 | 54.4% | 1004 | 52.7% |
| *Alchemilla_xanthochlora* | 480 | 52.5% | 883 | 46.3% |
| *Aphanes_arvensis* | 453 | 49.6% | 639 | 33.5% |
| *Aphanes_australis* | 444 | 48.6% | 619 | 32.5% |
| *Aphanes_cotopaxiensis* | 424 | 46.4% | 598 | 31.4% |
| *Aphanes_microcarpa* | 428 | 46.8% | 599 | 31.4% |
| *Aphanes_occidentalis* | 441 | 48.2% | 618 | 32.4% |
| *Aphanes_sp "boliviana"* | 433 | 47.4% | 608 | 31.9% |
| *Aphanes_sp "patagonica"* | 450 | 49.2% | 630 | 33.1% |

| | | | | |
|---|---|---|---|---|
| *Chamaerhodos_erecta* | 737 | 80.6% | 759 | 39.8% |
| *Comarum_palustre* | 773 | 84.6% | 830 | 43.5% |
| *Dasiphora_fruticosa* | 748 | 81.8% | 773 | 40.6% |
| *Drymocallis_arguta* | 756 | 82.7% | 777 | 40.8% |
| *Drymocallis_glandulosa* | 747 | 81.7% | 764 | 40.1% |
| *Farinopsis_salesoviana* | 849 | 92.9% | 933 | 49.0% |
| *Fragaria_vesca* | 715 | 78.2% | 740 | 38.8% |
| *Lachemilla_andina* | 720 | 78.8% | 1309 | 68.7% |
| *Lachemilla_angustata* | 653 | 71.4% | 1180 | 61.9% |
| *Lachemilla_aphanoides* | 721 | 78.9% | 1295 | 67.9% |
| *Lachemilla_barbata* | 726 | 79.4% | 1285 | 67.4% |
| *Lachemilla_diplophylla* | 731 | 80.0% | 1301 | 68.3% |
| *Lachemilla_equisetiformis* | 542 | 59.3% | 976 | 51.2% |
| *Lachemilla_erodiifolia* | 699 | 76.5% | 1261 | 66.2% |
| *Lachemilla_galioides* | 725 | 79.3% | 1346 | 70.6% |
| *Lachemilla_hirta* | 722 | 79.0% | 1294 | 67.9% |
| *Lachemilla_hispidula* | 711 | 77.8% | 1307 | 68.6% |
| *Lachemilla_jamesonii* | 734 | 80.3% | 1330 | 69.8% |
| *Lachemilla_llanganatensis* | 592 | 64.8% | 1087 | 57.0% |
| *Lachemilla_mandoniana* | 694 | 75.9% | 1225 | 64.3% |
| *Lachemilla_moritziana* | 549 | 60.1% | 1014 | 53.2% |
| *Lachemilla_nivalis* | 702 | 76.8% | 1283 | 67.3% |
| *Lachemilla_orbiculata* | 695 | 76.0% | 1240 | 65.1% |
| *Lachemilla_pectinata* | 736 | 80.5% | 1335 | 70.0% |
| *Lachemilla_pinnata* | 720 | 78.8% | 1281 | 67.2% |
| *Lachemilla_polylepis* | 727 | 79.5% | 1324 | 69.5% |
| *Lachemilla_pringlei* | 604 | 66.1% | 1073 | 56.3% |
| *Lachemilla_procumbens* | 712 | 77.9% | 1270 | 66.6% |
| *Lachemilla_ramosissima* | 646 | 70.7% | 1154 | 60.5% |
| *Lachemilla_sibbaldiifolia* | 598 | 65.4% | 1072 | 56.2% |
| *Lachemilla_standleyi* | 662 | 72.4% | 1195 | 62.7% |
| *Lachemilla_tanacetifolia* | 730 | 79.9% | 1319 | 69.2% |
| *Lachemilla_velutina* | 629 | 68.8% | 1097 | 57.6% |
| *Lachemilla_verticillata* | 734 | 80.3% | 1347 | 70.7% |
| *Lachemilla_vulcanica* | 732 | 80.1% | 1335 | 70.0% |
| *Potaninia_mongolica* | 737 | 80.6% | 761 | 39.9% |
| *Sibbaldia_procumbens* | 703 | 76.9% | 744 | 39.0% |
| *Sibbaldianthe_adpressa* | 717 | 78.4% | 752 | 39.5% |
| *Sibbaldianthe_bifurca* | 725 | 79.3% | 762 | 40.0% |
| *Potentilla_indica* | 849 | 92.9% | N/A | N/A |
| *Rosa_woodsii* | 913 | 99.9% | N/A | N/A |
| *Sanguisorba_menziesii* | 860 | 94.1% | N/A | N/A |
